# Activation of the medial entorhinal cortex drives memory-guided navigation behavior

**DOI:** 10.64898/2025.12.24.696388

**Authors:** Yan Ma, Taylor J. Malone, Nai-Wen Tien, June Hoan Kim, Maxym V. Myroshnychenko, Jean Tyan, Daniel Yusufi, Farid Shahid, David A. Kupferschmidt, Yi Gu

## Abstract

The cognitive map in the medial entorhinal cortex (MEC) is essential for spatial memory and exhibits experience-dependent changes. Yet, it remains unclear whether MEC activation is sufficient to bias the network toward memory-guided navigation, precluding a causal link between the MEC and spatial memory. To address this gap, we examined and optogenetically manipulated MEC activity as mice navigated a virtual track for a water reward based on memory. During learning, spatial activity in the pre-reward region showed the highest consistency along the track and closely paralleled improvements in reward-predictive behaviors. This elevated pre-reward consistency was widespread across MEC neurons, rather than confined to a particular cell type, indicating a network-level representation supporting reward prediction. Strikingly, optogenetic activation of MEC at non-reward locations biased reward-predictive behaviors toward stimulation sites, but only when stimulation patterns were spatially consistent. Consistent stimulation also induced anticipatory reward-seeking before stimulation onset and this behavior gradually developed with repeated stimulation, reflecting memory formation for stimulation locations. Moreover, in animals that failed to learn the track well, consistent pre-reward activation, alone or combined with landmark activation, significantly enhanced their reward prediction. Thus, spatially consistent MEC activation underlies memory-guided reward prediction, supporting a causal role for the MEC in spatial memory.

## Introduction

The hippocampal-entorhinal cognitive map is fundamental to memory-guided spatial navigation, a role that has been recognized for decades^1,2^. Among its components, the medial entorhinal cortex (MEC) is particularly crucial: dysfunction in this region, whether in animal models, humans, or patients with Alzheimer’s disease, is invariably associated with pronounced spatial memory deficits^3,4^. The MEC hosts diverse cell types, including grid cells^5^, head-direction cells^6^, speed cells^7^, border cells^8^, and object vector cells^9^/cue cells^10^, which together generate a cognitive map encoding position, orientation, speed, environmental boundaries, and landmarks during navigation^11^.

Importantly, the MEC cognitive map is highly dynamic, adapting with spatial experience. In novel environments, MEC spatial activity consistency increases as animals learn, a process tightly linked to successful memory formation; conversely, disruptions in this consistency impair spatial memory^12^. Grid cell activity, characterized by firing at multiple locations arranged at the vertices of a triangular lattice, undergoes experience-dependent changes in stability^5,12^, spacing between spatial fields^12–14^, regularity^13^ and symmetry^15^ of the firing pattern, and the continuity of spatial representation^16^. Environmental changes also induce experience-dependent directional biases in the head-direction cells^17^.

Collectively, these findings establish that MEC activity is necessary for spatial memory and tracks the learning of environmental features, underscoring the role of the MEC cognitive map in mnemonic spatial encoding. Supporting this, studies have shown that during reward-guided spatial learning in virtual reality (VR), MEC neurons increase their spatial fields near the reward site only after successful spatial learning, and the largest difference in spatial activity consistency between successful and unsuccessful learning exists in the pre-reward region^12^. In open arenas, firing patterns of grid and non-grid cells are distorted by shifting spatial fields toward reward locations^18,19^, leading to more precise encoding of these sites^18^. However, it remains unclear whether enhanced MEC activity near reward locations merely reflects the presence of reward, or whether it actively contributes to memory-guided reward prediction. More broadly, whether MEC activation is sufficient to engage the circuitry supporting spatial memory remains unresolved. Direct causal evidence linking the MEC cognitive map to spatial memory has therefore been lacking. Addressing this gap is critical both for defining MEC function and for exploring strategies to enhance memory through MEC modulation in clinical setting.

Here, we examined and optogenetically manipulated MEC neural activity during a VR navigation task, in which mice pursued water rewards based on memory. We demonstrate that the increase in spatial activity consistency of the MEC in the pre-reward region was most closely associated with improved reward-predictive behaviors during spatial learning. MEC cells, regardless of their activity features, generally exhibited the highest activity consistency in the pre-reward region compared to other track locations. These results indicate a potential role of consistent MEC activity in driving reward-prediction. Strikingly, optogenetically inducing spatially consistent MEC activation at non-rewarded locations was sufficient to elicit reward-predictive behaviors at those sites. Spatially consistent MEC activation before the reward, alone or combined with landmark activation, enhanced reward-predictive behaviors in mice that learned the track poorly. These results demonstrate the sufficiency of MEC activation for memory-guided reward prediction, establishing a causal relationship between the MEC cognitive map and spatial memory.

## Results

### MEC spatial activity consistency before reward best matches reward-predictive behaviors

We first investigated the correlation between MEC activity and memory-guided navigation using our published dataset, where MEC calcium activity was measured via cellular-resolution two-photon imaging as mice learned a novel virtual linear track over 10 days^12^ (**Fig. 1a**). The 4-meter track featured symmetrically arranged visual landmarks and a water reward at a fixed location. Water-restricted GP5.3 mice, stably expressing the fluorescent calcium indicator GCaMP6f in MEC layer 2 excitatory neurons^20–22^ (**Fig. 1b**), pursued the reward by unidirectionally navigating the track for multiple runs (**Fig. 1c**). During successful learning, the mice that well learned the track (“good performers”, see “**Methods**” for behavioral classification^12^) showed increased predictive slowing and licking specifically before the reward (**Fig. 1d**), indicating that the mice were expecting the reward and distinguishing the reward location from other areas based on the established spatial memory. Reward-predictive slowing and licking were quantified as the percentiles of slowing (i.e., speed decrease) and number of licks before the reward compared to the rest of the track, respectively. Over 10 days, the behaviors gradually improved (**Fig. 1e**), as described previously^12^. We previously reported that spatial consistency of MEC activity is necessary for spatial memory and varies across track locations^12^. To investigate how consistency at different locations contributes to spatial memory, we examined the association between the consistency along the track and reward-predictive behaviors of these mice. Consistency was calculated per cell as run-by-run spatial activity correlation within 25-cm rolling windows along the track and combined across all cells each day (**Fig. 1f, S1, and S2a**). To validate that run-by-run spatial activity consistency captures genuine spatially specific tuning rather than correlated activity fluctuations across runs, we also measured population coherence by calculating population-vector correlations across runs and track locations (**Fig. S3a**). Population coherence across runs was higher between the same track locations than between different locations, indicating that population activity exhibits location-specific spatial tuning across runs rather than co-fluctuating uniformly across locations. (**Fig. S3b**). This location-specific coherence was strongly correlated with run-by-run activity consistency during learning and at individual spatial bins (**Fig. S3c-h**), indicating that per cell run-by-run consistency can successfully capture population-level location-specific representations.

**Figure 1.**
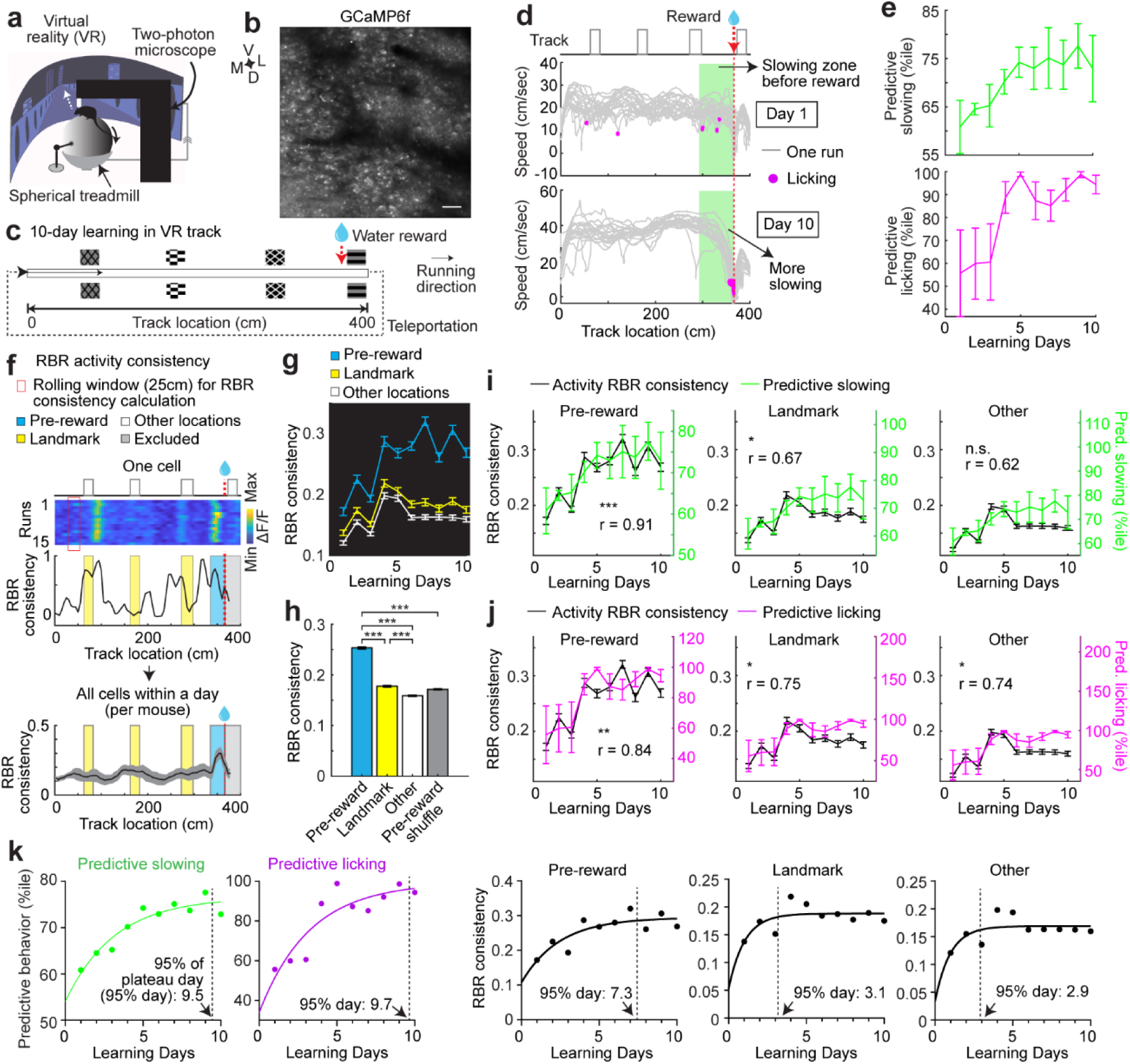
MEC spatial activity consistency before reward best matches reward-predictive behaviors. **a.** Virtual reality navigation and two-photon imaging of mouse MEC, adapted from^12^. **b.** An example two-photon FOV. Scale bar: 50 µm. **c.** 4m track, adapted from^12^. **d.** Examples of run-by-run (RBR) speed and licking of a mouse on days 1 and 10, showing increased reward-predictive slowing and licking (in green zone). Note that positive and negative speeds indicate that the mice move forward and backward along the track, respectively. The first 20 runs were plotted on each day. **e.** Reward-predictive slowing (top) and licking (bottom) across learning, from 4 mice **f.** Top: RBR calcium activity of one cell with its RBR activity consistency. Bottom: the consistency of all cells averaged per mouse within a day (day 8, N = 4 mice pooled from 7 FOVs with 194 ± 41 cells per mouse). The consistency of a cell was calculated by correlating its activity across runs in 25-cm rolling windows along the track, grouped in different types: pre-reward (blue), at landmarks (yellow), and in other locations (white). The region after reward was excluded (gray). To ensure complete coverage of all track locations and comparable calculations across FOVs, the first run, which typically had incomplete track coverage due to the delayed imaging initiation relative to the onset of VR navigation, was excluded from each session and only the next 15 runs were used. **g.** Comparison of the three consistency types across 10 days. The data on individual days contained 901 ± 41 cells pooled from 7 FOVs in 4 mice with 15 ± 0 runs per day. **h.** Comparison of the three consistency types and shuffled consistency at pre-reward the region (shuffling the consistency of each cell with respect to track location), pooling all values across 10 days (all cells from 7 FOVs of 4 mice). Paired Student’s t-test. p values: pre-reward vs landmark: 1.28×10^−222^; pre-reward vs other locations: 0; landmark vs other locations: 6.14×10^−54^; pre-reward vs shuffle: 0. **i.** Left to right: pre-reward, landmark, and other consistencies across days (left y axes), plotted with predictive slowing (right y axes) to show their correlation (r). “*” indicates correlation significance. **j.** Similar to **i** but for the three consistency types and predictive licking. **k.** Left to right: mean predictive slowing, predictive licking, pre-reward, landmark, and other consistencies across 10 days were fit with one-phase exponential association curves. *p ≤ 0.05, **p ≤ 0.01, ***p ≤ 0.001. Error bars represent mean ± SEM.

Spatial consistency was then calculated for three categories: immediately before the reward (pre-reward consistency), at landmarks (landmark consistency), and at other locations (other consistency, excluding locations between the reward and track end due to reward consumption effects). Pre-reward and landmark consistencies were higher than other consistency, with pre-reward consistency being the highest (**Fig. 1g, 1h, S4a, and S4e**), as previously reported^12^. Pre-reward consistency also exceeded shuffled controls generated by global spatial shuffling across the full track, with subsequent extraction of the pre-reward region (**Fig. 1h and S4e**).

While all three consistency types improved during learning, pre-reward consistency dynamics best matched the increase in predictive slowing (**Fig. 1i**) and licking (**Fig. 1j**), showing higher correlations. We further fitted the behaviors and activity consistencies with exponential curves and calculated the days they reached 95% of their plateaus (95% day). The 95% days for predictive slowing (9.5 days) and licking (9.7 days) were similar, and more comparable with pre-reward consistency (7.3 days) than with landmark (3.1 days) or other consistencies (2.9 days) (**Fig. 1k**). Thus, pre-reward consistency better tracks the improvements in reward-prediction.

To better quantify the relationship between spatial consistency and mouse behavior, we calculated the mutual information (MI) between spatial consistency and mouse velocity or lick probability. Mouse velocity and lick probability were calculated within 25-cm rolling windows along the track, as was performed above for activity consistency, for all pre-reward locations (**Fig. S3i**). MI was calculated between activity consistency and behavioral variables for each day and was significantly higher than when mouse behavior was shuffled across track locations (**Fig. S3j-m**), indicating that spatial consistency can explain a significant portion of velocity and licking variance.

We further used Generalized Linear Mixed Models (GLMMs) to directly quantify the ability of spatial consistency to predict behavior. Individual observations were the activity consistency and behavior at individual spatial bins averaged within each day and combined across all days and mice. Because lick probability was non-zero only within a narrow pre-reward region on most days (**Fig. S3i**), which limited reliable model estimation (data not shown), only velocity was effectively predicted. Spatial consistency, acceleration, distance to the reward location, and lick probability were included as fixed-effect predictors, while mouse identity and day number were used as random effects (**Fig. S3n**). Spatial consistency was a significant contributor to predictive ability and had the largest contribution to variance explained (15%) among the fixed-effects predictors (**Fig. S3o and S3p)**, indicating a meaningful predictive ability. Together, these results demonstrate that spatial activity consistency can predict learning-related behavioral variables, particularly mouse velocity.

Finally, using a previously published dataset in which calcium dynamics were measured during learning of a track with a reward located in the middle of the track^23^, we found that activity consistency in pre-reward regions, but not in post-reward regions, gradually increased during learning, and the consistency remained higher in pre-reward than post-reward regions after learning (**Fig. S5)**. This selective increase in pre-reward activity consistency supports a role of such activity in reward prediction, rather than reflecting a generalized reward-related stabilization of activity, which would be expected to produce similar changes in both pre- and post-reward activity consistency.

Aligned with the above observations, we previously showed that pre-reward consistency best distinguishes good performers from poor performers, the latter displaying weak reward-predictive behaviors and low activity consistency, particularly in the pre-reward zone, throughout learning^12^. Together, the robust experience-dependent increase in pre-reward MEC activity consistency and its strong association with reward-predictive behavior suggest that consistent pre-reward activity may play an important role in driving memory-guided reward prediction.

### High spatial activity consistency before reward is a general feature of MEC cells

If consistent MEC activity at the pre-reward location supports robust reward-predictive behavior, we hypothesized that this activity reflects a network-wide phenomenon, represented by the majority of cells rather than a small subset. To test this, we assessed activity consistency across different functional cell types in MEC layer 2. We examined grid cells^21,24^, which are known to shift their spatial fields toward reward locations during reward-guided navigation in open arenas^18,19^ (**Fig. 2a and S1a**). We also investigated cue cells, which are abundant during visual-landmark-enriched VR navigation and are specifically active near individual landmarks^10^ (**Fig. 2b and S1b**). The remaining cells were categorized as “other cells” (**Fig. 2c and S1c**). Cells were classified on a per-day basis, yielding 43.8 ± 1.5% grid cells, 15.6 ± 0.6% cue cells, and 40.7 ± 1.4% other cells.

**Figure 2.**
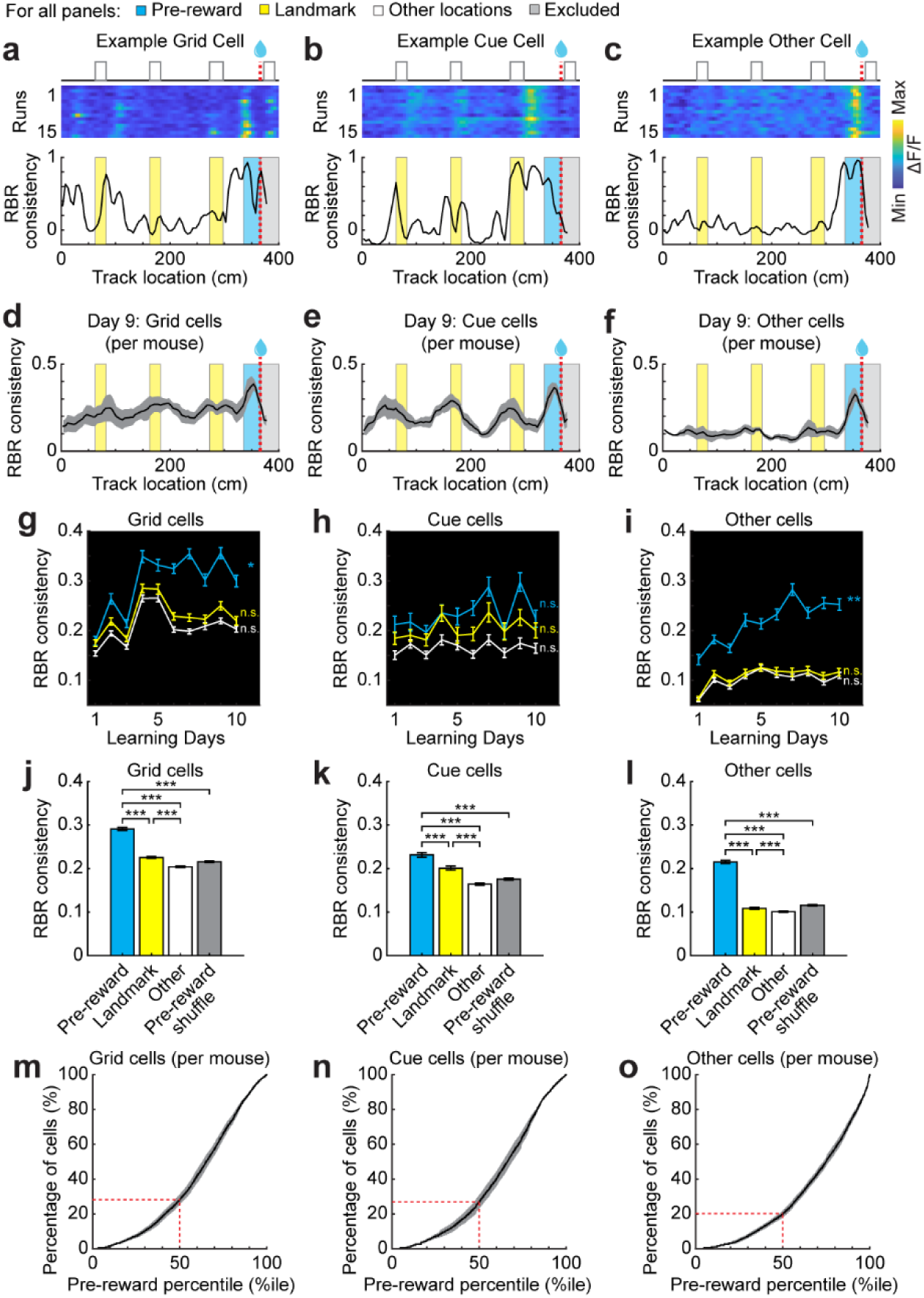
High spatial activity consistency before reward is present across functional cell types in the MEC. For all panels: blue, yellow, and white represent RBR activity consistency in pre-reward, landmarks, and other locations, respectively. The region after reward (gray) was excluded. **a-c.** Top: RBR calcium activity of an example grid cell (**a**), cue cell (**b**), and other cell (**c**). Bottom: RBR activity consistency of each example cell as a function of track location. N=15 runs. **d-f.** Mean consistency of grid cells (**d**), cue cells (**e**), and other cells (**f**) within day 9 averaged across mice. N=4 mice from 7 FOVs (cells per mouse: **d.** 92 ± 35. **e.** 33 ± 6. **f.** 86 ± 9). 15 runs per day. **g-i**. Comparison of three consistency types, consistencies at reward, landmarks, and other locations, across 10 days for grid cells (**g**), cue cells (**h**), and other cells (**i**). The data on individual days contained 421 ± 20 cells (**g**), 140 ± 10 cells (**h**), 340 ± 14 cells (**i**) pooled from 7 FOVs in 4 mice with 15 ± 0 runs per day. Pearson correlation was calculated between mean consistency and learning day. Significant correlations and p values: **g**. pre-reward: r=0.66, p=0.039; **i**. pre-reward: r=0.87, p=0.0012. **j-l.** Comparison of the three consistency types and shuffled pre-reward consistency by pooling all values across 10 days for grid cells (**j**), cue cells (**k**), and other cells (**l**). Paired Student’s t-test. p values: **j**. pre-reward vs landmark: 4.57×10^−74^; pre-reward vs other locations: 2.78×10^−156^; landmark vs other locations: 3.62×10^−29^; pre-reward vs shuffle: 4.54×10^−146^. **k**. pre-reward vs landmark: 7.48×10^−7^; pre-reward vs other locations: 2.71×10^−37^; landmark vs other locations: 1.96×10^−23^; pre-reward vs shuffle: 2.77×10^−32^. **l**. pre-reward vs landmark: 2.88×10^−192^; pre-reward vs other locations: 1.37×10^−247^; landmark vs other locations: 3.66×10^−7^; pre-reward vs shuffle: 1.80×10^−239^. **m-o.** Percentile of mean pre-reward consistency among all remaining bins (including landmark and other locations) for grid cells (**m**), cue cells (**n**), and other cells (**o**). Dashed red lines mark the percentage of cells below 50^th^ percentile. This percentage is below 30% for all cell types, therefore, the percentages of cells above 50^th^ percentile were more than 70%. N = 4 mice (pooled from 7 FOV and 10 days). *p ≤ 0.05, **p ≤ 0.01, ***p ≤ 0.001. Error bars represent mean ± SEM.

We observed that all cell populations reliably exhibited the highest run-by-run activity consistency at the pre-reward location (**Fig. 2d-f and S2b-d**), and this trend persisted across learning days (**Fig. 2g-I, S4b-d, and S4f-h**). Only pre-reward consistency gradually increased in both grid and other cells (**Fig. 2g, 2i, S4b, and S4d**). High pre-reward consistency was followed by lower consistency at landmarks and the lowest consistency at other locations (**Fig. 2j-l and S4f-h**) and pre-reward consistency was again higher than shuffled consistency (**Fig. 2j-l and S4f-h**).

At the individual cell level within each day and imaging field of view (FOV), we quantified the proportion of cells whose pre-reward activity consistency ranked at specific percentiles among those across the rest of the track. Across all cell types, more than 70% of cells exceeded the 50^th^ percentile (**Fig. 2m-o**), indicating that most cells showed relatively higher activity consistency at the pre-reward site than elsewhere along the track.

To further demonstrate synchronized pre-reward activity across MEC cells, we quantified population synchrony as the pairwise activity correlation between neurons within 25-cm rolling windows along the track. Like activity consistency, population synchrony generally increased during learning across the full track and in pre-reward bins, although only significantly for pre-reward bins (**Fig. S6a and S6b**). The increase in pre-reward synchrony was particularly apparent when comparing population synchrony as a function of track location in early (day 1) versus late (day 7-10) learning (**Fig. S6c and S6d**), suggesting that the overall MEC population progressively forms a coherent pre-reward representation during learning.

In summary, the high spatial activity consistency in the pre-reward location is a general feature of MEC cells. This network-wide feature aligns with prior findings that both grid and non-grid populations in the MEC shift their spatial fields toward learned reward sites during open-field navigation^18^. Together, these results suggest that MEC cells generate a collective signal that may support robust reward-predictive behaviors.

### Spatially consistent MEC activation at non-reward locations biases reward-predictive behaviors

To directly test if the consistent pre-reward activity in the MEC drives reward prediction, we asked whether consistent MEC activation at non-reward locations biases reward-predictive behaviors toward these locations. We attempted this by optogenetically activating the MEC using virally expressed channelrhodopsin-2 variant H134R (ChR2-GFP or ChR2-EYFP) predominately in excitatory neurons across a large dorsal-ventral extent of MEC layer 2 (ChR2 mice), with EGFP as an illumination control (EGFP mice)^12^ (**Fig. 3a and 3b; Fig. S7**). To reproduce the network-wide consistent activity observed earlier (**Fig. 2**), we utilized Lambda-B tapered optical fibers, which emit blue light from the entire tapered region to illuminate approximately 2 mm of the MEC along the dorsoventral axis^12,25^. In previous work, we confirmed robust ChR2 expression in excitatory neurons in MEC layer 2, enabling reliable light-evoked action potentials in the MEC *in vivo*^12^.

**Figure 3.**
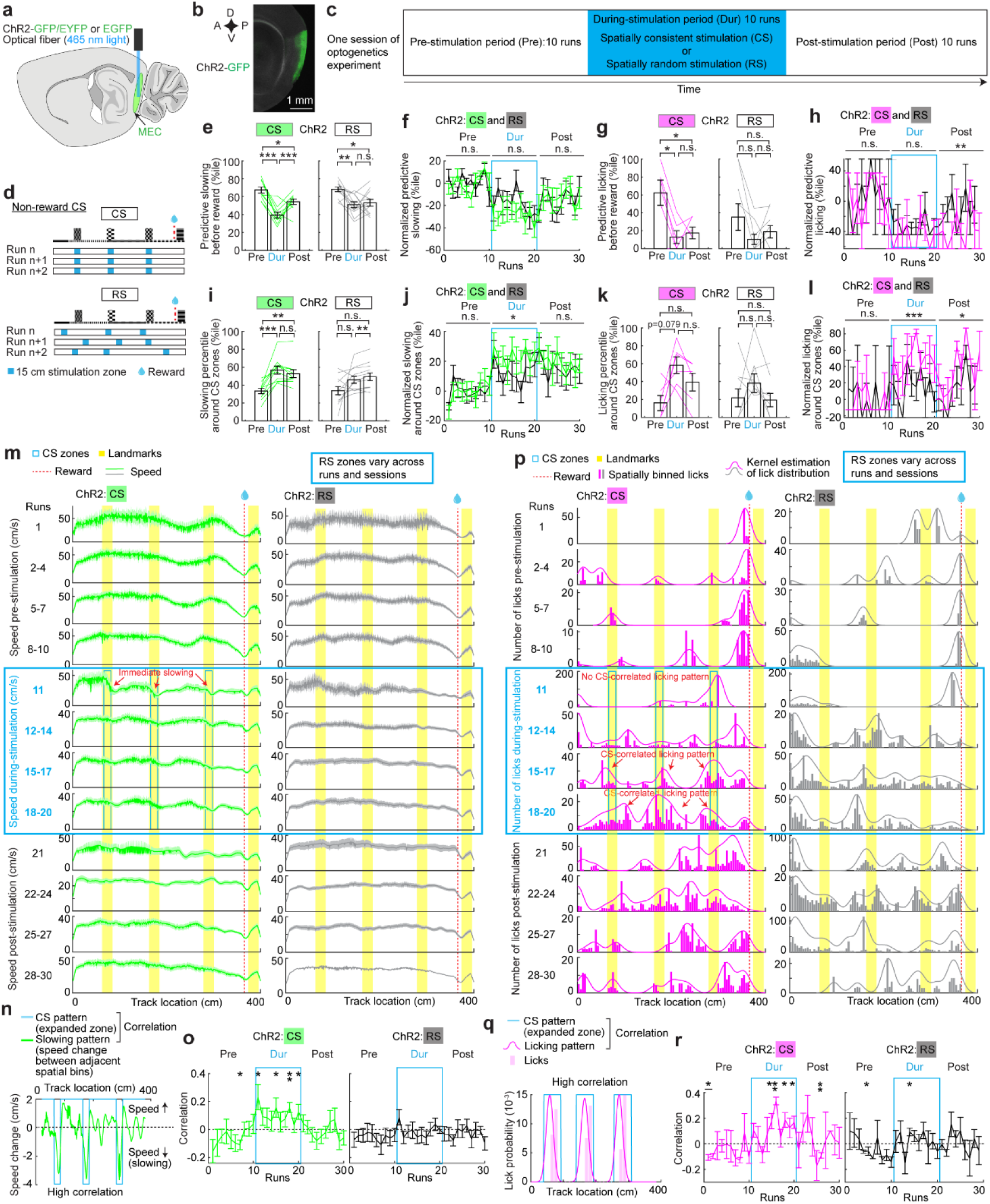
Spatially consistent optogenetic stimulation of the MEC at 15-cm non-reward locations biases reward-predictive behaviors. **a.** Schematic displaying the location of the optical fiber and blue LED light delivery to activate ChR2-GFP/EYFP- or EGFP-expressing neurons in the MEC, adapted from^12^. **b.** Epifluorescence image of a sagittal section showing ChR2-GFP expression in superficial layers of the MEC. D: dorsal; V: ventral; A: anterior; P: posterior. **c.** Schematic of an optogenetics session. **d.** Stimulation patterns for non-reward CS and RS. **e.** Predictive slowing of ChR2 mice in pre-, during- (Dur), and post-stimulation for non-reward CS and RS, calculated by averaging RBR values per session (green lines). Student’s paired t-test was used between different periods of the same session. P values were adjusted for multiple comparisons using Bonferroni–Holm method. The same method was used in similar plots. Significant p values for CS: Pre vs Dur: 3.02×10^−4^; Pre vs Post: 1.40×10^−2^; Dur vs Post: 2.61×10^−2^. Significant p values for RS: Pre vs Dur: 6.54×10^−3^; Pre vs Post: 2.49×10^−2^. **f.** RBR predictive slowing of ChR2 mice, CS vs RS. Data per session were normalized by subtracting the session’s mean pre-stimulation value. “n.s.” above horizontal lines indicates the p value comparing CS and RS during each period (two-way ANOVA). The same methods were applied to similar plots. **g.** Similar to **e** but for predictive licking. Due to the unreliability of licking behavior (i.e., mice often slowed down without licking), the values for runs without licking could not be calculated, resulting in invalid run-averaged mean value of a session within a certain period. Student’s paired t-test and multiple comparison correction were only performed on the sessions with valid mean values in both comparison periods. This method was applied to similar plots below. Significant p values for CS: Pre vs Dur: 1.69×10^−2^; Pre vs Post: 4.78×10^−2^. **h.** Similar to **f** but for RBR predictive licking. Due to invalid mean values for some sessions in pre-stimulation period, data per session were normalized by subtracting the mean value across all sessions in pre-stimulation period. Student’s t-test was used to compare CS and RS (“***” above horizontal line). This method was applied to similar plots below. Significant p values: CS vs RS Post: 9.89×10^−3^. **i.** Slowing percentile around stimulation zones of ChR2 mice, CS vs RS, similarly calculated as in **e** but for combined values in the three stimulation zones compared to other zones. Significant p values for CS: Pre vs Dur: 8.51×10^−4^; Pre vs Post: 6.12×10^−3^. Significant p values for RS: Dur vs Post: 1.62×10^−2^. **j.** RBR slowing percentile around stimulation zone of ChR2 mice, CS vs RS, similar to **f**. Note that during stimulation, CS values trend greater than RS. **k.** Licking percentile around stimulation zones of ChR2 mice, CS vs RS. It was similarly calculated as in **g** but for combined values in the three stimulation zones compared to other zones. **l.** RBR licking percentile around stimulation zone of ChR2 mice, CS vs RS, similar to **h**. **m.** Speed in pre-, during- and post-stimulation periods of ChR2 mice, CS vs RS. Except the first run in each period that shows the immediate effect of with and without stimulation, the speed was averaged across every three runs. **n.** Schematic for correlating CS pattern (expanded to compromise for speed variation) with slowing pattern of a run, reflected by speed change from one spatial bin to the next bin. Negative speed change corresponds to slowing. **o.** Correlation between CS and slowing patterns of ChR2 mice, CS vs RS. “*” indicates the comparison between the correlation in each run and zero (one-sample Student’s t-test). Significant p values from left to right: 1.17×10^−2^; 2.96×10^−2^; 4.80×10^−2^; 1.94×10^−3^; 4.16×10^−2^. **p.** Similar to **m** but for number of licks. Licks after reward delivery to the end of the track were removed. **q.** Schematic for correlating CS pattern (expanded to compromise for licking location variation) with licking pattern, calculated from averaged lick numbers per run. **r.** Correlation between CS and licking patterns of ChR2 mice, CS vs RS. “*” indicates the comparison between the correlation in each run and zero (one-sample Student’s t-test). Significant p values from left to right: 4.12×10^−2^; 4.37×10^−2^; 4.18×10^−2^; 7.71×10^−3^; 2.35×10^−2^; 2.51×10^−2^; 7.55×10^−3^; 3.45×10^−2^; 3.72×10^−2^. CS data were from four mice, 10 sessions. RS data were from four mice, 12 sessions. *p ≤ 0.05, **p ≤ 0.01, ***p ≤ 0.001. n.s. p > 0.05. Error bars represent mean ± SEM.

To assess optimal stimulation conditions, we first evaluated light-evoked neuronal responses in the MEC of anesthetized mice across 12 stimulation conditions. Stimulation trains lasted 1 second, with pulse durations of 2-20 ms (milliseconds) and frequencies of 10-40 Hz. Low-frequency components revealed progressively accommodating responses with repeated stimulation, with distinct decay profiles depending on stimulation parameters (**Fig. S8a**). Exponential fitting of peak amplitudes of the low-frequency components demonstrated that the 2 ms 10 Hz condition exhibited the longest decay time constant, indicating minimal attenuation across repeated stimuli (**Fig. S8b-c**). Based on this result, 2 ms 10 Hz stimulation was selected for further analyses. Analysis of the stimulus-response profile from high-frequency filtered traces revealed that neuronal activity increased transiently following light stimulation but rapidly returned to baseline levels within ∼20 ms, indicating a short-lived effect (**Fig. S8d-f**). These characterizations together guided our use of 2 ms 10 Hz stimulation in behaving mice, as it produced reliable and stimulation-locked neural response, without causing sustained or cumulative activation in the network. Notably, the 1-s stimulation duration used here approximated the average stimulation duration applied to mice engaged in the behavioral task (∼ 0.54 s when mice navigated through a 15-cm stimulation zone at an average speed of ∼ 28 cm/s) (**Fig. 4g**). Thus, similar transient effects would be expected under experimental conditions.

**Figure 4.**
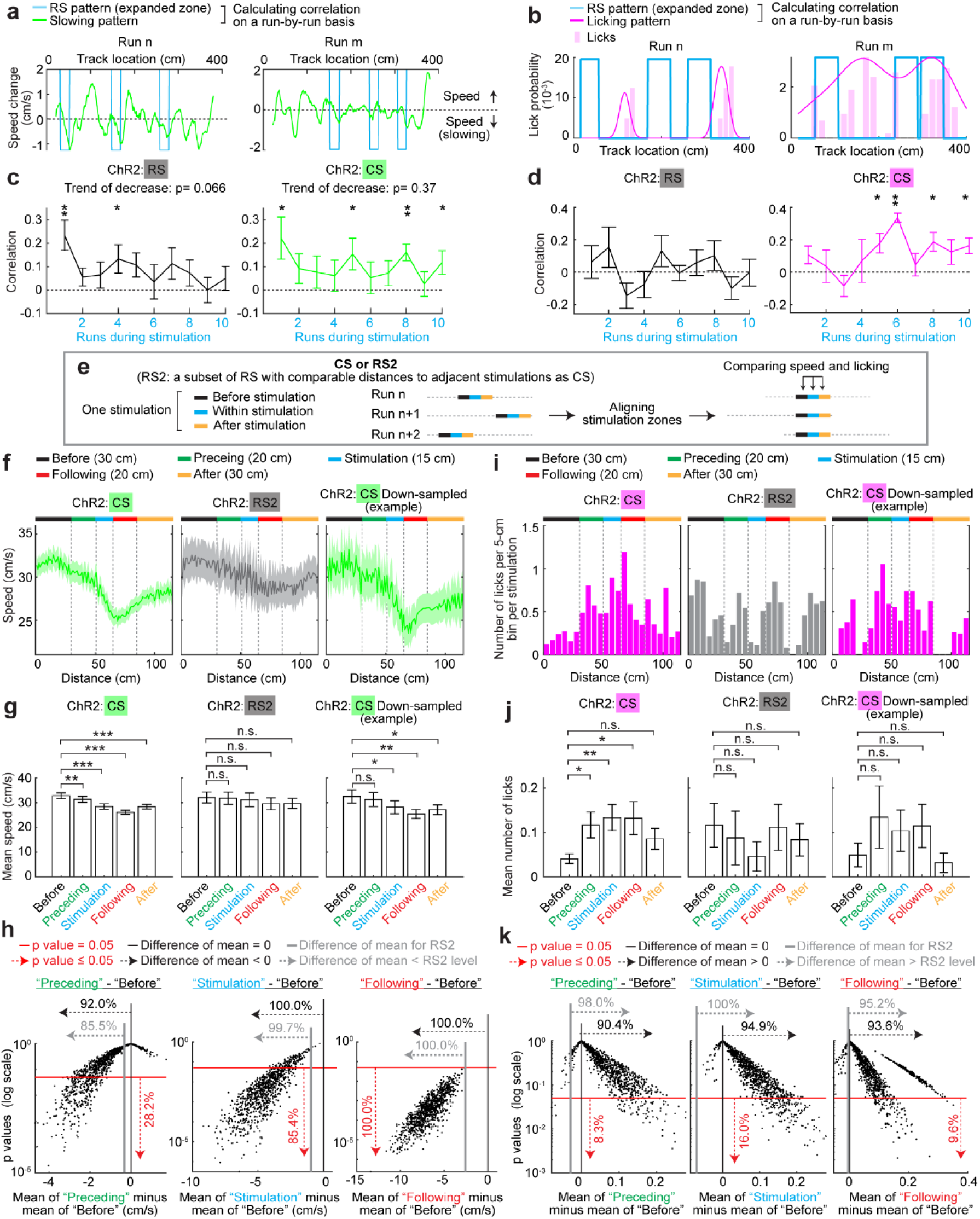
Spatially inconsistent MEC activation does not drive reliable reward-predictive behaviors. **a.** Schematic for correlating RS and speed change patterns in two runs, where the stimulation patterns (expanded to compromise for licking location variation) were different. **b.** Schematic for correlating RS and licking patterns in two runs, where the stimulation patterns (expanded to compromise for licking location variation) were different **c.** RBR correlations between stimulation and slowing patterns for RS and CS of ChR2 mice. The CS plot is highly similar to that for the stimulation period in Fig. 3o (left). Here, RBR correlations were computed between the slowing pattern and the actual stimulation location within each run, which deviated slightly from the predefined stimulation zone used in Fig. 3o due to minor computer timing offsets. As a result, the exact correlations and p values differ modestly between the two analyses, while the qualitative results and conclusions are unchanged. Significant p values from left to right (one-sample Student’s t-test): 4.20×10^−3^; 4.88×10^−2^; 3.57×10^−2^; 4.91×10^−2^; 1.37×10^−3^; 4.47×10^−2^. **d.** RBR correlations between stimulation and licking patterns for RS and CS of ChR2 mice. The CS plot is highly similar to that for the stimulation period in Fig. 3r (left), with slightly different correlations and p values but identical conclusions as explained in **c**. Significant p values from left to right (one-sample Student’s t-test): 4.09×10^−2^; 7.16×10^−3^; 2.31×10^−2^; 2.86×10^−2^. **e.** Schematic for aligning stimulation zones of individual CSs and RS2s. RS2s included a subset of RS, which had large distances to adjacent stimulations, comparable to CS. **f.** Speeds of ChR2 mice within five track zones (in 1-cm bins): before-stimulation zone (“before”, 30 cm), preceding zone (“preceding”, 20 cm), stimulation zone (“stimulation”, 15 cm), following zone (“following”, 20 cm), and after-stimulation zone (“after”, 30 cm). From left to right: all CS, all RS2, an example of down-sampled CS matching the stimulation number of RS2. **g.** Mean speeds within the five zones in **f**. The zones of the same stimulation were compared by paired Student’s t-test. Significant p values: CS before to: preceding: 3.98×10^−3^; stimulation: 7.47×10^−10^; following: 3.98×10^−3^; after: 1.34×10^−7^. Down-sampled CS before to: stimulation: 1.79×10^−3^; following: 2.33×10^−4^; after: 1.25×10^−2^. **h.** Left: speed comparison in “preceding” versus “before-stimulation” zones of 1000 down-sampled CS datasets. A slowing trend was observed in 92.0% of datasets, reflected by lower mean speed in “preceding” zone compared to “before-stimulation” zone (black vertical line indicates zero). 85.5% of datasets exhibited greater speed decrease (mean speed in “preceding” zone minus mean speed in “before-stimulation zone”) than RS2 (gray vertical line). 28.2% of datasets showed significant speed differences between the two zones (p ≤ 0.05, red horizontal line). Middle and right: similar to left but for “stimulation” versus “before stimulation” zones, and “following” versus “before stimulation” zones, respectively. **i.** Number of licks per stimulation, grouped in 5-cm bins in the same five zones in **f**. From left to right: all CS, all RS2, an example of down-sampled CS matching RS2 stimulation counts. **j.** Mean number of licks within the five zones in **i**. The zones of the same stimulation were compared by paired Student’s t-test. Significant p values: CS before to: preceding: 1.28×10^−2^; stimulation: 1.92×10^−3^; following: 1.57×10^−3^. **k.** Left: lick number comparison in “preceding” versus “before-stimulation” zones of 1000 down-sampled CS datasets. 90.4% of datasets showed increased licking in “preceding” zone, reflected by higher mean lick numbers in “preceding” zone compared to “before-stimulation” zone (black vertical line indicates zero). The differences between mean lick numbers in the two zones in 98% of datasets were above that of RS2 (gray vertical solid line). 8.3% of datasets showed significant lick number differences between the two zones (p ≤ 0.05, red horizontal line). Middle and right: similar to left but for “stimulation” versus “before stimulation” zones, and “following” versus “before stimulation” zones, respectively. CS data were from four mice, 10 sessions. RS data were from four mice, 12 sessions. *p ≤ 0.05, **p ≤ 0.01, ***p ≤ 0.001. n.s. p > 0.05. Error bars represent mean ± SEM.

The mice were trained to navigate the same 4-meter track (**Fig. 1c**), performing at least 30 runs per session. Optogenetic stimulation was applied during the middle 10 runs to compare reward-predictive behaviors across pre-, during-, and post-stimulation periods (**Fig. 3c**). Consistent run-by-run stimulation (CS) was applied at three 15-cm non-reward locations near landmarks (non-reward CS). Non-reward CS was compared with random stimulation (RS) within three 15-cm blocks randomized along the track on a run-by-run basis (**Fig. 3d; Fig. S9** for run-by-run stimulation patterns).

We observed that while mice initially displayed high reward-predictive slowing or licking in the pre-stimulation period, both CS and RS significantly disrupted these behaviors by reducing their percentiles compared to pre-stimulation by session (**Fig. 3e and 3g**). CS and RS also had similar effects across runs (**Fig. 3f and 3h**). However, the percentiles of slowing (**Fig. 3i and 3j**) and licking (**Fig. 3k and 3l**) around CS zones exhibited more robust trends of increased in CS than in RS, both with comparable pre-stimulation levels. These effects were absent in EGFP mice (**Fig. S10a-h**). Thus, while both CS and RS disrupt reward-predictive behaviors, only CS specifically biases the behaviors toward CS locations.

We next investigated run-by-run dynamics of speed and licking during stimulation. By averaging speed of all CS sessions per run, we observed that non-reward CS induced slowing around the three stimulation zones from the first run, persisting through all 10 stimulation runs (**Fig. 3m and Fig. S11a**). This was quantified as the correlation between CS pattern and the pattern of speed change (**Fig. 3n**), which confirmed an immediate and largely sustained CS-linked slowing during stimulation (**Fig. 3o**). Interestingly, while pre-reward licking was immediately disrupted by CS from the first run, licking near CS zones was only seen in later runs (**Fig. 3p and S11b**), as reflected by the gradually developed correlation between licking and CS patterns (**Fig. 3q and 3r**). The CS-zone-specific slowing and licking were absent during RS (**Fig. 3m-r and S11**) and in EGFP mice (**Fig. S10i-l and S12**). The immediate association between CS and slowing contrasted with the gradual association between CS and licking, suggesting that spatially consistent MEC activation across multiple runs is necessary to reliably bias reward-predictive behaviors, as evidenced by both slowing and licking. The CS-correlated slowing and licking quickly decayed after the stimulation was removed (**Fig. 3o and 3r**), suggesting that this CS protocol did not lead to long-term memory about the stimulation locations.

We further asked whether large spatial coverage of CS (i.e., 15 cm) is necessary to drive the above reward-predictive behaviors. We conducted CS in three 5-cm zones centered at the same track locations as non-reward CS (short-non-reward CS) (**Fig. S13a**). The experiments involved ChR2 and EGFP mice, both of which started with comparable reward-predictive slowing and licking as the mice receiving the 15-cm non-reward CS (“Pre” levels in **Fig. S13b and S13d** were close to those in **Fig. 3e and 3g**). The short-non-reward CS for ChR2 mice did not significantly reduce reward-predictive slowing (**Fig. S13b and S13c**) or licking (**Fig. S13d and S13e**), nor did it increase slowing (**Fig. S13f and S13g**) or licking (**Fig.S13h and S13i**) around CS locations. During stimulation of ChR2 mice, the run-by-run slowing pattern (**Fig. S13j and S13k**), but not licking pattern (**Fig. S13l and S13m**), weakly correlated with the stimulation pattern. No effect was observed in EGFP mice. Therefore, short-non-reward CS did not induce reliable reward-predictive behaviors around stimulation locations, indicating that larger spatial coverage of stimulation is necessary for such behaviors. We propose that the large-coverage stimulation recruits a broader and more coherent MEC ensemble than spatially restricted stimulation, and that such ensemble-level recruitment is required to drive reward-predictive behaviors. This interpretation is consistent with our finding that high spatial activity consistency before reward is a network-level feature, rather than a property of a small subset of cells.

To examine the specificity of MEC in driving reward-predictive behaviors, we tested the effects of optogenetic stimulation in visual cortex of mice expressing ChR2, with EGFP-expressing mice as controls (**Fig. S14a and S14b**). The same 15-cm non-reward CS zones were applied (**Fig. S14c**). Overall, stimulation in visual cortex did not produce behaviorally significant effects in ChR2 mice, either near the reward location (**Fig. S14d–g**) or within the stimulation zones (**Fig. S14h–k**). Although a transient slowing near stimulation zones was observed during the first few runs of stimulation, this effect diminished with continued stimulation (**Fig. S14l and S14m**). Moreover, stimulation did not induce selective accumulation of licking within the stimulation zones (**Fig. S14n and S14o**). Together, these results indicate that the robust reward-predictive behaviors observed around stimulation sites are specific to the MEC activation and are unlikely to arise from nonspecific effects of stimulating arbitrary cortical regions.

Thus, spatially consistent MEC activation across a large track zone drives reward-predictive behaviors in the surrounding area. The initial activation induces slowing, likely reflecting the mouse’s anticipation of receiving a reward. Continuous activation at the same location triggers both slowing and licking, indicating more proactive reward-seeking actions. These results suggest that reliable reward-prediction signals arise when MEC activity is consistently aligned with environmental features, enabling the translation of spatial representations into goal-directed actions.

### Spatially consistent, but not inconsistent, MEC activation drives reliable reward-predictive behaviors

If reliable reward prediction requires spatially consistent MEC activation, as suggested by the non-reward CS effect, we hypothesized individual non-reward RS would not trigger stimulation-associated reward-predictive behaviors. To test this, we compared the behavioral effects of non-reward RS and CS on both run-by-run and stimulation-by-stimulation levels.

We first analyzed correlations between RS patterns and the patterns of speed change or licking on a run-by-run basis (**Fig. 4a and 4b**). The first RS run showed a significantly positive correlation between speed change and stimulation, similar to CS, consistent with the immediate slowing effect of the stimulation. However, the correlations in subsequent RS runs were mostly insignificant and exhibited a strong trend of decrease toward zero, unlike the consistently positive correlations in CS across 10 runs (**Fig. 4c**). RS also failed to elicit stimulation-correlated licking, in contrast to the correlated licking developed in later CS runs (**Fig. 4d**). Therefore, RS did not evoke stimulation-associated reward predictive behaviors.

Importantly, CS and RS differed not only in spatial consistency, but also in the distance between adjacent stimulations within a run (“stim-distances”) (**Fig. S9**): CS stimulations were farther apart (minimum 99 cm), while RS stimulations could be much closer (minimum 35 cm). The small stim-distances in RS may hinder precise behavioral responses to each stimulation. To isolate the behavioral effects of CS and RS attributable to their differences in stimulation location consistency rather than stim-distance, we focused on a subset of RS (“RS2”) with stim-distances similar to CS (**Fig. S15**), aligning RS2 and CS locations and comparing speed and lick counts before, within, and after the stimulation zone (**Fig. 4e**).

CS induced slowing within both the stimulation zone and in the 20-cm zone immediately following stimulation offset (“following zone”) compared to the “before-stimulation zone” (20-50 cm before stimulation onset) (**Fig. 4f and 4g**, left). Slowing was also evident within 20 cm immediately preceding stimulation onset (“preceding zone”), indicating that mice anticipated the learned stimulation locations. Speed gradually increased in the “after-stimulation zone” (20-50 cm after stimulation offset). In contrast, RS2 did not produce significant slowing in any zone compared to the before-stimulation zone (**Fig. 4f and 4g**, middle). This difference was not explained by fewer RS2 stimulations, as down-sampling CS to match RS2 still produced significant and stimulation-associated slowing (**Fig. 4f and 4g**, right). Over 90% of 1000 down-sampled CS datasets showed slowing in preceding, stimulation, and following zones, and over 85% had greater slowing than RS2; all showed significant slowing in the following zone (**Fig. 4h**). These results indicate that individual RSs did not reliably induce stimulation-associated slowing.

We also compared lick numbers in the five zones. CS increased licking in the preceding, stimulation, and following zones compared to the before-stimulation zone (**Fig. 4i and 4j**, left). The increase in the preceding zone aligns with the preceding slowing (**Fig. 4f**), indicating memory of stimulation locations. These trends of increased licking were absent in RS2 (**Fig. 4i and 4j**, middle) but persisted after down-sampling CS to match RS2 (**Fig. 4i and 4j**, right). In over 90% of 1000 down-sampled CS datasets, there was an increased lick trend in the preceding, stimulation, and following zones, and more than 95% of the licking difference between these zones and the before-stimulation zone exceeded RS2 levels (**Fig. 4k**). Significant licking increases in down-sampled CS datasets were rare, likely because licking gradually developed in later stimulation runs (**Fig. 3r**). Thus, unlike CS, RS did not induce stimulation-associated licking.

In summary, RS’s failure to induce reward-predictive behaviors further highlights the necessity of spatially consistent MEC activation for effective reward prediction. Notably, during CS runs, mice exhibited anticipatory slowing and licking before stimulation onset, indicating their memory of stimulation locations.

### Memory about stimulation locations gradually develops during consistent stimulation

If the anticipatory slowing and licking observed before CS onset indeed reflected memory of stimulation locations, then these behaviors should emerge gradually during the stimulation period, but not in the pre- or post-stimulation periods. To test this, we analyzed running speed and licking in the before-stimulation and preceding zones across the three periods of non-reward CS sessions.

During the pre-stimulation period, mice ran faster in the preceding zone than in the before-stimulation zone (**Fig. 5a**, top, “Pre”), yielding a positive speed differences (preceding minus before-stimulation zone; **Fig. 5a**, bottom, “Pre”). Because non-reward CS zones were aligned with landmarks (**Fig. 3d**), this pattern likely reflects a natural tendency to increase speed when approaching landmarks. However, this trend was disrupted once stimulation began, as overall running speed decreased and speeds in the preceding zone gradually fell below those in the before-stimulation zone, producing significantly negative speed differences in later stimulation runs (**Fig. 5a**, “Dur”). After stimulation ended, overall speed initially remained low for several runs, but the trend of lower preceding speed disappeared, and speed gradually increased in later runs (**Fig. 5a**, “Post”).

**Figure 5.**
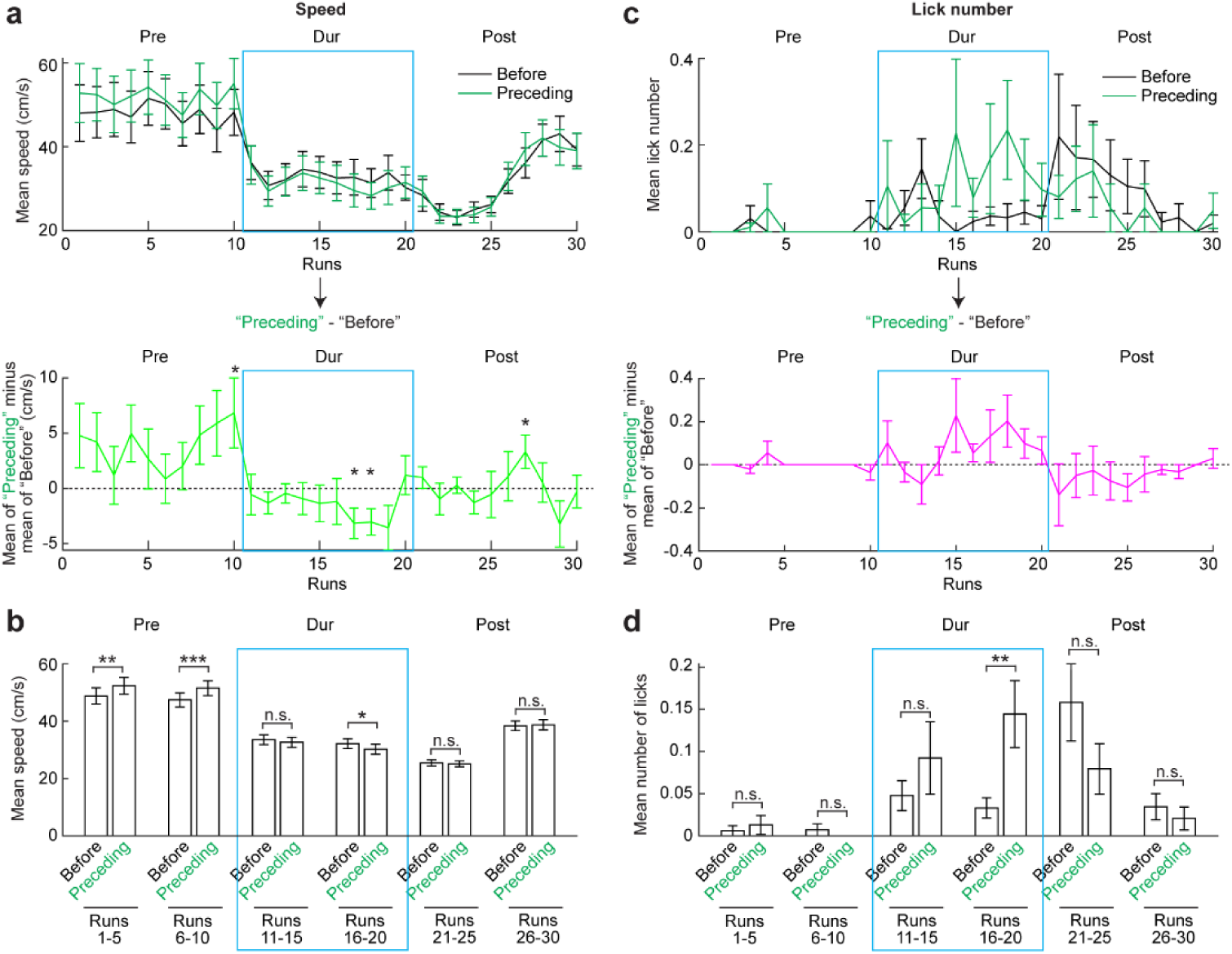
Anticipatory slowing and licking before stimulation onset gradually develops during consistent stimulation. **a.** Top: mean speed across sessions in individal runs in before-stimulation (black) and preceding (green) zones. Each data point represent the speeds of one stimulation (3 stimulations per run) in one run of one session. Bottom: mean speed in preceding zone minus that in before-stimulation zone. “*”indicates statistical difference with zero (dashed horizontal line), evaluated by one-sample Student’s paired t-test to zero. Significant p values from left to right: 3.94×10^−2^; 3.07×10^−2^; 1.65×10^−3^; 3.70×10^−2^. Note that in pre-stimulation period, the mice tended to run faster near landmarks, showing higher speed in preceding zone than before-stimulation zone. This trend was abolished during stimulation period. In later stimulation runs, preceding zone exhibited lower speed than before-stimulation zone. **b.** Comparison between speed in before-stimulation and preceding zones in pre-, during-, and post-stimulation periods, evaluated by Student’s paired t-test for comparisons within individual stimulations. Each period was separated into the first 5 runs and the second 5 runs. Note that only the second 5 runs during stimulation (runs 16-20) showed significantly reduced speed in preceding zone. Significant p values from left to right: 3.43×10^−3^; 7.58×10^−4^; 1.27×10^−2^. **c.** Similar to **a** but for lick numbers. **d.** Similar to **b** but for lick numbers. Only the second 5 runs during stimulation (runs 16-20) showed significantly increased licking in preceding zone (p value: 5.60×10^−3^). CS data were from four mice, 10 sessions. RS data were from four mice, 12 sessions. *p ≤ 0.05, **p ≤ 0.01, ***p ≤ 0.001. n.s. p > 0.05. Error bars represent mean ± SEM.

A direct comparison of running speeds in the before-stimulation and preceding zones across the first and last five runs of each period confirmed the above trends. During the pre-stimulation period, mice consistently ran faster in the preceding zone. In contrast, during the stimulation period, speeds in the preceding zone became lower than in the before-stimulation zone, but only in the last five runs. No speed differences were detected post-stimulation (**Fig. 5b**). These results indicate that anticipatory slowing before CS onset emerged gradually, becoming apparent only in the later phase of the stimulation period.

Licking behavior followed a similar trajectory. During the pre-stimulation period, licking was low and comparable between preceding and before-stimulation zones (**Fig. 5c and 5d**, “Pre”). During stimulation, licking increased overall and became higher the preceding zone, but only in the last few runs (**Fig. 5c and 5d**, “Dur”). After stimulation, licking remained elevated for a few runs but no longer differed between zones, and eventually returned to pre-stimulation levels (**Fig. 5c and 5d**, “Post”). This pattern points to a gradual development of anticipatory licking during stimulation. Together, these results demonstrate that anticipatory reward-predictive behaviors—slowing and licking—gradually emerged during non-reward CS runs, supporting the formation of memory for stimulation locations. Therefore, we propose that, during reward-guided spatial learning, reward delivery reinforces consistent MEC activation, which in turn acts as a training signal that gradually establishes memory for reward locations. Our results further show that after stimulation ceased, mice briefly continued to seek rewards based on the newly established memory, as reflected by low speeds and high licking during the first few post-stimulation runs. However, without ongoing stimulation, the precision of these anticipatory behaviors quickly declined, represented as comparable speed and licking in before-stimulation and preceding zones (**Fig. 5**, the first 5 runs in “Post”). The speed and licking slowly returned to pre-stimulation levels. These dynamics likely reflect memory extinction once the consistent MEC activity that supported memory formation was removed.

### Spatially consistent MEC activation at pre-reward location enhances reward-predictive behaviors

We further investigated whether consistent MEC activation over a long distance before the reward enhances reward-predictive behaviors in mice that poorly learned the task. We applied CS within 15 cm before the reward (pre-reward CS) and compared it to randomized 15-cm stimulation (RS) (**Fig. 6a**). The experiments involved poor performers, defined by weak reward-predictive slowing and licking in the pre-stimulation period (**Fig. 6b and 6d**, see **Methods** for behavioral classification), reflecting a lack of track memory.

**Figure 6.**
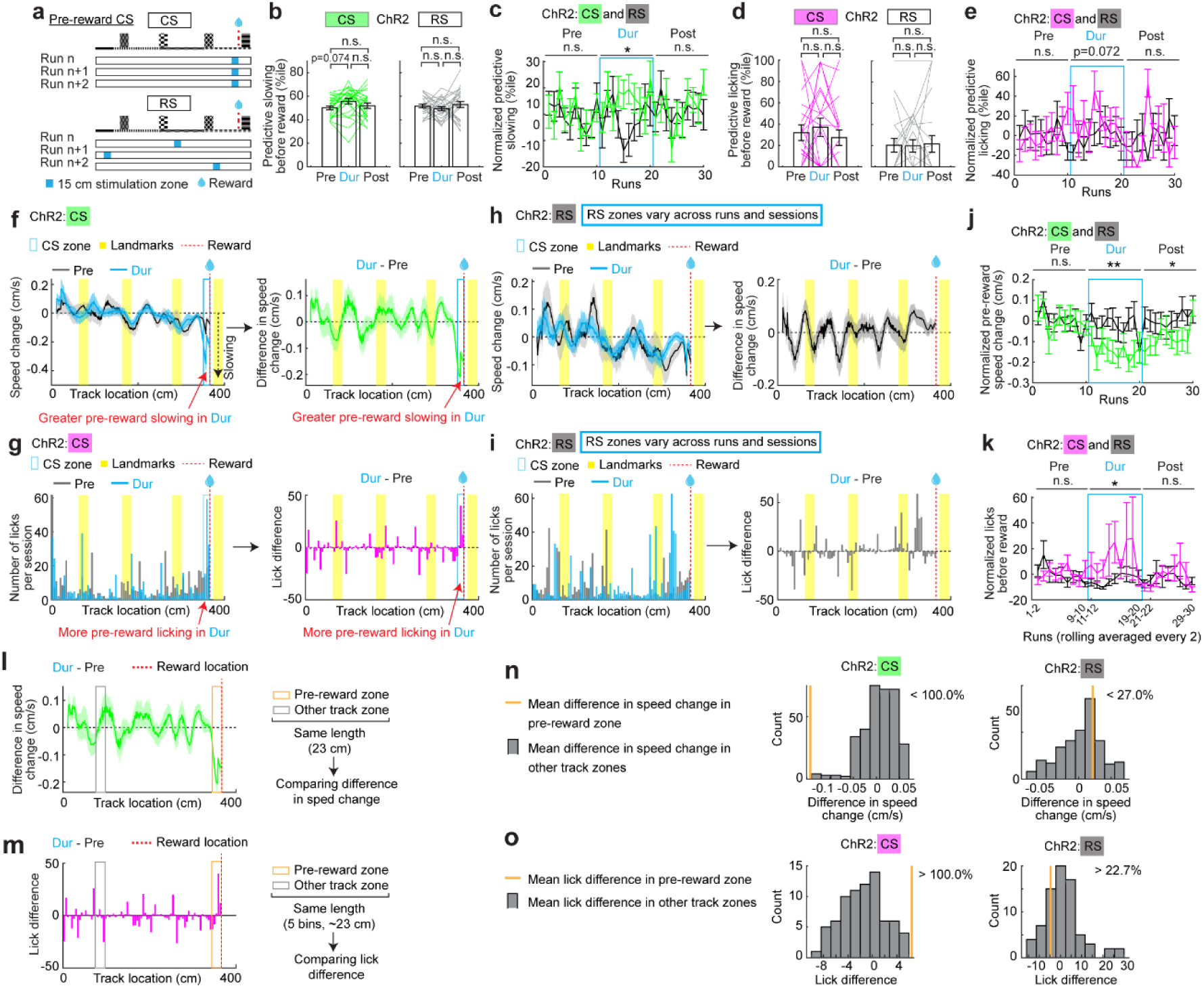
Spatially consistent optogenetic MEC stimulation at pre-reward locations enhances reward-predictive behaviors of ChR2 mice. **a.** Stimulation patterns. **b.** Predictive slowing of ChR2 mice, per CS or RS sesison. Student’s paired t-test was used between different periods of the same session. P values were adjusted for multiple comparisons using Bonferroni–Holm method. The same method was used in similar plots. **c.** RBR predictive slowing of ChR2 mice, CS vs RS. “n.s.” above horizontal lines indicates the p value comparing CS and RS during each period (two-way ANOVA). The same methods were applied to similar plots. Significant p value for Dur: 1.77×10^−2^. **d.** Predictive licking of ChR2 mice, per CS or RS sesison. **e.** RBR predictive licking of ChR2 mice, CS vs RS. **f.** Difference in slowing between pre- and during-CS periods for ChR2 mice. Only track locations before reward delivery were analyzed. The data in the first 10 cm on the track was removed due to teleportation artifact. Slowing was quantified as the speed change between consecutive spatial bins (speed in the current bin minus that in the previous bin), with negative values indicating slowing. Left: slowing in pre- (gray) and during-stimulation (blue), represented as the mean (solid curve) and SEM (shade) of the slowing across sessions (run-averaged within a session). Right: session-by-session difference in slowing (during-CS minus pre-CS). Negative values indicate greater slowing during stimulation. **g.** Licking difference between pre- and during-CS periods for ChR2 mice. Only track locations before reward delivery were analyzed. Left: spatially binned lick numbers during pre-CS (gray) and during-CS (blue), shown as the mean across sessions (all runs pooled within each session due to variable run-by-run licking). Right: difference in licking from pre- to during-CS, calculated by subtracting lick numbers in the pre-CS period from those in the during-CS period on a session-by-session basis. **h.** Similar to **f** but for ChR2 mice during RS. **i.** Similar to **g** but for ChR2 mice during RS. **j.** Comparison of pre-reward slowing (i.e., speed change) of ChR2 mice, CS vs RS. RBR values per session were normalized by subtracting the session’s mean pre-stimulation value. Group difference between CS and RS in each period was evaluated by two-way ANOVA (“*” above horizontal line). Significant p values: Dur: 3.34×10^−3^; Post: 3.60×10^−2^. **k.** Comparison of pre-reward lick numbers of ChR2 mice, CS vs RS. Due to invalid per-run and per-session values, rolling 2-run averaged lick numbers in a session were normalized by subtracting all sessions’ mean pre-stimulation value. Student’s t-test was used to evaluate the group difference of the combined normalized RBR values between CS and RS in each period (“*” above horizontal line). Significant p value for Dur: 1.09×10^−2^. **l.** Schematic for calculating stimulation-induced slowing (i.e., speed change) difference in pre-reward zone (23 cm before reward, determined based on the slowing and licking location of good-performing mice in **Fig. S11 and S12**, pre-stimulation runs). The calculation was conducted on the data in the right panel of **f**. Slowing difference in pre-reward zone was the mean across the zone. Slowing differences were similarly calculated in other track zones, which had the same length as the pre-reward zone and were chosen on a rolling basis (1-5 cm, 2-6 cm, 3-7 cm, ……). The difference in each zone was the average of all sessions within the zone. **m.** Schematic for calculating stimulation-induced licking difference in pre-reward zone (5 bins before reward, 4.58 cm per bin, total 22.9 cm, comparable to the 23 cm zone in **I**). The calculation was conducted on the data in the right panel of **g**. Lick difference in pre-reward zone was the mean across the 5 bins. Lick differences were similarly calculated in other track zones, which had the same length as the pre-reward zone and were chosen on a rolling basis (bins 1-5, 2-6, 3-7, ……). The difference was the average of all sessions within the zone. **n.** Slowing differences in pre-reward zone (orange) compared to other track zones (gray), shown as histograms. The percentage indicates the fraction of other zones with slowing improvement smaller than that observed in pre-reward zone (i.e., shifted toward more positive values relative to that in pre-reward zone). **o.** Lick differences in pre-reward zone (orange) compared to other track zones (gray), shown as histograms. The percentage indicates the fraction of other zones with licking improvement smaller than that in pre-reward zone (i.e., shifted toward more negative values relative to that in pre-reward zone). CS data were from nine mice, 25 sessions. RS data were from nine mice, 26 sessions. *p ≤ 0.05, **p ≤ 0.01, ***p ≤ 0.001. n.s. p > 0.05. Error bars represent mean ± SEM.

Remarkably, pre-reward CS improved predictive slowing of ChR2 mice per session compared to the pre-stimulation period, an effect not seen with RS (**Fig. 6b and 6c**). Predictive licking showed a modest trend toward increase across sessions (**Fig. 6d**), and a strong increase trend when comparing CS to RS across runs (**Fig. 6e**). No effects were observed in EGFP mice (**Fig. S16a-d**).

To directly assess whether pre-reward CS enhanced slowing and licking before reward location, we examined differences in these behaviors along the track between pre- and during-stimulation periods. During CS, mice slowed more strongly before the reward (i.e., greater negative speed change from one spatial bin to the next) (**Fig. 6f and S17a**) and showed increased pre-reward licking (i.e., higher lick numbers in **Fig. 6g and S17b**). These effects were absent during RS runs (**Fig. 6h, 6i, S17a, and S17b**). Quantifications confirmed that both behaviors were enhanced before reward during CS compared to RS (**Fig. 6j and 6k**). While slowing enhancement persisted across 10 runs (**Fig. 6j**), the licking increase became more pronounced in later stimulation runs (**Fig. 6k**, higher mean values in later runs of CS, also **Fig. S17b**), consistent with the gradual development of licking behavior during consistent stimulations (**Fig. 3r**).

We then asked whether these improvements were specific to the pre-reward location. We compared the improvements in pre-reward zone versus other track zones with equal length (**Fig. 6I and 6m**). We found that both slowing and licking improvement during CS exceeded 100% of those in other track zones (**Fig. 6n and 6o**, left), indicating specific effects for pre-reward zone. Such effects were not observed in RS (**Fig. 6n and 6o**, right) and in EGFP mice (**Fig. S16e-l**).

Overall, these findings demonstrate that spatially consistent MEC activation before the reward selectively enhances reward-predictive slowing and licking. This aligns with the strong correlation observed between pre-reward MEC activity and reward-predictive behaviors (**Fig. 1i-k**). Together, they suggest that pre-reward MEC activity is not merely correlated with, but also causally contributes to, memory-guided reward prediction.

### Spatially consistent MEC activation at pre-reward location enhances reward-predictive behaviors in the presence of landmark representation

In addition to the high spatial consistency at the pre-reward location, activity around landmarks showed the second-highest consistency, which further increased during learning (**Fig. 1g and 1h**). Our previous work showed that landmark-related consistency also distinguishes good from poor performers, with the latter exhibiting markedly lower consistency^12^. These findings suggest that landmark representation is another key component of the cognitive map, highlighting salient environmental features^9,10,12,26,27^. We therefore asked whether pre-reward stimulation would still robustly enhance reward-predictive behaviors when landmark representation was also present. To test this, we combined three 5-cm CSs at landmark locations and the 15-cm pre-reward CS (“combined CS”) in poor performers (**Fig. 7a**, top). The brief 5-cm CSs provided consistent landmark activation without significantly driving reward-predictive behaviors at the stimulation locations (**Fig. S13**), potentially mimicking the landmark representation of good performers, as suggested by our previous study^12^. The effects of combined CS were compared with RS using randomized stimulation blocks (**Fig. 7a**, bottom).

**Figure 7.**
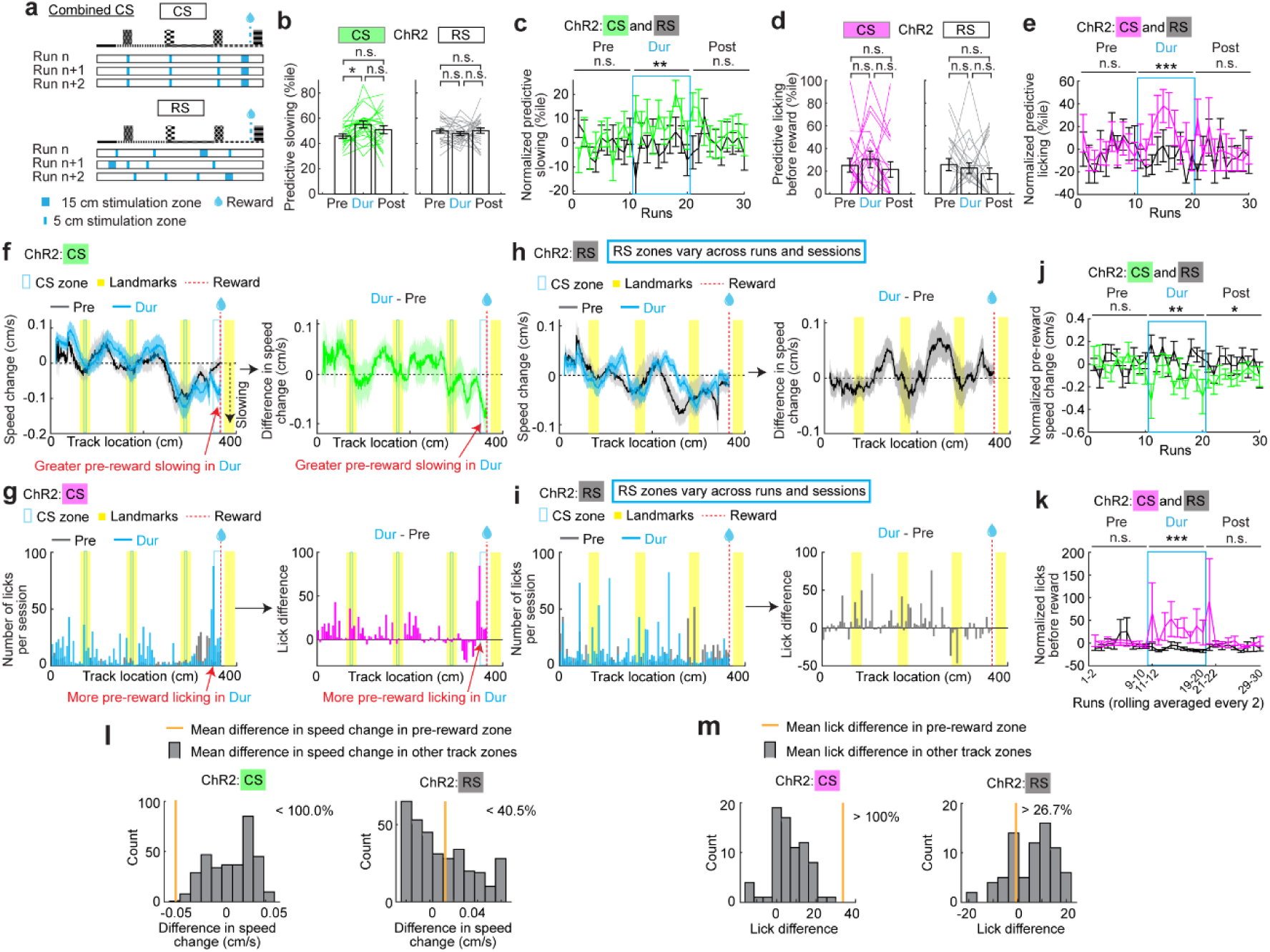
Combined consistent optogenetic MEC stimulations at pre-reward and landmark locations enhance reward-predictive behaviors of ChR2 mice. These panels are similar to those in Fig. 6 but for combined CS vs RS. **a.** Stimulation patterns. **b.** Predictive slowing of ChR2 mice, per CS or RS sesison. Student’s paired t-test was used between different periods of the same session. P values were adjusted for multiple comparisons using Bonferroni–Holm method. The same method was used in similar plots. Significant p value for CS, Pre vs Dur: 1.89×10^−2^. **c.** RBR predictive slowing of ChR2 mice, CS vs RS. “n.s.” above horizontal lines indicates the p value comparing CS and RS during each period (two-way ANOVA). The same methods were applied to similar plots. Significant p value for Dur, CS vs RS: 1.88×10^−3^. **d.** Predictive licking of ChR2 mice, per CS or RS sesison. **e.** RBR predictive licking of ChR2 mice, CS vs RS. Significant p value for Dur, CS vs RS: 2.64×10^−5^. **f.** Difference in speed change (representing slowing) between during- and pre-stimulation peroids for CS of ChR2 mice, before reward delivery. **g.** Licking difference between during- and pre-stimulation peroids for CS of ChR2 mice, before reward delivery. **h.** Similar to **f** but for RS of ChR2 mice. **i.** Similar to **g** but for RS of ChR2 mice. **j.** Comparison of pre-reward slowing (i.e., speed change) of ChR2 mice, CS vs RS. Group difference between CS and RS in each period was evaluated by two-way ANOVA (“*” above horizontal line). Significant p values for CS vs RS: Dur: 1.92×10^−3^; Post: 2.59×10^−2^. **k.** Comparison of pre-reward lick numbers of ChR2 mice, CS vs RS. Student’s t-test was used to evaluate the group difference of the combined normalized RBR values between CS and RS in each period (“*” above horizontal line). Significant p value for Dur, CS vs RS: 3.79×10^−6^. **l.** Slowing difference (i.e., difference in speed change) in pre-reward zone (orange) versus those in other track zones (gray). **m.** Lick difference in pre-reward zone (orange) versus those in other track zones (gray). CS data were from ten mice, 22 sessions. RS data were from ten mice, 27 sessions. *p ≤ 0.05, **p ≤ 0.01, ***p ≤ 0.001. n.s. p > 0.05. Error bars represent mean ± SEM.

We observed that combined CS, but not RS, significantly enhanced predictive slowing in ChR2 mice (**Fig. 7b and 7c**). Although CS did not significantly enhance predictive licking per session (**Fig. 7d**), it significantly increased predictive licking per run compared to during-RS runs (**Fig. 7e**). Combined CS also resulted in greater slowing and licking before the reward compared to pre-CS and during-RS runs (**Fig. 7f-k, S17c and S17d**). The slowing increase during CS was specific to the pre-reward zone (**Fig. 7l**). Although CS induced a general licking increase along the track (**Fig. 7g**), the increase near the reward exceeded those in other track zones (**Fig. 7m**). These effects were absent in EGFP mice (**Fig. S16m-x**). Overall, combined CS robustly increases both slowing and licking, especially near the reward.

Together, our results demonstrate that consistent pre-reward activation, whether alone or in conjunction with landmark activation, enhances reward-predictive slowing and licking, supporting the ability of this activation to robustly engage reward prediction.

## Discussion

Our study provides crucial evidence that the hippocampal-entorhinal cognitive map is fundamental for spatial memory. We identified a strong correlation between pre-reward activity consistency and reward-predictive behaviors during spatial learning. The activity consistency at pre-reward locations was the highest compared to those near landmarks and in other locations, a feature observed across various cell types. These observations suggest that spatially consistent representation of pre-reward location across the MEC network drives reward-predictive behaviors. Aligning with this idea, by applying spatially consistent MEC stimulation at non-reward locations, we biased the behaviors to these locations, suggesting a reward prediction signal that is generated by integrating MEC activity and with consistent track features. Consistent stimulation also led to gradually developed memory about stimulation locations, as demonstrated by anticipatory reward-predictive behaviors before stimulation onsets. Furthermore, consistent MEC activation pre-reward, alone or when combined with short landmark activation, enhanced reward-predictive behaviors of mice with poor performance. Our findings demonstrate that MEC activation can drive memory-guided behavior, supporting a causal role for MEC activity within the circuitry underlying spatial memory.

Our finding that spatially consistent pre-reward MEC activation drives reward-predictive behaviors potentially explains several previous observations in reward-oriented spatial tasks, such as the enrichment of spatial fields near reward locations^12,18,19^ and improved spatial decoding around these sites^18,28^. The development of highly consistent MEC activation at reward locations likely signals more than reward presence. Instead, it may actively contribute to the neural computations underlying reward prediction and associated behavior. Consistent with this idea, we show that animals can gradually form a memory of stimulation locations through repeated, spatially consistent MEC activation.

During natural reward-guided navigation, repeated and location-specific inputs to MEC neurons may similarly strengthen synaptic representations of locations near reward, leading to reliable ensemble-level activation. Dopaminergic signals, which exhibit experience-dependent, reward-predictive ramping during reward/goal-directed navigation^29–32^, are well positioned to facilitate this process by promoting synaptic plasticity at behaviorally relevant inputs^33^. Over repeated experience, such dopamine-dependent plasticity may stabilize pre-reward ensemble activity, supporting the formation of a predictive representation of reward location that guides behavior. Consistent with this idea, our results demonstrated that pre-reward, rather than post-reward, MEC activity consistency selectively increases during learning. Our optogenetic manipulation effectively bypasses this gradual learning process by directly imposing spatially structured ensemble activity in MEC, thereby inducing reward-predictive behavior at both pre-reward and non-reward locations. Notably, the memory of the stimulation location did not persist after stimulation ceased, resembling the extinction of reward-associated memory when reinforcing outcomes are no longer present.

A more direct test of these mechanisms would require simultaneous recording of neural activity during optogenetic stimulation^34^, which was not performed here. Although our electrophysiological recordings in anesthetized mice confirmed reliable, stimulation-locked increases in excitatory responses, it remains unclear how repeated, spatially localized stimulation during behavior influences activity consistency across the full track or the broader MEC network. Thus, while our results demonstrate that spatially consistent MEC activation is sufficient to evoke localized reward-predictive behavior, they do not establish a global increase in spatial coding stability or widespread learning-induced plasticity. Nevertheless, the observed predictive behaviors before stimulation onsets suggest that repeated MEC activation may have transiently strengthened synapses representing the stimulated locations, consistent with a local and experience-dependent enhancement of the cognitive map.

In addition to the MEC, the hippocampus also significantly contributes to the cognitive map of space through its reciprocal connections with the MEC^35^ and its place cells, which are active at a single or a few locations in an open arena^2^. MEC activity affects hippocampal neural dynamics^36^ and vice versa^37^. Therefore, MEC activation before reward likely corresponds with similar hippocampal activity, as reward-predictive place cell activity is shaped by entorhinal inputs^38^. Supporting these ideas and our findings, targeted activation of reward-zone place cells in a non-reward location biases reward-associated licking in mice^34^. Notably, this activation also led to a gradual formation of memory of the activation location, which was not maintained once the stimulation ended, consistent with our observations. Together, our study and the place cell study illustrate the sufficiency of the cognitive map in the hippocampal-entorhinal circuit for memory-guided reward prediction during navigation. Importantly, the behavioral effects are likely produced by distributed circuit interactions, rather than by the hippocampus or the MEC acting in isolation. Other connected brain regions, such as retrosplenial cortex^39^ and prefrontal cortex^40^, could also be involved in this process.

Beyond spatial memory, the MEC cognitive map also underlies other memory types. Optogenetic inhibition of MEC layer 2 island cells induces mouse freezing behavior during trace fear conditioning^41^. Activation of entorhinal cortex engram cell inputs to dentate gyrus (DG) engram cells restores engram-specific spine deficits in DG and recovers memories in contextual fear conditioning, inhibitory avoidance, and novel object recognition tasks^42^. Together with our findings, these studies provide strong evidence that the hippocampal-entorhinal cognitive map causally drives memory-guided behaviors across contexts. They also highlight the need to uncover underlying circuit mechanisms and pave the way for strategies to enhance memory by modulating cognitive map activity in clinical settings. Our study is particularly relevant in this regard, as we discovered that global modulation of MEC excitatory cells, without the additional need of functional-or molecular-cell-type-specific targeting^34,41,42^, can robustly improve goal-directed behavioral performance. This network-wide strategy points to a feasible clinical approach for memory enhancement through entorhinal modulation.

## Acknowledgments

We thank all members in the Gu laboratory for supporting the work, Daniel Yusufi, Dr. Lujia Chen, and Dr. Lorna Role for constructive suggestions on the manuscript, the Section on Instrumentation at National Institute of Mental Health for help with building the virtual reality setup, Dr. Joshua Gordon for guidance on *in vivo* electrophysiological recording, and Dr. Chris McBain for supporting Dr. June Hoan Kim during the execution of this work. This research was supported by the Intramural Research Program of the National Institutes of Health (NIH) to YG (ZIA NS009415), to Joshua Gordon (ZIA NS003168) and to Chris McBain (Z1A-HD001205-32). The contributions of the NIH authors were made as part of their official duties as NIH federal employees, are in compliance with agency policy requirements, and are considered Works of the United States Government. However, the findings and conclusions presented in this paper are those of the authors and do not necessarily reflect the views of the NIH or the U.S. Department of Health and Human Services.

## Author Contributions

Y.M., T.M., N.T., M.M, D.K., and Y.G. designed research; Y.M., T.M., N.T., J.T., F.S., and Y.G. performed research; Y.M., T.M., J.K., and Y.G. analyzed data; Y.M., T.M., J.K., N.T., J.T., and Y.G. wrote the paper.

## Supplementary information

**Figure S1.**
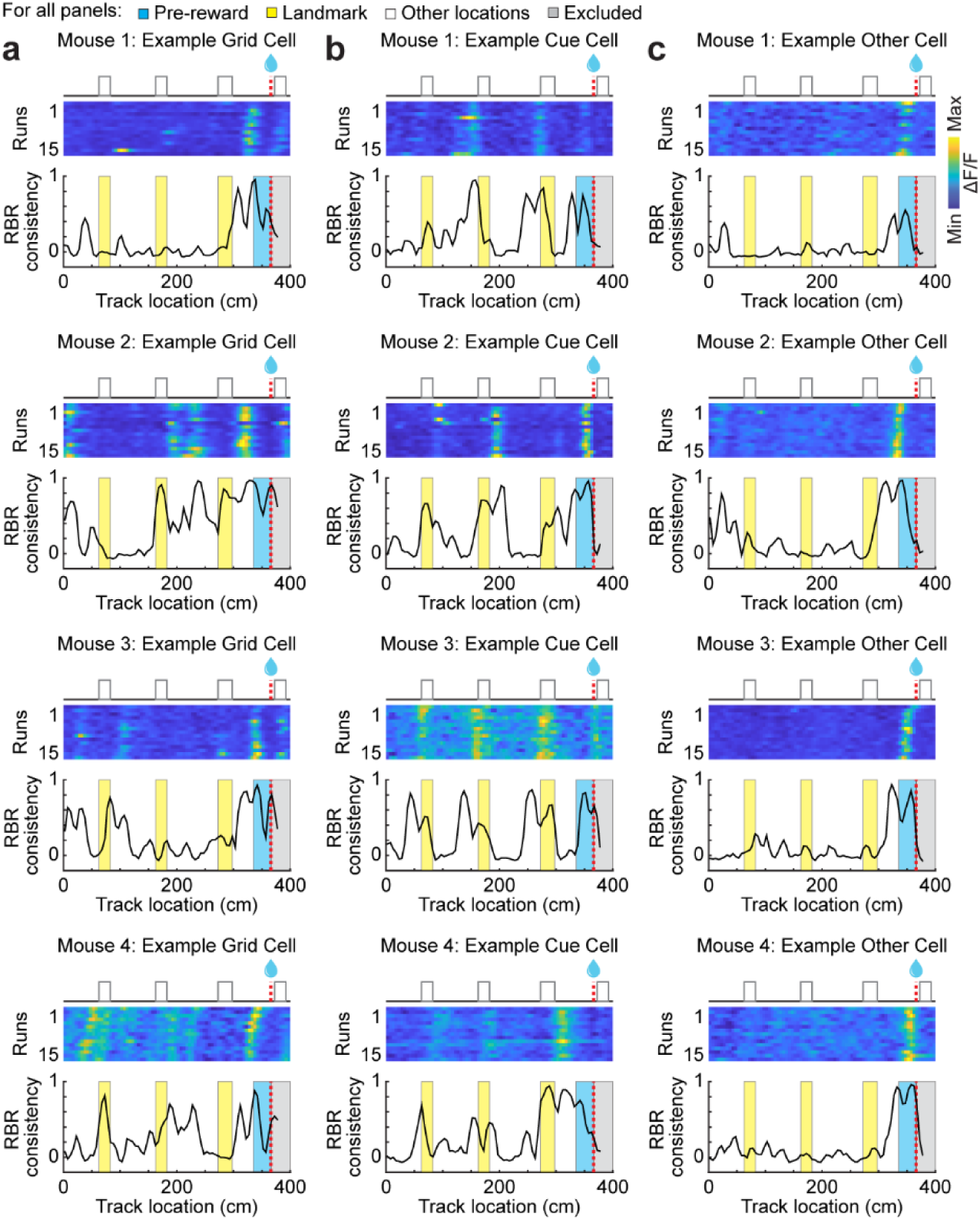
Example run-by-run consistency in individual cells. **a-c.** Top: Example run-by-run (RBR) calcium activity from example grid cells (**a**), cue cells (**b**), and other cells (**c**). Bottom: RBR activity consistency of the example cells.

**Figure S2.**
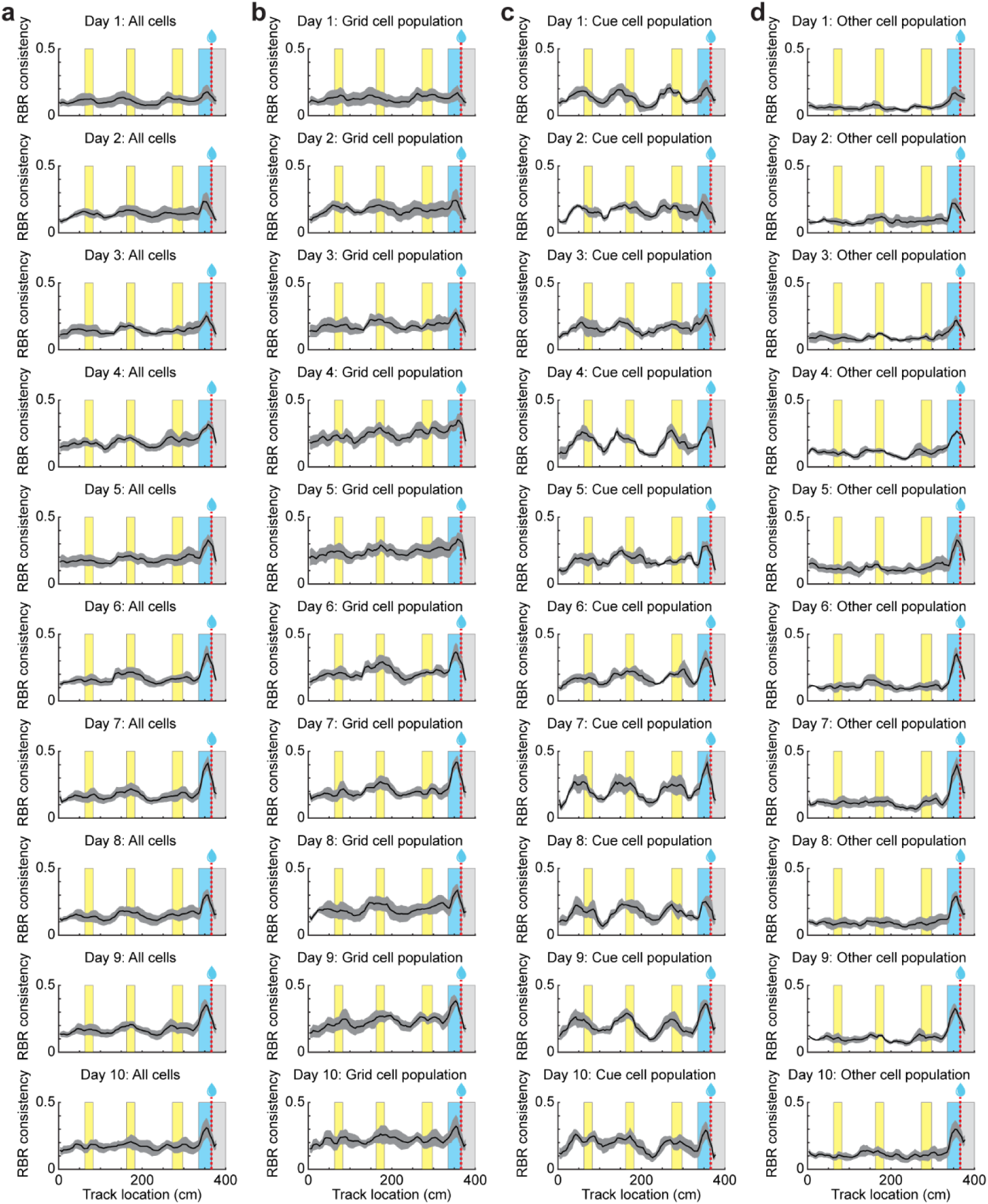
Mean spatial consistency across learning. **a-d.** Mean run-by-run (RBR) spatial consistency of all cells (**a**), grid cells (**b**), cue cells (**c**), and other cells (**d**) across the 10 days of learning averaged across mice. N = 4 mice from 7 FOVs. 15 ± 0 runs per mouse per day. Number of cells per mouse per day: **a.** 225 ± 16 cells; **b.** 105 ± 10 cells; **c.** 35 ± 3 cells; **d.** N = 85 ± 6 cells. Error bars represent mean ± SEM.

**Figure S3.**
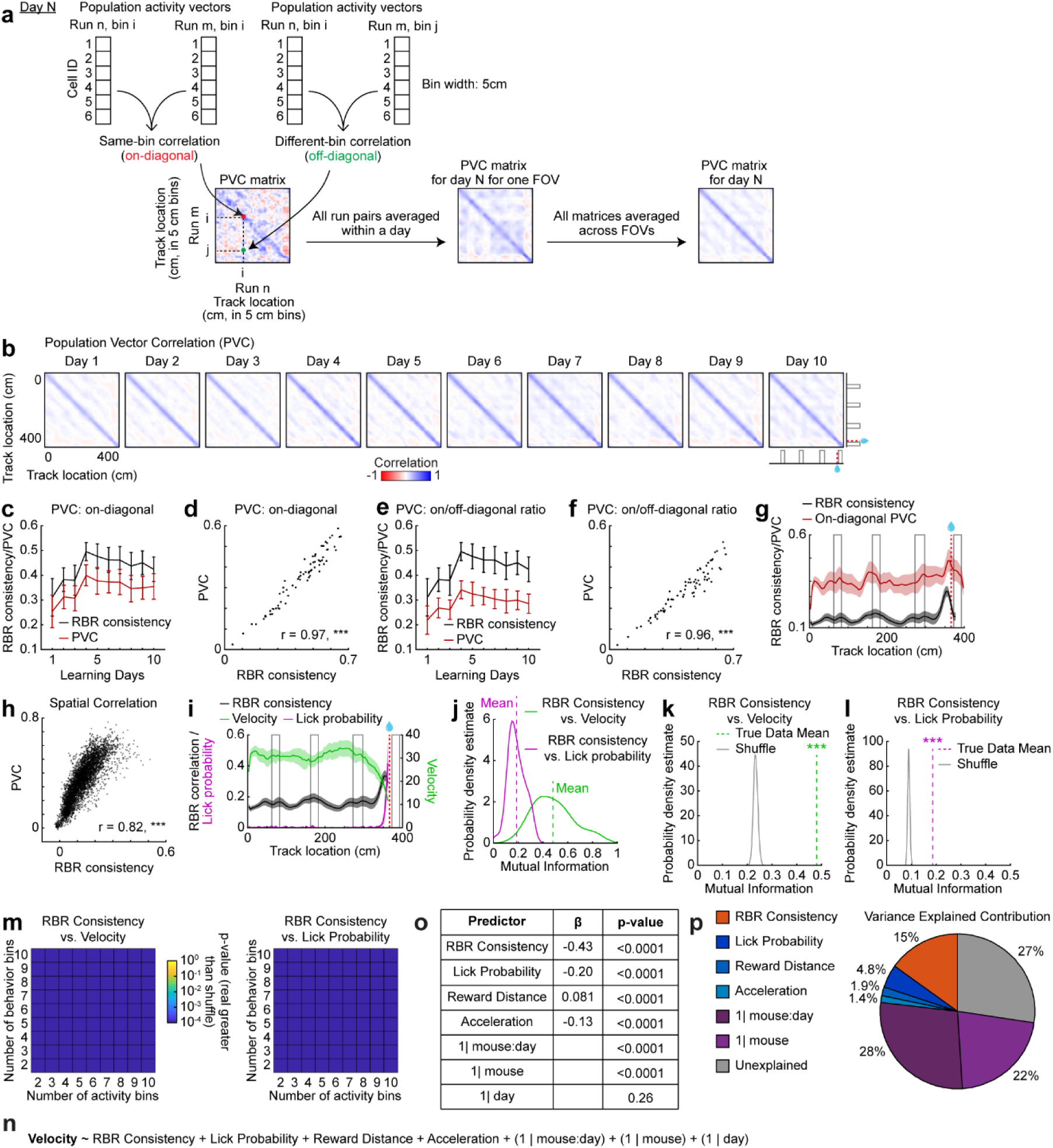
Run-by-run activity consistency captures MEC population coherence and predicts mouse behavior. **a.** Schematic for PVC calculation within a day. **b.** MEC population coherence as measured by run-by-run population vector correlation (PVC) between pairs of track locations across learning averaged across fields of view (FOVs). **c and e.** Run-by-run (RBR) activity consistency and on-diagonal PVC (**c**) or on/off-diagonal PVC ratio (**e**) across learning. N = 7 FOVs from 4 mice. **d and f.** Correlation between mean RBR activity consistency and on-diagonal PVC (**d**) or on/off-diagonal PVC ratio (**f**) across days. N = 70 sessions from 10 days and 7 FOVs from 4 mice. Peason Correlation. **d:** p=2.8×10^−45^; **f:** p=8.0×10^−38^. **g.** RBR activity consistency and on-diagonal PVC across track locations, averaged across late learning (days 7-10). N = 7 fields of view (FOVs) from 4 mice. **h.** Correlation between RBR activity consistency and on-diagonal PVC across pre-reward spatial bins (0-365 cm) and days. N = 5510 spatial bins from 73 bin locations, 10 days, and 7 FOVs from 4 mice. Peason Correlation. p=0. **i.** Run-by-run activity consistency, mouse velocity, and mouse lick probability across track locations, averaged across late learning. RBR consistency: N = 7 FOVs from 4 mice. Velocity/lick probability: N = 4 mice. **j.** Mutual information between run-by-run activity consistency and mouse velocity or lick probability. N=40 sessions from 4 mice across 10 days. RBR consistency was averaged across 1 to 2 FOV per mouse. Dashed lines indicate distribution means. **k and l.** Mutual information averaged across days compared to 10,000 shuffles for RBR consistency versus velocity (**k**) or lick probability (**l**). Each variable was discretized into 6 uniform bins for this calculation. Permutation test (compared to shuffle) **k**: p = 0.0001. **l:** p = 0.0001. **m.** P-values comparing mutual information between RBR consistency versus velocity (left) or lick probability (right) for various combinations of discretizing bin numbers. Permutation test (compared to shuffle). All p-values = 0.0001. **n.** GLMM equations used to predict mouse velocity from run-by-run activity consistency and behavioral variables. **o.** Summary table of fixed-effects estimates (β) and predictor significance (fixed effects: one-sample t-test; random effects: likelihood ratio test) from the GLMMs predicting mouse velocity. **p.** Relative contribution of significant predictors from the GLMMs predicting mouse velocity to behavioral variance explained based on dominance analysis. *p ≤ 0.05, **p ≤ 0.01, ***p ≤ 0.001. n.s. p > 0.05. Error bars represent mean ± SEM.

**Figure S4.**
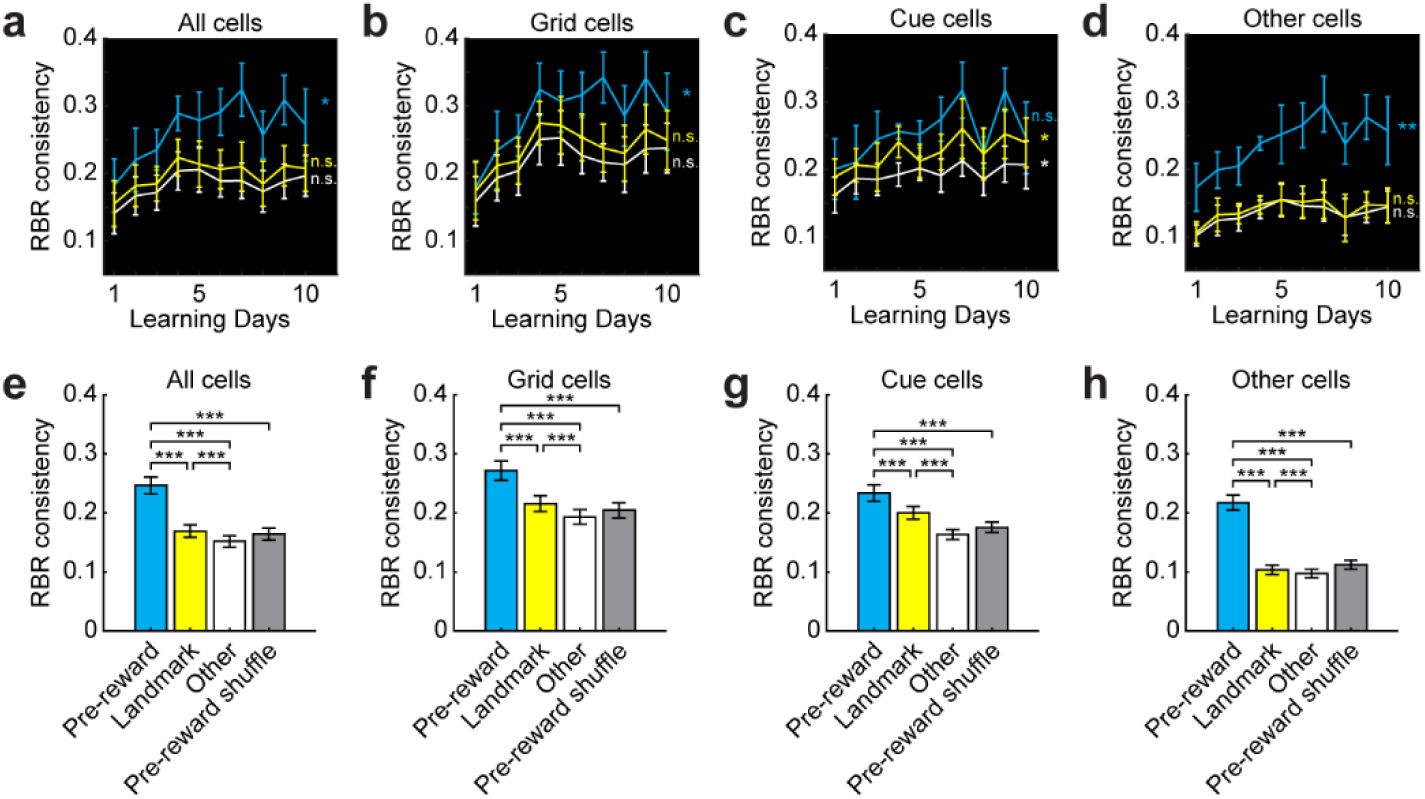
Comparison of spatial consistency types. **a-d**. Comparison of three consistency types, consistencies at reward, landmarks, and other locations, across 10 days for all cells (**a**), grid cells (**b**), cue cells (**c**), and other cells (**d**). N = 4 mice from pooled from 7 FOV (cells per mouse: **a**. 225 ± 16. **b**. 105 ± 10. **c**. 35 ± 3. **d**. 85 ± 6) with 15 ± 0 runs per day. Significant correlations (Pearson correlation) and p-values between mean consistency and learning day: **a**. pre-reward: r=0.70, p=0.023; **b**. pre-reward: r=0.70, p=0.024; **c**. landmark: r=0.76, p=0.011; other locations: r=0.74, p=0.014. **d**. pre-reward: r=0.80, p=0.0057. **e-h.** Comparison of the three consistency types and shuffled pre-reward consistency by pooling all values across 10 days for all cells (**e**), grid cells (**f**), cue cells (**g**), and other cells (**h**). Paired Student’s t-test. p values: **e**. pre-reward vs landmark: 7.02×10^−12^; pre-reward vs other locations: 5.44×10^−14^; landmark vs other locations: 1.02×10^−9^; pre-reward vs shuffle: 1.01×10^−13^. **f**. pre-reward vs landmark: 3.65×10^−6^; pre-reward vs other locations: 1.52×10^−9^; landmark vs other locations: 4.81×10^−9^; pre-reward vs shuffle: 4.17×10^−9^. **g**. pre-reward vs landmark: 0.0017; pre-reward vs other locations: 1.44×10^−9^; landmark vs other locations: 1.09×10^−7^; pre-reward vs shuffle: 8.02×10^−9^. **h**. pre-reward vs landmark: 3.26×10^−16^; pre-reward vs other locations: 1.71×10^−16^; landmark vs other locations: 0.0042; pre-reward vs shuffle: 1.86×10^−16^. *p ≤ 0.05, **p ≤ 0.01, ***p ≤ 0.001. n.s. p > 0.05. Error bars represent mean ± SEM.

**Figure S5.**
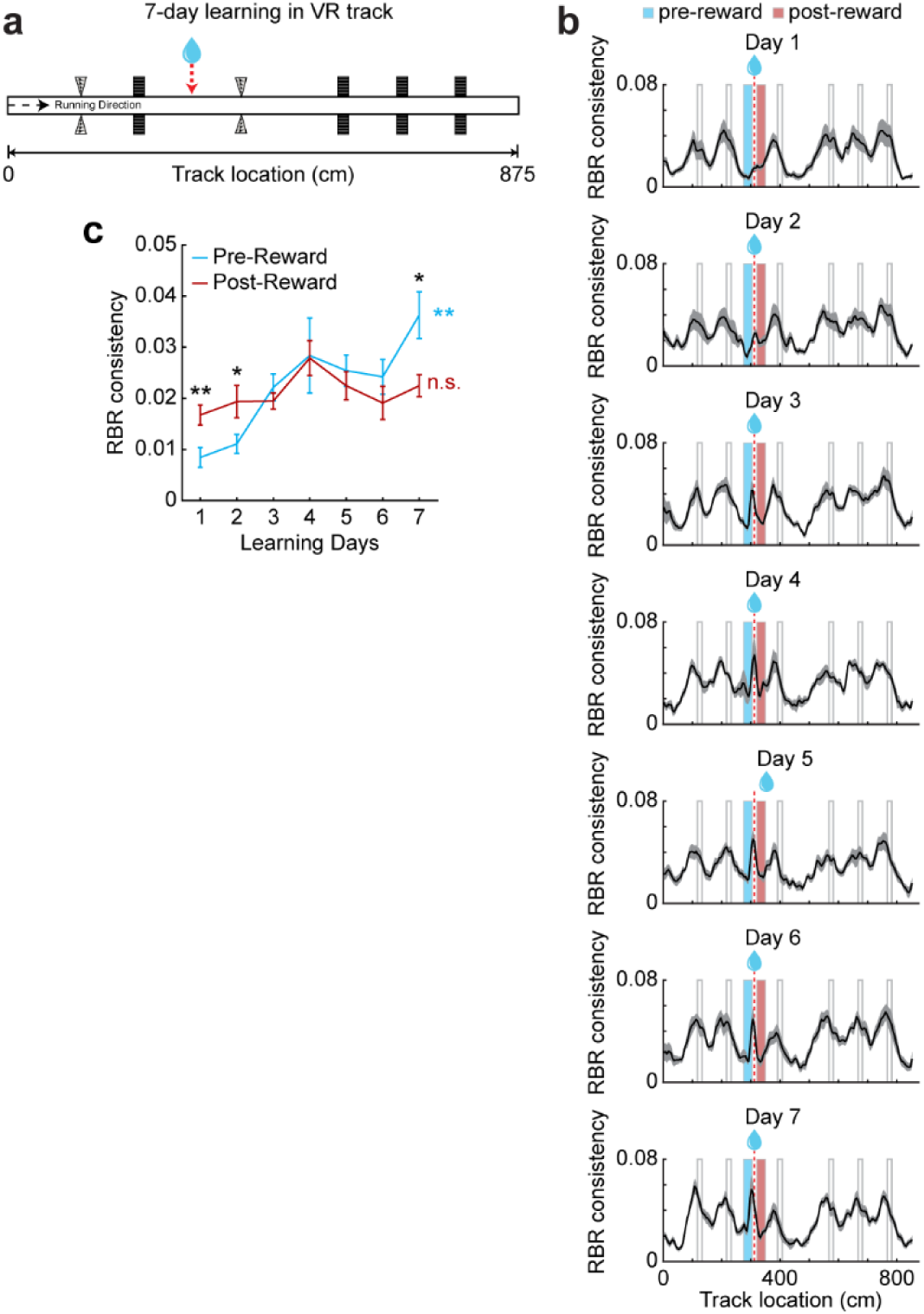
Comparison of pre-reward and post-reward spatial activity consistency. **a.** 875 cm track used in the dataset from^23^. **b.** Mean run-by-run (RBR) spatial activity consistency of all cells across 7 days of learning in the 875 cm track averaged across FOV. N = 10 FOV from 5 mice (177 ± 5.5 cells per FOV). **c.** RBR activity consistency across learning in the pre- and post-reward regions indicated in **b**. Significant p-values (Student’s t-test): Day 1, p=0.0077; Day 2, p=0.037; Day 10, p=0.014. Colored p-values indicate significant increase across learning (Pearson Correlation). Pre-reward: p=0.0057 *p ≤ 0.05, **p ≤ 0.01, ***p ≤ 0.001. n.s. p > 0.05. Error bars represent mean ± SEM.

**Figure S6.**
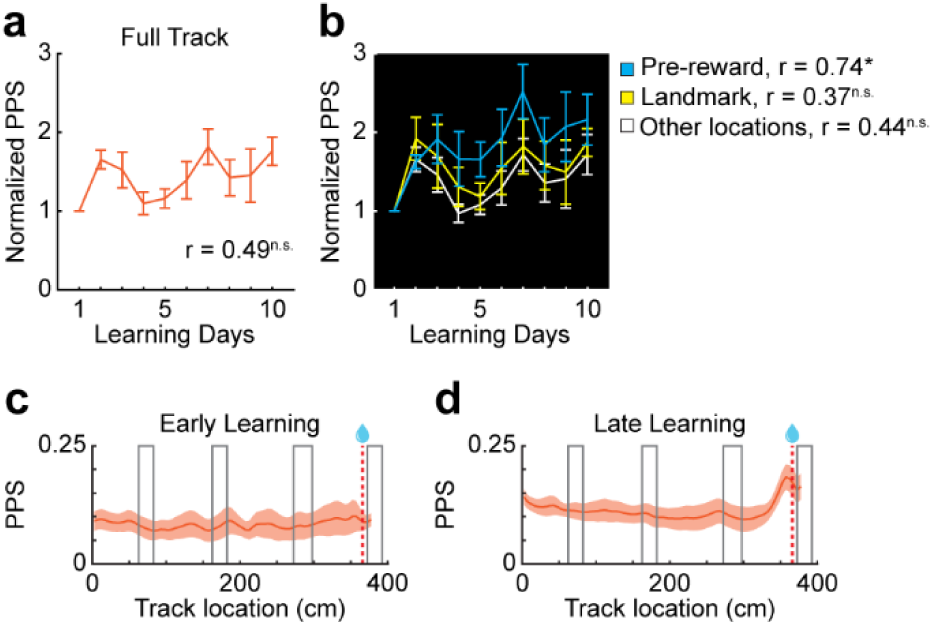
Pre-reward pairwise population synchrony increases during learning. **a.** Pairwise population synchrony (PPS) across learning normalized to day 1 by FOV. **b.** Comparison of pairwise population synchrony in the three spatial categories across learning normalized to day 1 by FOV. **a and b.** N = 7 FOVs from 4 mice. Significant correlations (Pearson correlation) and p-values between mean consistency and learning day: **b.** Pre-reward: p=0.015. **c and d.** Pairwise population synchrony across track locations, averaged across early (**c**, day 1) or late (**d**, days 7-10) learning. *p ≤ 0.05, **p ≤ 0.01, ***p ≤ 0.001. n.s. p > 0.05. Error bars represent mean ± SEM.

**Figure S7.**
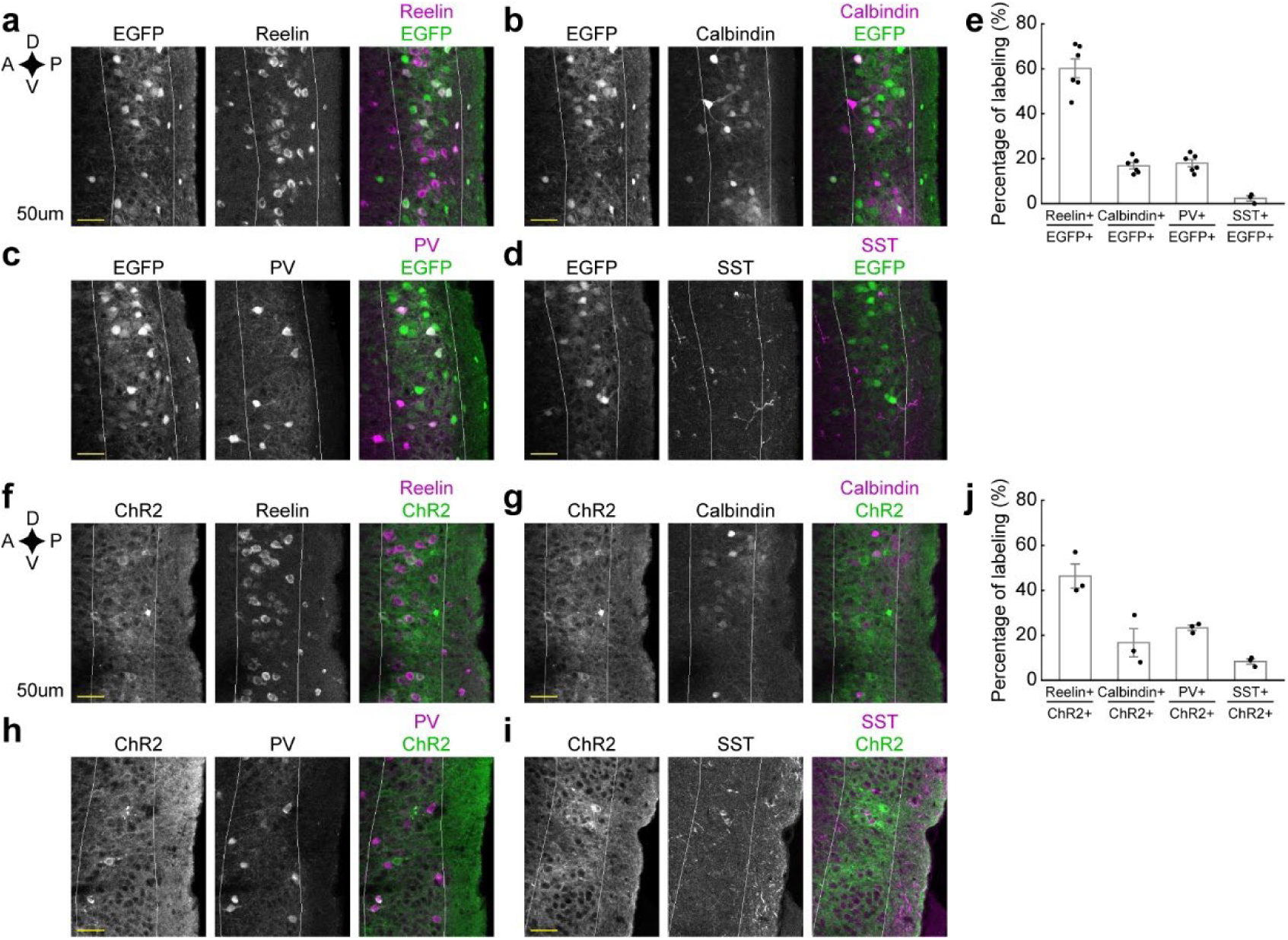
Synapsin promoter drives expression primarily in excitatory cells in MEC layer 2. **a-d and f-g.** Histological staining of Reelin (**a** and **f**), Calbindin (**b** and **g**), Parvalbumin (PV, **c** and **h**) and Somatostatin (SST, **d** and **i**) in the MEC for EGFP (**a-d**) or ChR2-GFP (**f-i**) injected mice. D: dorsal; V: ventral; A: anterior; P: posterior. **e and j.** Overlap of Reelin+, Calbindin+, PV+, and SST+ neurons with EGFP (**e**) or ChR2-GFP (**j**) expression. Each column represents the percentage of cells in the numerator category among cells in the denominator category. n = 3-6 FOV from 1-2 mice. Error bars represent mean ± SEM.

**Figure S8.**
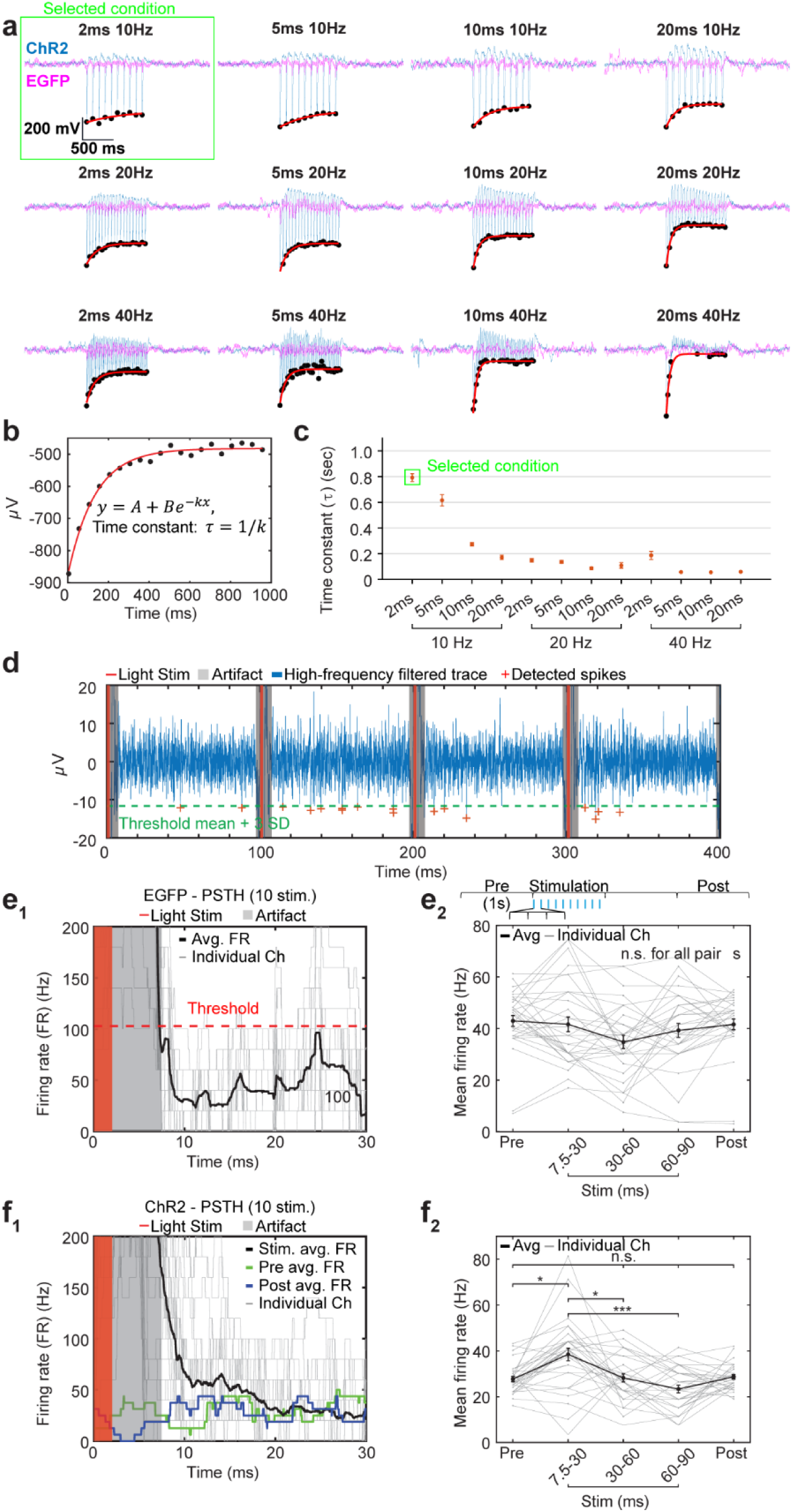
Optimization of optogenetic stimulation parameters and neuronal response dynamics. **a.** Representative low-frequency filtered traces (0.7–170 Hz) under 12 stimulation conditions (4 pulse durations × 3 frequencies) from ChR2- (blue) and EGFP-expressing (magenta) mice. Black dots indicate peak amplitudes and red lines represent exponential fits (*A* + *Be*^−*kx*^) for measuring stability of responses (decay time constant) in ChR2 traces. **b.** Example exponential fitting of peak decay. The time constant (*τ* = 1/*k*) was used to quantify response stability. **c.** Decay time constants across stimulation conditions. Data are average ± SEM from 32 channel signals of a ChR2 mouse. The 2 ms 10 Hz condition showed the slowest decay (i.e., greatest stability). **d.** Representative high-frequency filtered trace (250–8,000 Hz) showing spike detection (threshold: mean + 3 SD, green dashed line), stimulation period (red), and artifact period (gray). **e. (e_1_)** Determination of artifact exclusion window using EGFP recordings. Mean peri-stimulation time histograms (PSTH) showing mean firing rate (thick black) and firing rates in individual channels (32 channels, gray). The red dashed line indicates threshold (mean + 4.2 SD) defined as the time when the firing rate decayed to the threshold, which captured 90% of 1-ms binned baseline activity events (30ms after stimulation and 5 ms before the next stimulation). (**e_2_)** Comparison of mean firing rate across pre-stimulation (1-s window before stimulation), stimulation (three windows after first pulse of the stimulation train), and post-stimulation periods (1-s window following a 1-s stabilization period after the last stimulation pulse) in EGFP recordings; no significant differences (p=0.0528) were observed by Friedman test with post hoc multiple comparisons (Turkey-Kramer adjusted). **f. (f_1_)** Mean firing rate (thick black) in ChR2 recordings showing transient increase in firing rate followed by rapid decay (∼20 ms). Light gray lines represent firing rates in individual channels; green and blue lines indicate the 32-channel averaged firing rates during representative 100-ms windows before (pre) and after (post) the 1-s stimulation, respectively. (**f_2_)** Statistical comparison of mean firing rate across time windows (thick black: average, light gray: individual channels). Only the response during the first of three stimulation periods (7.5–30 ms) differed significantly (p=7.357×10^−5^, Friedman’s test); Pre vs 7.5-30: 0.0217; 7.5-30 vs 30-60: 0.0217; 7.5-30 vs 60-90: 1.299×10^−5^ *p ≤ 0.05, **p ≤ 0.01, ***p ≤ 0.001. n.s. p > 0.05. Error bars represent mean ± SEM.

**Figure S9.**
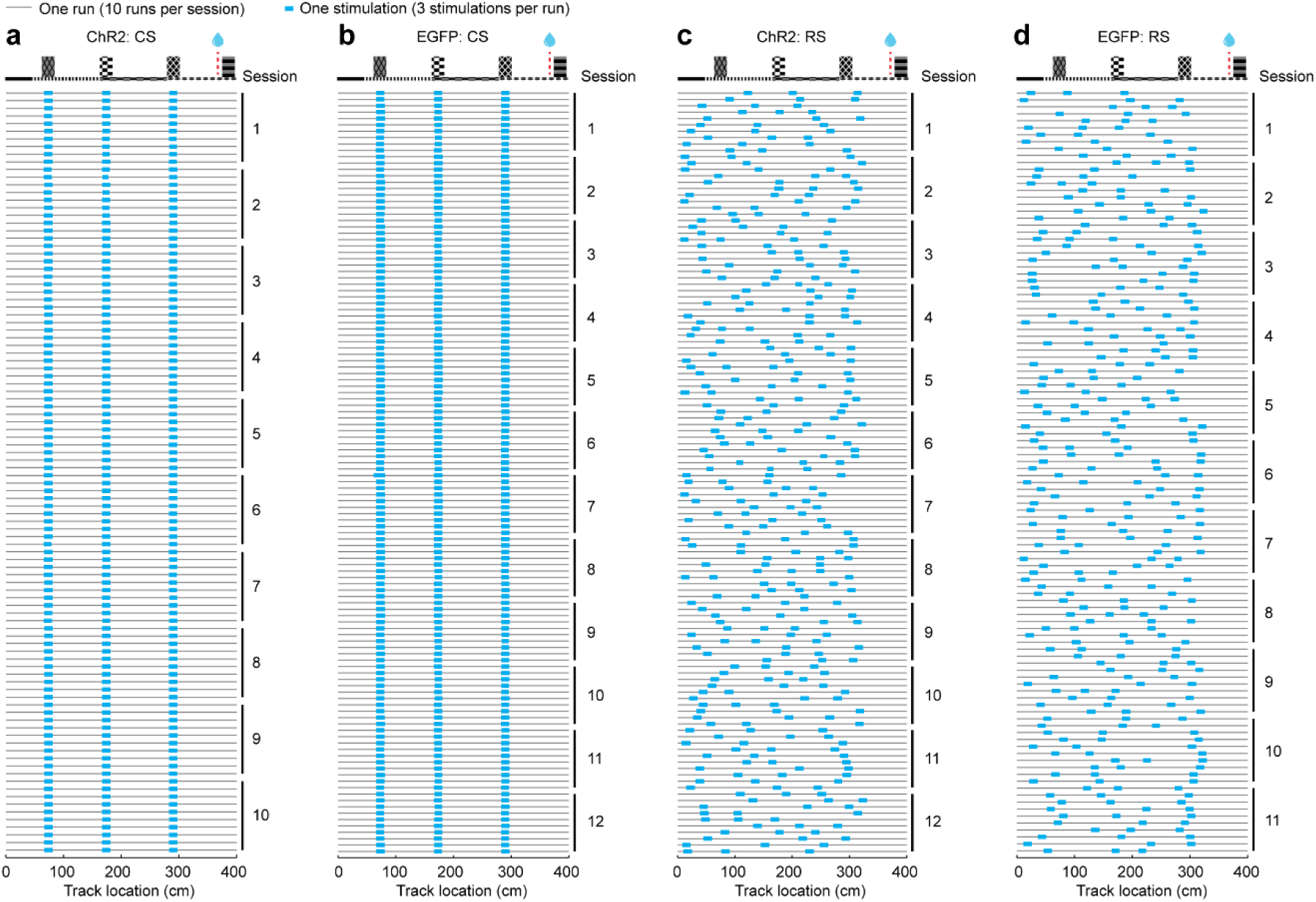
Run-by-run stimulation patterns. Run-by-run stimulation patterns of ChR2 CS (**a**), EGFP CS (**b**), ChR2 RS (**c**), and EGFP RS (**d**). All sessions in each condition are listed without specific order. Within each session, the 10 runs are displayed from top to bottom in the order they occurred during the actual experiments.

**Figure S10.**
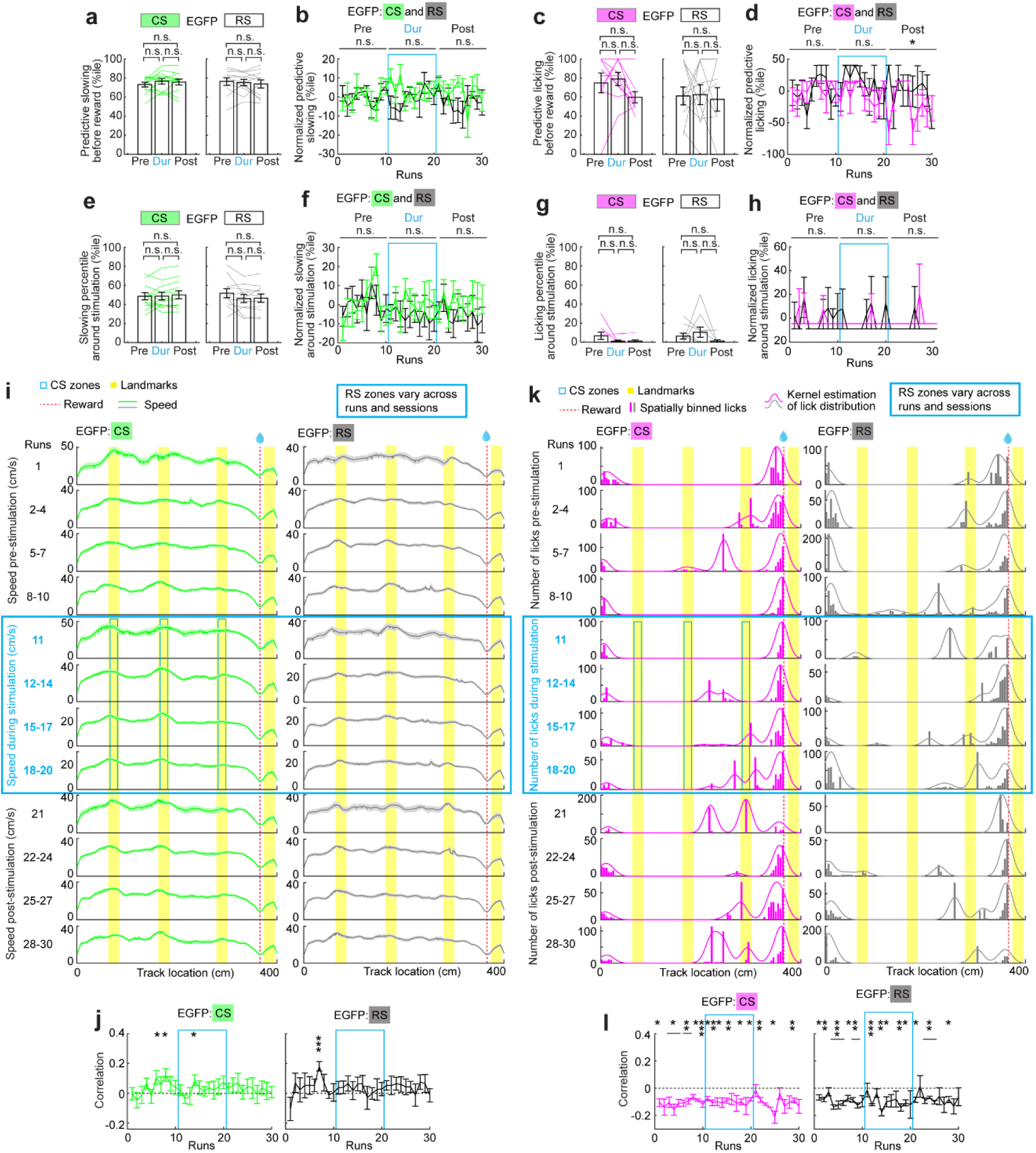
Behavioral effects of non-reward CS and RS on EGFP mice. **a.** Predictive slowing in pre-, during- (Dur), and post-stimulation per session. Student’s paired t-test was used between different periods of the same session. P values were adjusted for multiple comparisons using Bonferroni-Holm method. The same method was used in similar plots. **b.** RBR predictive slowing. “n.s.” above horizontal lines indicates the p value comparing CS and RS during each period (two-way ANOVA). The same methods were applied to similar plots. **c.** Predictive licking in pre-, during- (Dur), and post-stimulation per session. **d.** RBR predictive licking. Student’s t-test was used to compare CS and RS (“***” above horizontal line). This method was applied to similar plots below. Significant p values: CS vs RS Post: 1.28×10^−2^. **e.** Slowing percentile around stimulation zones, per session. **f.** RBR slowing percentile around stimulation zones. **g.** Licking percentile around stimulation zones, per session. **h.** RBR licking percentile around stimulation zones. **i.** Speed in pre-, during- and post-stimulation periods. **j.** Correlation between CS and slowing patterns. Significant p values from left to right (one-sample Student’s t-test): 4.00×10^−2^; 1.44×10^−2^; 3.63×10^−2^;1.77×10^−4^. **k.** Similar to **i** but for number of licks. **l.** Correlation between CS and licking patterns. Significant p values from left to right (one-sample Student’s t-test): 1.01×10^−2^; 3.51×10^−2^; 1.19×10^−2^; 2.58×10^−3^; 4.30×10^−3^; 4.30×10^−3^; 4.15×10^−3^; 2.05×10^−4^; 6.53×10^−3^; 1.08×10^−2^; 6.70×10^−3^; 3.65×10^−2^; 2.99×10^−3^; 3.91×10^−2^; 1.89×10^−2^; 2.71×10^−3^; 8.09×10^−4^; 1.79×10^−2^; 6.33×10^−3^; 1.85×10^−2^; 1.46×10^−3^; 3.20×10^−3^; 9.26×10^−3^; 1.64×10^−3^; 1.56×10^−2^; 7.44×10^−3^; 2.16×10^−3^; 5.01×10^−5^; 5.92×10^−3^; 4.20×10^−3^; 6.31×10^−3^; 3.69×10^−3^; 4.47×10^−2^; 8.77×10^−3^; 5.24×10^−3^; 3.44×10^−3^; 2.57×10^−2^. CS data were from three mice, 12 sessions. RS data were from three mice, 11 sessions. *p ≤ 0.05, **p ≤ 0.01, ***p ≤ 0.001. n.s. p > 0.05. Error bars represent mean ± SEM.

**Figure S11.**
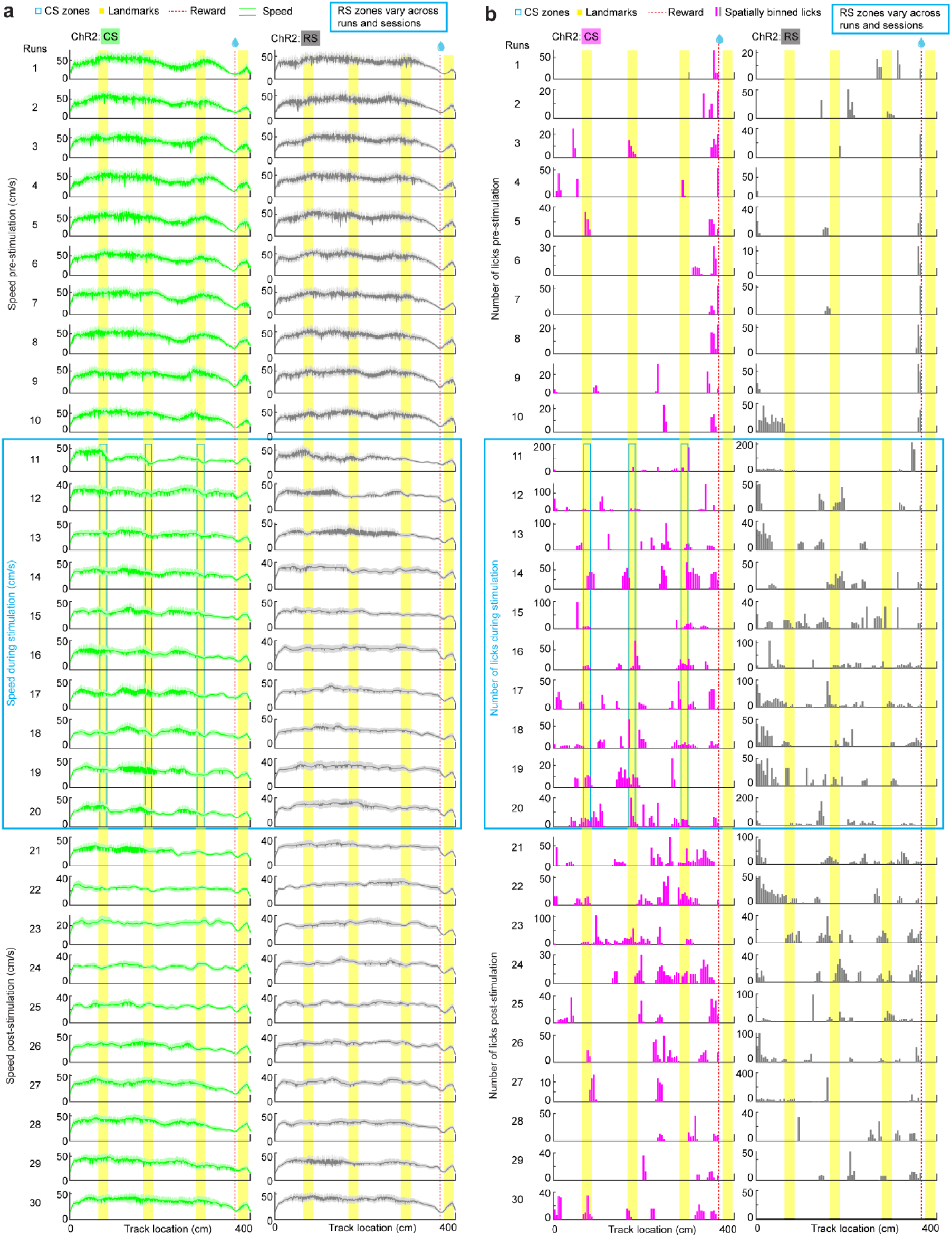
Run-by-run effects of non-reward CS and RS on ChR2 mice. **a.** Speeds in individual runs in pre-, during- and post-stimulation periods. **b.** Numbers of licks in individual runs in pre-, during- and post-stimulation periods. CS data were from four mice, 10 sessions. RS data were from four mice, 12 sessions.

**Figure S12.**
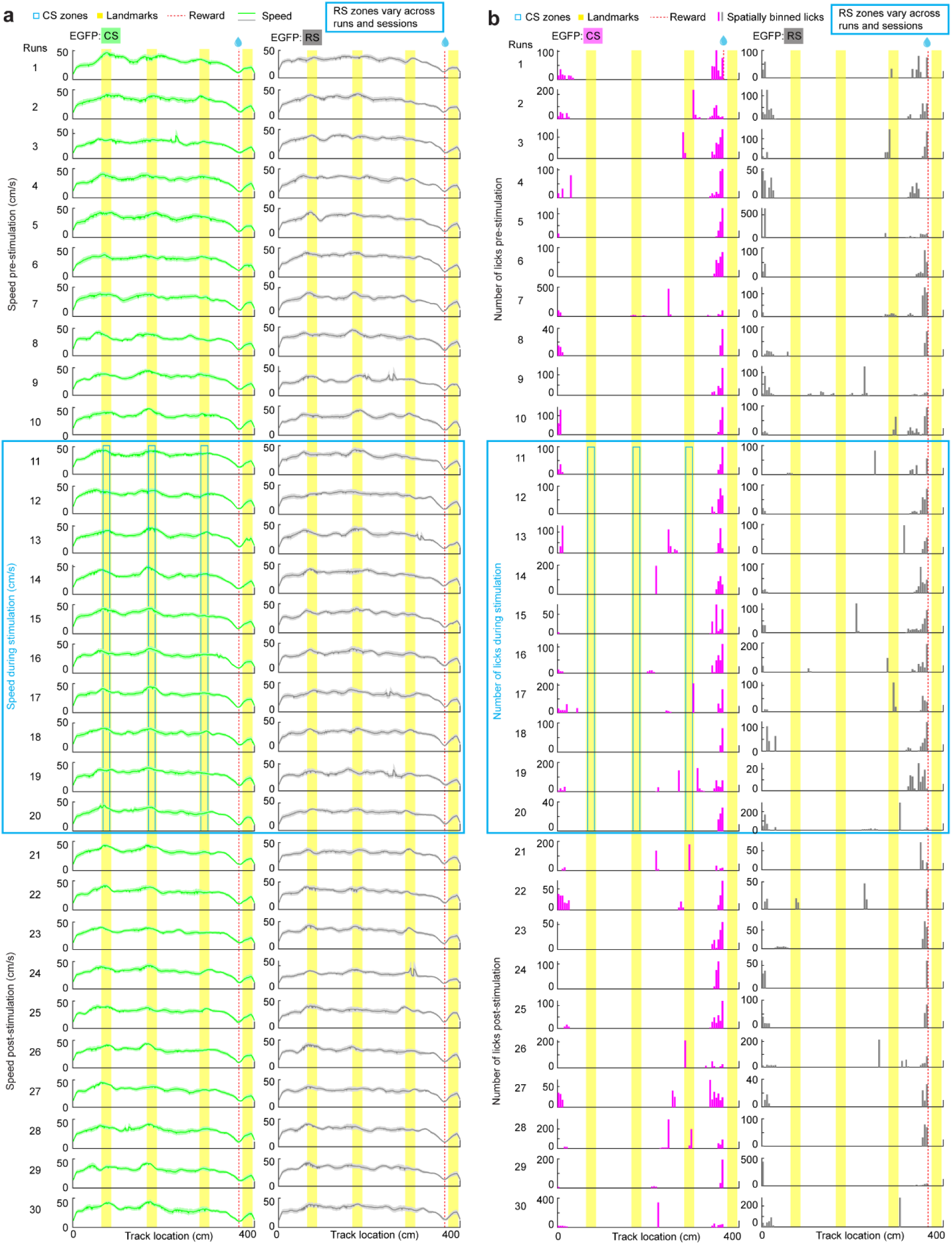
Run-by-run effects of non-reward CS and RS on EGFP mice. **a.** Speeds in individual runs in pre-, during- and post-stimulation periods. **b.** Numbers of licks in individual runs in pre-, during- and post-stimulation periods. CS data were from three mice, 12 sessions. RS data were from three mice, 11 sessions.

**Figure S13.**
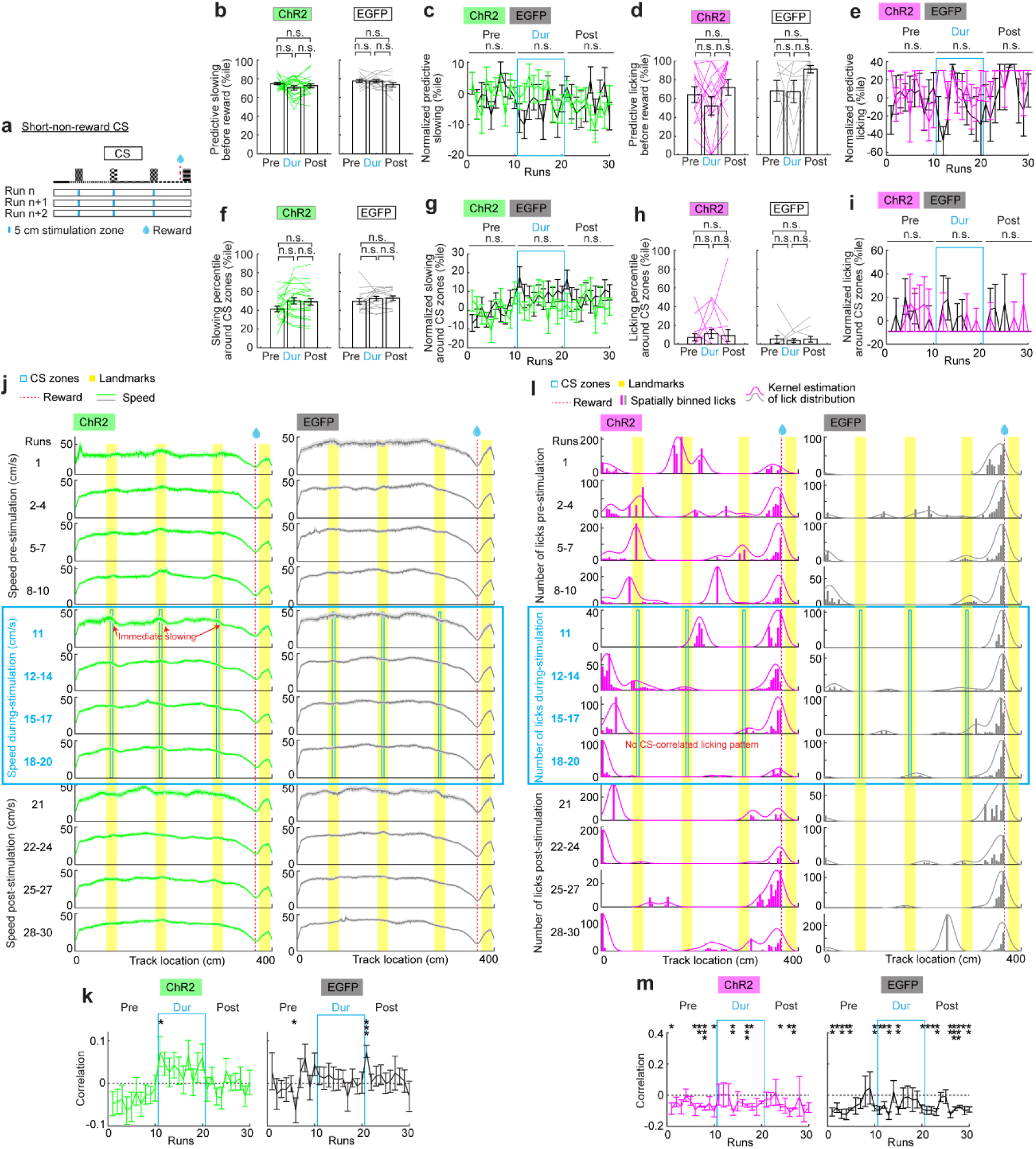
Spatially consistent optogenetic stimulation of the MEC of ChR2 and EGFP mice at 5-cm non-reward locations (short-non-reward CS) **a.** Stimulation patterns for short-non-reward CS. **b.** Predictive slowing of ChR2 and EGFP mice in pre-, during- (Dur), and post-stimulation. Student’s paired t-test was used between different periods of the same session. P values were adjusted for multiple comparisons using Bonferroni–Holm method. The same method was used in similar plots. **c.** RBR predictive slowing of ChR2 and EGFP mice. “n.s.” above horizontal lines indicates the p value comparing CS and RS during each period (two-way ANOVA). The same methods were applied to similar plots. **d.** Similar to **b** but for predictive licking. **e.** Similar to **c** but for RBR predictive licking. Student’s t-test was used to compare CS and RS (“***” above horizontal line). This method was applied to similar plots below. **f.** Slowing percentile around stimulation zones of ChR2 and EGFP mice. **g.** RBR slowing percentile around stimulation zone of ChR2 and EGFP mice. **h.** Licking percentile around stimulation zones of ChR2 and EGFP mice. **i.** RBR licking percentile around stimulation zone of ChR2 and EGFP mice. **j.** Speed in pre-, during- and post-stimulation periods of ChR2 and EGFP mice. **k.** Correlation between CS and slowing patterns of ChR2 and EGFP mice. Significant p values from left to right (one-sample Student’s t-test)a: 4.43×10^−2^; 4.78×10^−2^; 1.36×10^−4^. **l.** Similar to **j** but for number of licks. **m.** Correlation between CS and licking patterns of ChR2 and EGFP mice. Significant p values from left to right (one-sample Student’s t-test): 4.86×10^−2^; 4.10×10^−2^; 2.94×10^−3^; 3.30×10^−4^; 1.76×10^−2^; 5.72×10^−2^; 4.84×10^−4^; 1.65×10^−2^; 2.84×10^−2^; 2.89×10^−2^; 2.05×10^−4^; 9.49×10^−3^; 4.27×10^−2^; 9.44×10^−3^; 1.16×10^−2^; 8.19×10^−3^; 2.69×10^−3^; 3.10×10^−2^; 4.18×10^−2^; 2.85×10^−3^; 6.19×10^−3^; 2.62×10^−2^; 1.51×10^−2^; 1.44×10^−2^; 4.46×10^−3^; 2.34×10^−3^; 2.81×10^−4^; 1.70×10^−4^; 3.42×10^−2^; 2.87×10^−3^. ChR2 data were from eight mice, 23 sessions. EGFP data were from five mice, 15 sessions. *p ≤ 0.05, **p ≤ 0.01, ***p ≤ 0.001. n.s. p > 0.05. Error bars represent mean ± SEM.

**Figure S14.**
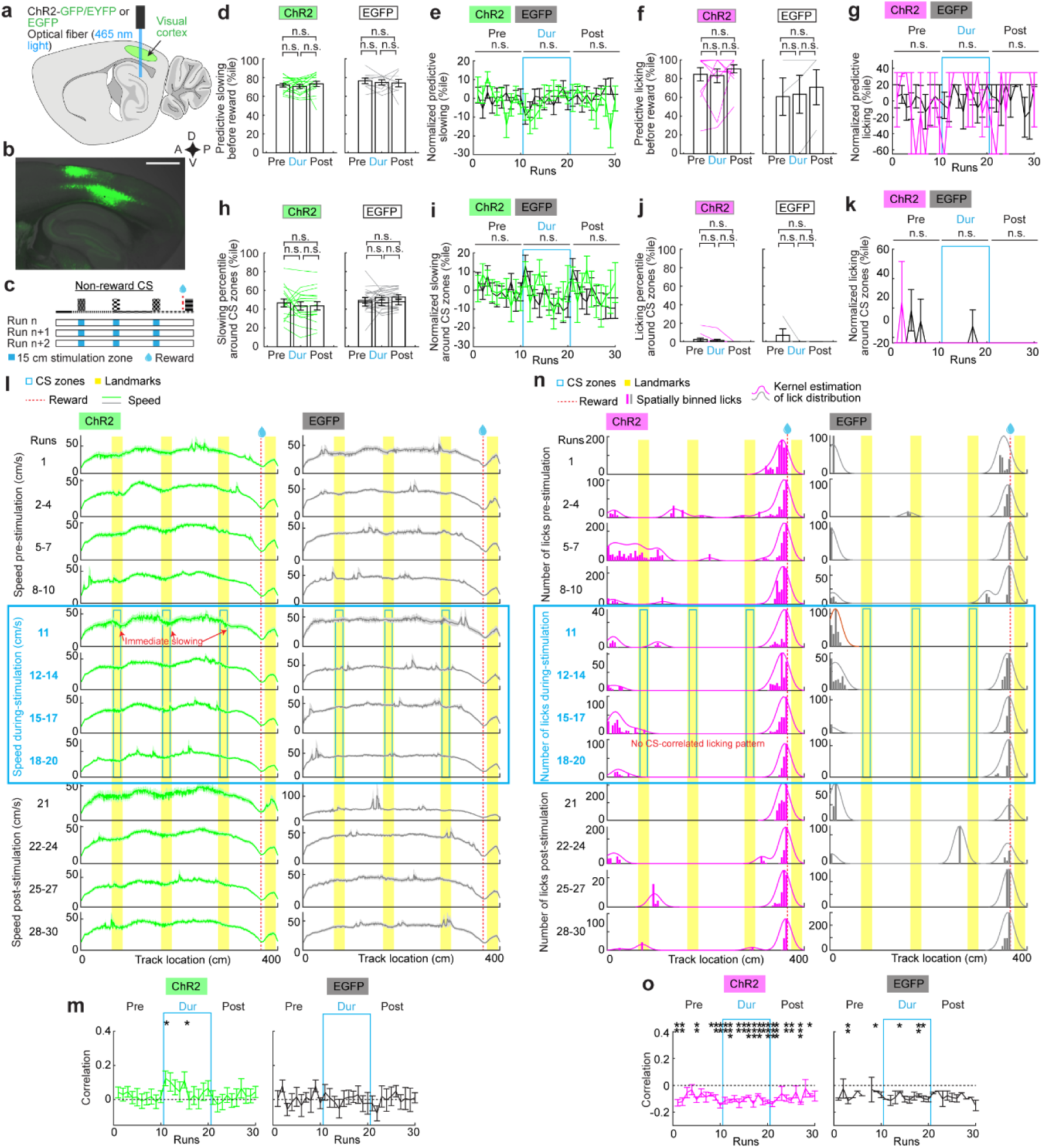
Optogenetic stimulation of visual cortex. **a.** Schematic displaying the location of the optical fiber and blue LED light delivery to activate ChR2-GFP/EYFP- or EGFP-expressing neurons in the visual cortex, adapted from^12^. To ensure consistency with the MEC experiments, the same fiber in Figure 3a was used. **b.** Epifluorescence image of a sagittal section showing ChR2-GFP expression in the visual cortex. Scale bar: 1mm. **c.** Stimulation patterns for non-reward CS, same as that in Figure 3d. **d.** Predictive slowing of ChR2 and EGFP mice in pre-, during- (Dur), and post-stimulation. Student’s paired t-test was used between different periods of the same session. P values were adjusted for multiple comparisons using Bonferroni–Holm method. The same method was used in similar plots. **e.** RBR predictive slowing of ChR2 and EGFP mice. “n.s.” above horizontal lines indicates the p value comparing CS and RS during each period (two-way ANOVA). **f.** Similar to **d** but for predictive licking. **g.** Similar to **e** but for RBR predictive licking. Student’s t-test was used to compare CS and RS (“***” above horizontal line). This method was applied to similar plots below. **h.** Slowing percentile around stimulation zones of ChR2 and EGFP mice. **i.** RBR slowing percentile around stimulation zone of ChR2 and EGFP mice. **j.** Licking percentile around stimulation zones of ChR2 and EGFP mice. **k.** RBR licking percentile around stimulation zone of ChR2 and EGFP mice. **l.** Speed in pre-, during- and post-stimulation periods of ChR2 and EGFP mice. **m.** Correlation between CS and slowing patterns of ChR2 and EGFP mice. Significant p values from left to right (one-sample Student’s t-test): 1.17×10^−2^; 2.72×10^−2^. **n.** Similar to **l** but for number of licks. **o.** Correlation between CS and licking patterns of ChR2 and EGFP mice (one-sample Student’s t-test). Significant p values from left to right: 1.72×10^−3^; 3.88×10^−3^; 3.99×10^−3^; 1.02×10^−2^; 6.79×10^−3^; 1.33×10^−3^; 3.46×10^−3^; 9.08×10^−6^; 6.39×10^−3^; 1.77×10^−3^; 5.94×10^−5^; 5.11×10^−4^; 9.82×10^−4^; 9.53×10^−3^; 6.87×10^−4^; 6.59×10^−4^; 1.93×10^−4^; 9.13×10^−3^; 3.41×10^−3^; 4.30×10^−4^; 2.64×10^−2^; 7.78×10^−3^; 3.25×10^−2^; 1.16×10^−2^; 7.55×10^−3^; 4.25×10^−2^. ChR2 data were from five mice, 16 sessions. EGFP data were from three mice, 9 sessions. *p ≤ 0.05, **p ≤ 0.01, ***p ≤ 0.001. n.s. p > 0.05. Error bars represent mean ± SEM.

**Figure S15.**
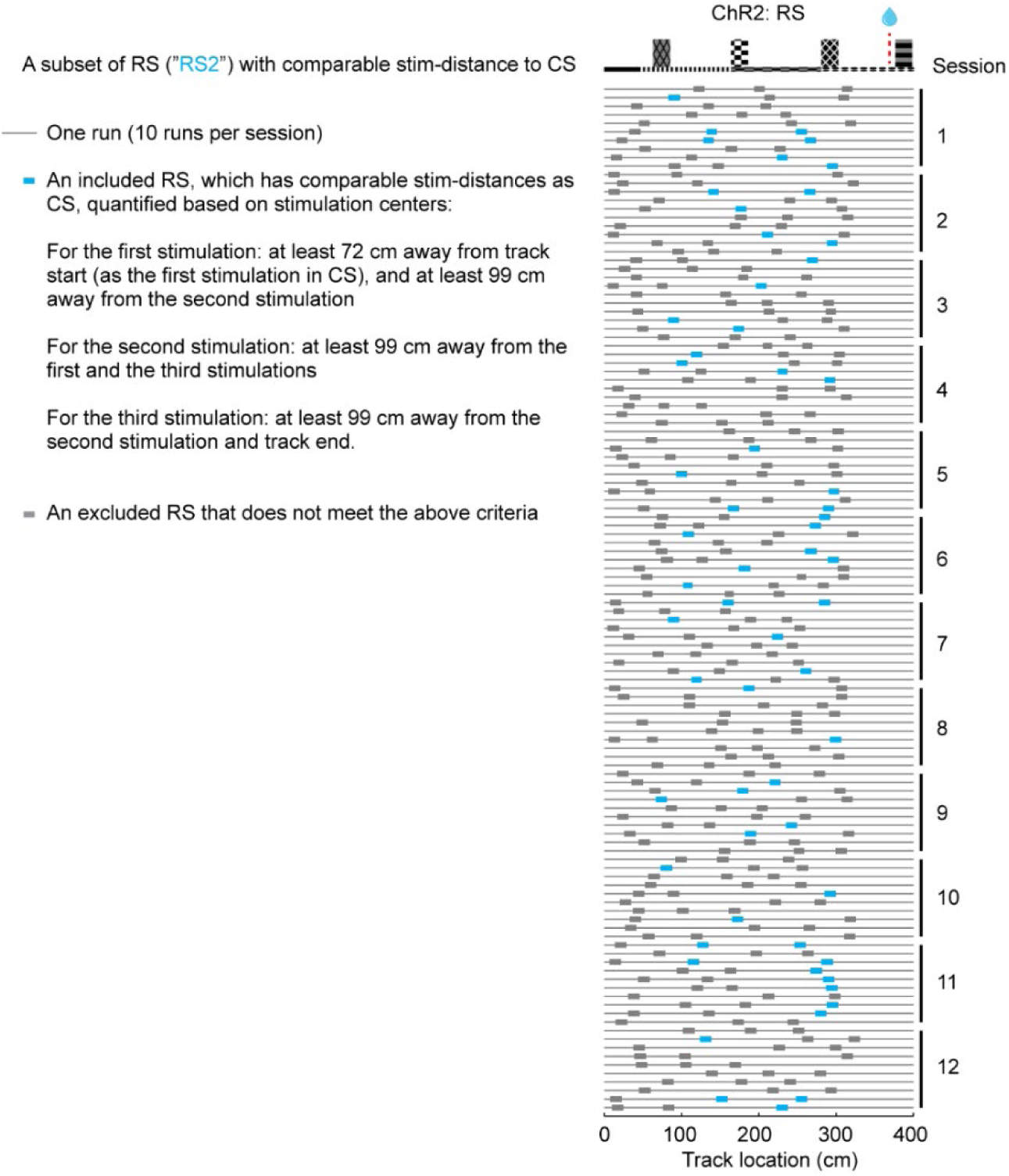
Sub-sampling RS (RS2) Left: the criteria for RS2. Right: similar **Figure S9c**, with unused RS marked in gray and used RS (RS2) marked in blue.

**Figure S16.**
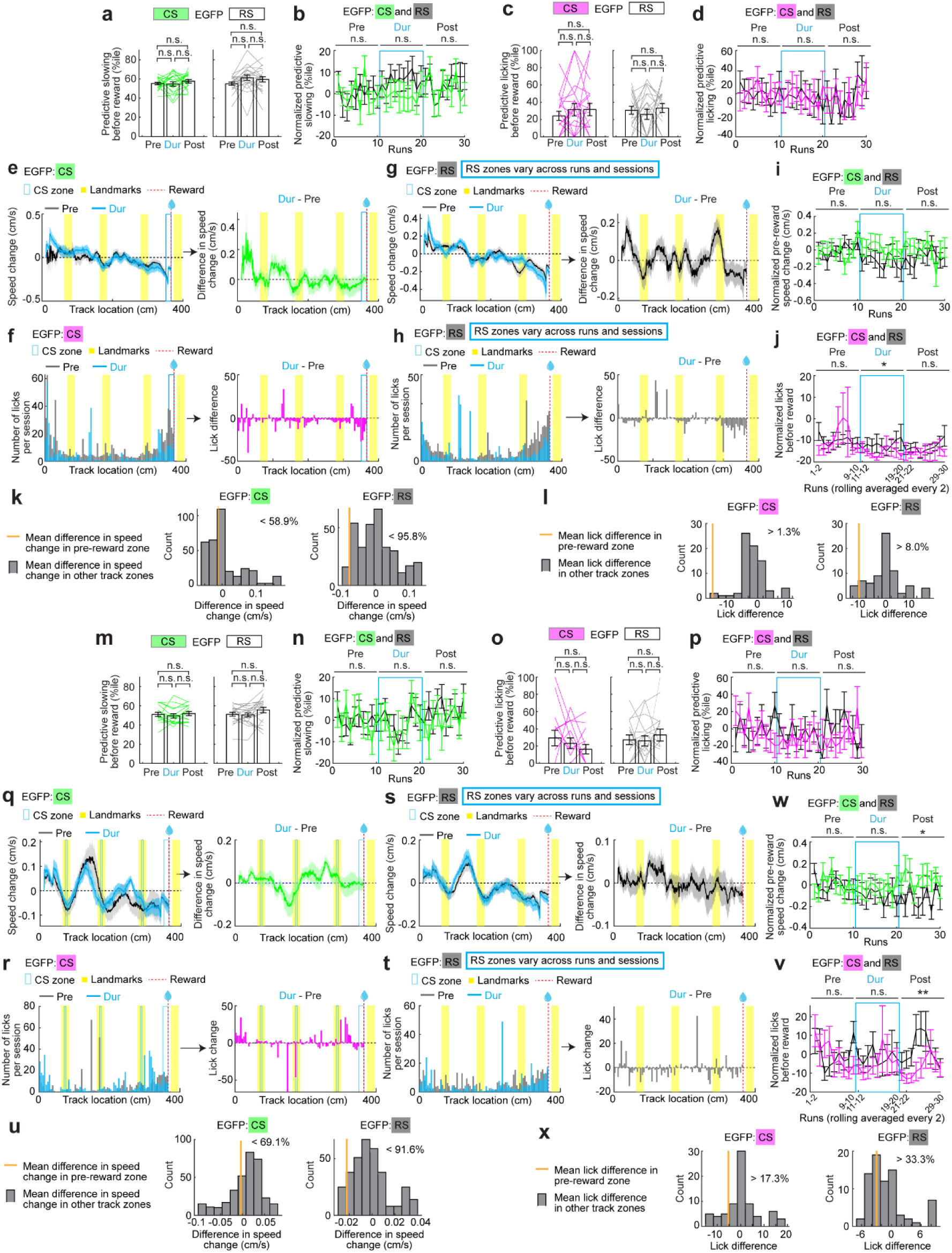
Behavioral effects of pre-reward and combined CS vs RS on EGFP mice. **a-l.** Pre-reward CS vs RS. **a.** Predictive slowing per session. Student’s paired t-test was used between different periods of the same session. P values were adjusted for multiple comparisons using Bonferroni–Holm method. The same method was used in similar plots. **b.** RBR predictive slowing. “n.s.” above horizontal lines indicates the p value comparing CS and RS during each period (two-way ANOVA). **c.** Predictive licking per session. **d.** RBR predictive licking. Student’s t-test was used to compare CS and RS (“***” above horizontal line). This method was applied to similar plots below. **e.** Difference in speed change (represneting slowing) between during- and pre-stimulation peroids for CS. **f.** Licking difference between during- and pre-stimulation peroids for CS. **g.** Difference in speed change (represneting slowing) between during- and pre-stimulation peroids for RS. **h.** Licking difference between during- and pre-stimulation peroids for RS. **i.** Comparison of RBR pre-reward slowing (i.e., speed change), CS vs RS. **j.** Comparison of RBR pre-reward number of licks, CS vs RS. Significant p value for Dur, CS vs RS: 2.72×10^−2^. **k.** Slowing difference (i.e., difference in speed change) in pre-reward zone (orange) versus those in other track zones (gray). **l.** Lick difference in pre-reward zone (orange) versus those in other track zones (gray). **m-x.** Similar to **a-l** but for combined CS vs RS. Significant p values for Post, CS vs RS: **w**: 1.18×10^−2^; **v**: 4.11×10^−3^. For pre-reward CS versus RS: CS data were from seven mice, 21 sessions. RS data were from seven mice, 21 sessions. For combined CS versus RS: CS data were from seven mice, 16 sessions. RS data were from seven mice, 21 sessions. *p ≤ 0.05, **p ≤ 0.01, ***p ≤ 0.001. n.s. p > 0.05. Error bars represent mean ± SEM.

**Figure S17.**
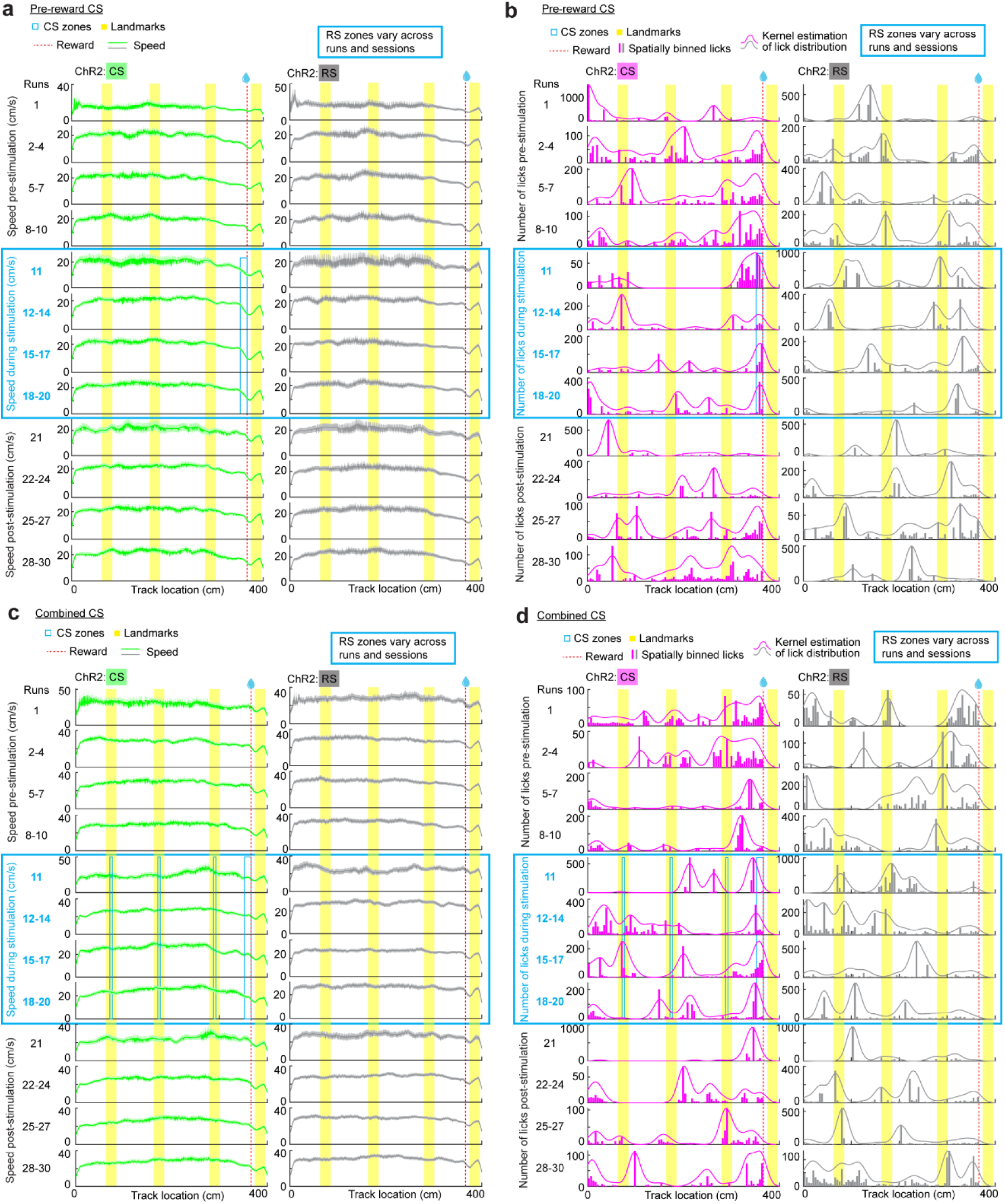
Behavioral effects of pre-reward and combined CS vs RS within runs for ChR2 mice. **a.** For pre-reward CS vs RS: speed in pre-, during- and post-stimulation periods of ChR2 mice. Except the first run in each period that shows the immediate effect of with and without stimulation, the speed was averaged across every three runs. **b.** Similar to **a** but for number of licks. **c.** Similar to **a** but for combined CS vs RS. **d.** Similar to **b** but for combined CS vs RS. For pre-reward CS versus RS: CS data were from nine mice, 25 sessions. RS data were from nine mice, 26 sessions. For combined CS versus RS: CS data were from ten mice, 22 sessions. RS data were from ten mice, 27 sessions.

## Materials and Methods

### Animals

All animal procedures were performed in accordance with animal protocol 1524 approved by the Institutional Animal Care and Use Committee (IACUC) at NIH/NINDS. For two-photon imaging experiments on the 4 m track, 6 GP5.3 mice^20^ (C57BL/6J-Tg (Thy1-GCaMP6f) GP5.3Dkim/J, JAX stock #028280, 1 male, 5 females) were initially used. The mice were 8-10-month-old at the start of imaging. The analyses focused only on the data from 4 mice (1 male, 3 females), which successfully learned the novel environment (good performers), as described in “***Behavioral classification***”. For two-photon imaging experiments on the 875 cm track, 5 GP5.3 mice (3 males, 2 females) were used. The mice were 4-5-months old at the start of imaging. The optogenetics experiments used C57BL/6J mice ranging from 3-6.5 months old at the time of fiber implantation. The electrophysiological recording experiments used 2 ChR2 mice (1 male and 1 female). The non-reward CS/RS experiments used 4 ChR2 mice (4 males) and 3 EGFP mice (1 male and 2 females). The short-non-reward CS used 8 ChR2 mice (5 males and 3 females) and 5 EGFP mice (2 males and 3 females). The pre-reward CS/RS used 9 ChR2 mice (5 males and 4 females) and 7 EGFP mice (3 males and 4 females). The combined CS/RS experiments used 10 ChR2 mice (6 males and 4 females) and 7 EGFP mice (3 males and 4 females). The visual cortex experiments used 5 ChR2 mice (2 males and 3 females) and 3 EGFP mice (1 male and 2 females). Histology experiments used on ChR2 mouse (1 female) and 2 EGFP mice (2 females). One three-month old male mouse was used to demonstrate the expression of ChR2-GFP in the MEC. Mice were maintained on a reverse 12-hr on/off light schedule, with all experiments performed during the light off period.

### Rodent surgeries

#### General procedures for all surgeries

Mice were anesthetized using a tabletop laboratory animal anesthesia system (induction: 3% isoflurane, 1 L/min oxygen, maintenance: 0.5–1.5% isoflurane, 0.7 L/min oxygen, VetEquip, 901806) and the surgery was performed on a stereotaxic alignment system (Kopf Instruments, 1900). A homeothermic pad and monitoring system (Harvard Apparatus, 50-7220 F) was used to maintain a body temperature of 37 °C. After anesthesia induction, dexamethasone (2 mg/kg, VetOne, 13985-533-03) and saline (500 µL, 0.9% NaCl, McKesson, 0409-4888-50) were administered by intraperitoneal (IP) injection, and slow-release buprenorphine (1 mg/kg, ZooPharm, Buprenorphine SR-LAB) was administered subcutaneously. Enroflo× 100 (10 mg/mL, VetOne, 13985-948-10) was used as an antimicrobial wash just after the skull was exposed and just prior to sealing the skull. At the end of surgery, the exposed skull was coated with n-butyl cyanoacrylate tissue adhesive (Vetbond, 3 M, 1469SB). A single-sided steel headplate for head fixation was mounted to the right side of the skull and adhered with dental cement (Metabond, Parkell, S396).

#### Microprism construction

Microprism implants were constructed similarly to those described previously^12^. A cannula (MicroGroup, 304H11XX) was attached to a circular cover glass (3 mm, Warner Instruments, 64-0720). A right angle microprism coated with aluminum on the hypotenuse (1.5 mm, OptoSigma, RBP3-1.5-8-550), was then attached to the opposite cover glass side. All attachments were performed using UV-curing optical adhesive (ThorLabs, NOA81).

#### Microprism implantation surgery

Microprism implantations were performed similarly to those described previously^12^. All insertions were performed on the left hemisphere, aligning with previous observations of more favorable vasculature^43^. A 3 mm craniotomy was centered at 3.4 mm lateral to the midline and

0.75 mm posterior to the center of the transverse sinus (approximately 5.4 mm posterior to the bregma). Mannitol (3 g/kg, Millipore Sigma, 63559) was administered by IP, and a durotomy was then performed over the cerebellum. The microprism was inserted into the transverse sinus and sealed to the skull with Vetbond. The head plate was then mounted on the skull opposite the craniotomy. Finally, the prism and head plate were adhered to the skull with Metabond.

#### Viral injection surgery for MEC stimulation

Mice were bilaterally injected with AAV8-hSyn-ChR2(H134R)-GFP (Addgene: 3.3 × 10¹³ vg/mL; diluted 6 times in mannitol), AAV5-hSyn-ChR2(H134R)-EYFP (Addgene: 2.4 × 10¹³ vg/mL; diluted 3 times in mannitol), or AAV8-hSyn-EGFP (Addgene: 3.0 × 10¹³ vg/mL; diluted 9 times in mannitol). 100 or 200 nl of virus was pressure-injected through a glass micropipette at each injection site at a rate of 100 nl/min. On each hemisphere, mice were injected at 2 sites in the MEC (0.77 mm anterior to the transverse sinus, 3 mm lateral to bregma, 1.79 mm from the surface of the brain; 0.6 mm anterior to the transverse sinus, 3.36 mm lateral to the bregma, 1.42 mm from the surface of the brain) with the head tilted up 18°.

#### Viral injection surgery for visual cortex stimulation

Mice were bilaterally injected with AAV5-Syn-ChR2(H134R)-EYFP (Addgene: 1.8 × 10^13^ vg/mL; diluted 10 times in mannitol), AAV5-hSyn-EGFP (Addgene: 2.26 × 10^13^vg/mL; diluted 10 times in mannitol). 400 nl of virus was pressure-injected through a glass micropipette at each injection site at a rate of 100 nl/min. On each hemisphere, mice were injected at 1 site in the visual cortex (3.2 mm posterior to the bregma, 2.5 mm lateral to bregma, 0.4 mm from the surface of the brain).

#### Fiber implantation for MEC stimulation

At least 3 weeks following the viral injection, mice were chronically implanted bilaterally with Lambda fibers (Plexon) at 0.3 mm anterior to the transverse sinus, 3.2 mm lateral to bregma, and inserted to a depth of 2.5 mm from the brain surface as described previously^12^. Fiber dimensions were as follows: 0.66 NA (numerical aperture), 3.0 mm total length (1.0 mm implant length; 2.0 mm active length; 200/230 µm core fiber). Following the fiber insertion, a thin layer of Vetbond was applied, followed by a thick layer of Metabond to cover the exposed skull.

#### Fiber implantation for visual cortex stimulation

The stimulation for visual cortex used the same fibers for consistency with the experiments in the MEC. The fiber was inserted to a depth of 2mm from the brain surface.

### Virtual reality setup

For all behavioral experiments, a customized virtual reality (VR) setup was used to project a one-dimensional (1D) virtual environment based on the running of a mouse, similar to that described previously^12^. Mice were head-fixed onto an air-supported polystyrene ball (8” diameter, Smoothfoam) using the mounted head plate. The ball rotated on an axle, allowing only forward and backward rotation. The virtual environment was projected onto the inside of a hemispherical dome filling the visual field of the mouse (270° projection). An optical flow sensor (Paialu, paiModule_10210) with infrared LEDs (DigiKey, 365-1056-ND) was used to measure the rotation of the ball and thereby control the motion of the virtual environment. The optical flow sensor output to an Arduino board (Newark, A000062), which transduced the motion signal to the computer controlling the virtual reality. An approximately 4 μl water reward was provided via a lick tube at a fixed location using a solenoid. A lick sensor connected to both the lick tube and head plate holder was used to detect mouse licking. A mouse licking the lick tube created a closed circuit between the lick sensor, the lick tube, the mouse (from the tongue to the skull), the headplate (which directly contacts the skull), and the head plate holder. The solenoid and lick sensor were controlled using a Multifunction I/O DAQ (National Instruments, PCI-6229). The virtual environments were generated and projected using ViRMEn software (Princeton, version 2016-02-12). Imaging and behavior data were synchronized by recording a voltage signal of behavioral parameters from the VR system using the DAQ. ViRMEn environments were updated at 60 Hz. The DAQ input/output rate was 1 kHz. The synchronization voltage signal was updated at 20 kHz. Final behavioral outputs were matched to the imaging frame rate (30 Hz, see below) for synchronization.

Environments were colored blue and projected through a blue wratten filter (Kodak, 53-700) to reduce contamination of the imaging path with projected light. Virtual environments were 1D linear tracks with patterned walls and patterned visual landmarks at fixed locations. At the end of the track, mice were immediately teleported to the start of the track.

### Behavior

#### General training

Mice were allowed to recover for 5 days post-surgery (prism implantation, virus injection, or optogenetic fiber implantation) and were then put on water restriction, receiving 1 mL of water per day. After approximately 3 days of water restriction, mice were trained daily in VR. One session, which lasted around 45 minutes, was conducted on each day.

#### Training of 4 m track imaging mice

For the 4 m track imaging mice, one training session (30∼60 minutes) was conducted per day. 5 out of 6 mice were first trained on training tracks to facilitate running, and all mice were trained in other tracks for prior experiments. After all mice were able to continuously run in VR tracks, they were further trained on a 4 m track until familiarization, as measured by consistent running (>40 trials per hour) and stable significant anticipation of reward locations for 3 days (less than 5% change in averaged predictive licking 1 and predictive slowing within a day, see “***Predictive Licking 1***” and “***Predictive Slowing***” for quantification). The mice were then switched to the 4 m novel environment for spatial learning experiments.

#### Training of 875 cm track imaging mice

For the 875 cm track imaging mice, one training session (30∼60 minutes) was conducted per day. All mice were first trained on a 1 m track to facilitate running and were trained in other tracks for prior experiments. The mice were then switched to the 875 cm novel environment for spatial learning experiments.

#### Training of optogenetic mice

After recovery from viral injection surgery, optogenetic mice were trained for one session (30∼60 minutes) per day. They were first trained on shorter training tracks (1-2 meters) to facilitate running and then were trained directly on the 4 m track used for optogenetic experiments. Fiber implantation was conducted when half of the mice in a cohort (∼10 mice/cohort) reached the criterion for “good performers” (see “***Behavioral classification***”), generally in 5–8 weeks after viral injection. After recovery from fiber implantation surgery, the mice were trained on the same 4 m track. The optogenetics experiments were conducted when the mice resumed continuous running.

#### Predictive licking 1 (previous whole-session method, only used in training and mouse classification)

Predictive licking 1 was calculated per session following the previous method^12^, as the percentage of licks within 20 cm before the reward location across all runs relative to those at other locations across all runs (excluding 30 cm after reward due to reward consumption). This calculation was only used to determine the timing to switch the imaging mice to the novel 4m track (“***Training of imaging mice***”), the timing to conduct fiber implantation for optogenetics mice (“***Training of optogenetic mice****”)*, and whether the imaging mice had successfully learned the track as described in “***Behavioral classification***”. This parameter was not calculated on a run-by-run basis nor used to evaluate the effect of optogenetics.

#### Predictive licking 2 (new per-run method, used in all figures)

To evaluate the effect of optogenetics per run and be better aligned with the calculation of predictive slowing below, predictive licking 2 was calculated on a run-by-run basis. For each run, it was the percentile of number of licks in a “pre-reward zone” among those within other track zones, which had the same length as pre-reward zone and were sampled at a rolling basis with 1-cm interval (excluding 30 cm after reward due to reward consumption). For the runs where the mice did not lick in the pre-reward zone or any track location prior to the reward, the value was set as “invalid”. This could be due to two reasons: (1) the mice were unable to predict reward location; (2) the mice predicted reward location, as indicated by reliable predictive slowing, but did not lick until the reward was delivered. The length of “pre-reward zone” was 20 cm for all stimulation conditions. Predictive licking 2 per session was the average across runs.

Predictive licking 2 was used in all figures and all “***Data analysis***” below.

#### Predictive slowing

Predictive slowing was calculated per run as described previously^12^. In each run, the pre-reward slowing value was calculated as the mean speed changes (Δv) within a 70 cm window pre-reward (i.e., 296–366 cm). If the mice indeed slowed down before the reward, Δv was largely negative (i.e., “slowing”). The rest of the track was analyzed using a rolling average of Δv for the same window size at 1 cm intervals, generating a series of “comparison slowing values”. Predictive slowing for the run was quantified as the percentile of the “pre-reward slowing” among the “comparison slowing values” from the rest of the track, such that a higher percentile signified a greater slowing and thus, better predictive slowing. The rolling averages calculated for the comparison values excluded any window intersecting the edge of the track or areas close to the reward (from 90 cm before to 30 cm after reward delivery) to avoid edge effects and behaviors related to reward consumption, respectively. Predictive slowing of a session was computed as the average slowing across all runs. Predictive slowing per session was used to determine the timing to switch imaging mice to the novel 4m track (“***Training of imaging mice***”), the timing to conduct fiber implantation for optogenetic mice (“***Training of optogenetic mice***”), and whether imaging mice had successfully learned the track (“***Behavioral classification***”). Given its higher reliability compared to predictive licking^12^, run-averaged predictive slowing in pre-stimulation period was used to determine whether a session of pre-reward and combined CS/RS was included for analyses (“***Behavioral classification***”). Per-session and per-run predictive slowing were also used in all figures.

#### Behavioral classification

The analyses of imaging data were focused on the mice that successfully learned the novel environment (“good performers”), as reflected by their high predictive slowing and predictive licking 1 based on previously developed criteria in a similar task^12^. We calculated the averaged predictive slowing (avgPS) and averaged predictive licking 1 (avgPL1) in the last 6 days in the novel environment. If the avgPS and avgPL1 of a mouse were below 65.7^th^ percentile and 22.2%, respectively, the mouse was considered as a “poor performer” that have not learned the environment well, and its data were excluded.

For the optogenetics experiments, fiber implantation was conducted when half of the mice in a cohort reached the criterion for “good performers”, the avgPS or avgPL1 of which were above 65.7^th^ percentile or 22.2%, respectively, for at least two consecutive days.

During optogenetics, the performances were determined on a per-session basis and were only based on predictive slowing, which could represent the prediction of reward location and showed much lower variation than licking^12^. For a session, if its 10-run averaged predictive slowing during pre-stimulation was equal or above the 65.7^th^ percentile, the session was determined as a “good performance session”. Otherwise, it was a “poor performance session”.

Non-reward (15 cm) CS/RS included all sessions without behavioral classification. Short-non-reward (5 cm) CS/RS experiments included good performance sessions. The pre-stimulation reward-predictive slowing and licking in non-reward and short-non-reward sessions were comparable. Pre-reward CS/RS and combined CS/RS experiments focused on “poor performance sessions”, when predictive behaviors in pre-stimulation period were low so that the behaviors could potentially be improved by stimulations.

### Two-photon imaging

Imaging was performed using an Ultima 2Pplus microscope (Bruker) configured with the above VR setup. A tunable laser (Coherent, Chameleon Discovery NX) set to a 920 nm excitation wavelength was used. Laser scanning was performed using a resonant-galvo scanner (Cambridge Technology, CRS8K). GCaMP fluorescence was isolated using a bandpass emission filter (525/25 nm) and detected using GaAsP photomultiplier tubes (Hamamatsu, H10770PB). A 16x water-immersion objective (Nikon, MRP07220) was used with ultrasound transmission gel (Sonigel, refractive index: 1.3359^44^; Mettler Electronics, 1844) as the immersion media.

The anterior-posterior (AP) and the medial-lateral (ML) angle of the prism (i.e., the angle of the surface of the prism along to the AP or ML direction of the mouse) relative to the head-fixed position of the mouse were measured prior to the first imaging session. The head plate holder and rotatable objective angles were set daily to align the objective with the prism in the AP and ML direction, respectively, such that the objective was parallel to the prism surface. Black rubber tubing was wrapped around the objective and imaging window to prevent light leakage into the objective. Microscope control and image acquisition were performed using Prairie View software (Bruker, version 5.5 or 5.7). Raw data was converted to images using the Bruker Image-Block Ripping Utility. For 4 m track imaging experiments, imaging was performed at 30 Hz with 512 × 512 resolution at a single focal plane. For 875 cm track imaging experiments, dual-plane imaging was performed through a Z-Piezo, which enabled the objective to travel between two planes (30-µm apart) within layer 2. Imaging data in each plane were collected at 7.4 Hz and at 512 × 512 resolution. For all imaging experiments, average beam power at the front of the objective was typically 70-115 mW. Imaging and behavior data were synchronized as described above.

### Image processing

Imaging data was down-sampled by a factor of three by taking the average of each consecutive block of 3 frames and was processed as previously described using published MATLAB scripts^12^. Motion correction was performed using cross-correlation based, rigid motion correction. For the 4 m track imaging experiments, identification of regions of interest (ROIs, active cells) with correlated fluorescence changes was performed using principal component analysis combined with independent component analysis^45^. For the 875 cm track imaging experiments, identification of ROIs and the extraction of their fluorescence time course were performed using Suite2p v0.11.158^46^. For all imaging mice, the fluorescence time course of individual ROIs was then extracted. The fractional change in fluorescence with respect to baseline (ΔF/F) was calculated as (F(t) – F0(t)) / F0(t)^43^.

The mean ΔF/F per run for a cell was calculated as a function of location along the track in 5 cm bins. Data points when the mouse was moving below a speed threshold were excluded from this analysis. The speed threshold was calculated by generating a 100-point histogram of all instantaneous velocities greater than 0 and taking the value twice the center of the first bin (approximately 1% of max positive speed).

For the 4 m track imaging experiments, to remove artifactual ROIs occasionally caused by light leak, ROIs with an extreme non-circularity (ROIs were the ratio of the major to minor axis of the ellipse with the same normalized second central moments as the region was greater than 3.3) were removed from further analysis. The first and last runs of an imaging session were removed from further analysis for all imaging experiments.

### Optogenetics

A blue LED module was coupled to the Lambda fibers, and the blue light was used at ∼8–9 mW. Each stimulation session began with 10 no-stimulation runs (pre-stimulation), followed by 10 consecutive stimulation runs (during stimulation), and then switched back to 10 no-stimulation runs (post-stimulation). For each stimulation run, specific locations along the 4 m track (excluding the first 5 cm and the track area between the reward and the end of the track) were selected for stimulation. When the mouse location was within the stimulation zone and the speed was above 1 cm/s, a 10 Hz pulse train (2 ms duration; controlled by Pulse Pal 2, Sanworks) of 465 nm light was delivered. If the mouse was within the stimulation zone and paused (speed <1 cm/s), the pulse train was not delivered until the mouse moved again (speed > 1 cm/s). The connection area between the fiber and optic patch-cord was covered by black aluminum foil (BKF12, Thorlabs) to minimize potential visual distraction caused by blue light leakage. The first five runs of each behavioral session were excluded from behavioral analysis, as some mice required a few runs to warm up. For all CS and RS experiments, each mouse underwent multiple stimulation sessions until we collected at least three CS sessions and three RS sessions. The orders of CS and RS sessions were random.

Non-reward CS was conducted at 66.5 to 81.5 cm (for the landmark centered at 74 cm), 166.5 to 181.5 cm (for the landmark centered at 174 cm), 282.5 to 297.5 cm (for the landmark centered at 290 cm). Short-non-reward CS was conducted at 71.5 to 76.5 cm (for the landmark centered at 74 cm), 171.5 to 176.5 cm (for the landmark centered at 174 cm), 277.5 to 302.5 cm (for the landmark centered at 290 cm). Pre-reward CS was conducted at 351 to 366 cm (the reward was at 366 cm). Combined CS was conducted at 71.5 to 76.5 cm (for the landmark centered at 74 cm), 171.5 to 176.5 cm (for the landmark centered at 174 cm), 287.5 to 292.5 cm (for the landmark centered at 290 cm), and 351 to 366 cm (before reward). The corresponding RS for each CS condition was done by randomizing CS zones along the track on run-by-run and session-by-session bases.

#### Optogenetic stimulation and electrophysiological recording

To determine the optimal light stimulation parameters, 32-channel multi-unit recordings were performed in anesthetized mice injected with AAV8-hSyn-ChR2(H134R)-EGFP or AAV8-hSyn-EGFP, as previously described^12^. A total of 12 stimulation conditions were tested, consisting of four pulse durations (2, 5, 10, and 20 ms) and three frequencies (10, 20, and 40 Hz). Each stimulation was delivered for 1 s with 10-s intervals between stimulation conditions, and conditions were presented in a randomized order.

#### Signal processing and feature extraction

Recorded signals were band-pass filtered into low (0.7-170 Hz)- and high (250-8,000 Hz)-frequency components to assess population-level neuronal activation and spiking activity, respectively. To evaluate response stability across repeated stimuli, the decay of peak amplitudes in the low-frequency traces was quantified by fitting an exponential function of the form *y* = *A* + *Be*^−*kx*^, where the time constant was defined as *τ* = 1/*k*. x was time and y was voltage.

Spikes were detected from high-frequency signals using a threshold of mean + 3 standard deviations (STDs). Spikes occurring within the photoelectric artifact exclusion time window (defined below) were excluded. In addition, spikes within 5 ms prior to stimulation were excluded to avoid high-frequency filtering artifact arising from large photoelectric deflections after stimulation.

To exclude photoelectric artifacts, an exclusion window was determined from recordings in control (EGFP-only) mice. The exclusion time was defined as the time when the firing rate decayed to the threshold, which captured 90% of 1-ms binned baseline activity events (30ms after stimulation and 5 ms before the next stimulation), thereby minimizing exclusion of bona fide neural responses. This threshold was mean + 4.2 STDs in the average PSTH across 32 channels, resulting in an exclusion window of 7.5 ms after stimulation onset. Under this condition, the stimulation artifacts were clearly removed in EGFP with no significant difference among pre-, during (early, intermediate, late)-, post- stimulation periods. Pre-stimulation period was 1-s duration prior to the stimulation start; post-stimulation period was 1-s duration following 1-s stabilization time after the last stimulation.

### Histology

To show the expression of EGFP or ChR2-GFP in the MEC after virus injection, mice were injected with EGFP or ChR2-GFP. Three weeks after the injection, mice were anesthetized with a ketamine (200 mg/kg, VetOne, 13985-584-10) and xylazine (20 mg/kg, VetOne. 13985-612-50) cocktail and were transcardially perfused with 4% paraformaldehyde (PFA, Electron Microscopy Sciences,15713) in phosphate buffer solution (PBS, Corning, 46-013-CM). Brain tissues were dissected and fixed in 4% PFA in PBS overnight at 4 °C. Sagittal slices (40 µm thick) were prepared using a VT1200S vibratome (Leica Biosystems). The image of the slice was taken under a Leica epifluorescence microscope (Leica 165FC).

#### Immunohistochemistry

Slices were washed in PBS (3 × 10 min), and then blocked with 10% bovine serum albumin (BSA, Millipore Sigma, A3294), 0.5% triton X-100 (Millipore Sigma, T9284), and PBS for 1 h at room temperature. Primary and secondary antibodies were diluted in 2% BSA, 0.4% triton, and PBS. Slices were incubated in diluted primary antibodies for 48-72 h at 4 °C, then washed in PBS (3 × 10 min). Slices were incubated in diluted secondary antibodies (1:500 dilution) for 1.5–2 h at room temperature, then washed in PBS (3 × 10 min) and mounted with mounting medium (Vector Laboratories, H-1000-10). Images were collected using a Zeiss 880 spectra confocal. Primary antibodies used include mouse monoclonal IgG1 anti-reelin (1:2000, Abcam, AB78540), rabbit monoclonal anti-calbindin (1:1000, Abcam, AB108404), mouse monoclonal IgG1 anti-parvalbumin (1:1000, Millipore Sigma, P3088), and rabbit monoclonal anti-somatostatin 28 (1:250-500, Abcam, AB111912). Secondary antibodies included Alexa 568 conjugated goat anti-rabbit antibody (Invitrogen, A11036) and Alexa 647 conjugated goat anti-mouse IgG1 antibody (Invitrogen, A21240).

Cells positive for EGFP, ChR2-GFP, reelin, calbindin, parvalbumin (PV), and somatostatin (SST) were manually selected for 3 focal planes per slice. Overlaps between cell-type markers and viral expression markers were manually determined as the percentage cells positive for the viral expression marker that were also positive for the cell-type marker.

### Data analysis

#### Run-by-run consistency of calcium activity along the track

Run-by-run (RBR) activity consistency was calculated per cell within a rolling window of 5 spatial bins (5 cm per bin) along the track. For a 400 cm track with 80 spatial bins, the correlations were calculated for areas within bins 1 to 5, 2 to 6, 3 to 7, … , and 76 to 80. Within each window, the spatially binned mean ΔF/F per run was correlated with that of every remaining runs. The average of these correlations was the run-by-run consistency value for this window.

For the 4 m track imaging experiments, the activity before reward included bins 67 to 73. The activity at landmarks included bins 13 to 17, 33 to 37, and 55 to 60. The other activity, which was outside landmarks and the before-reward area, included bins 1 to 12, 18 to 32, 38 to 54, and 61 to 66. The bins between the reward and the end of the track (74 to 76) were removed due to potential effect of reward consumption. All bin numbers here refer to the bin numbers after taking the rolling average.

For the 875 cm track imaging experiments, the reward was delivered at bin 63. Pre-reward activity consistency included bins 55 to 61, and post-reward activity consistency included bins 65 to 70. All bin numbers here refer to the bin numbers after taking the rolling average.

RBR activity consistency was not used as an exclusion criterion for mice, sessions, or neurons. To ensure complete coverage of all track locations, the first run, which typically had incomplete track coverage due to delayed imaging initiation relative to the onset of VR navigation, was excluded from each session. Analyses were then restricted to the subsequent 15 runs to enable comparable comparisons across sessions and FOVs.

#### 95% day for behaviors and RBR activity consistencies across learning

To determine the number of days when the reward-predictive slowing, licking, and RBR activity consistencies at different track zones reached 95% of their plateaus (95% day), their averaged curves (across mice) were fitted with a one phase association exponential function (Prism 9.3.1, GraphPad), where the y axis was the behavior or activity consistency, and the x axis was days. The start points of fitted curves were set to be “undetermined”. Once the values for the plateau and K, a rate constant, were derived, 95% day was calculated as ln20/K.

#### Run-by-run population coherence

Population coherence was measured using population vector correlations (PVC), as described previously^47^. For every track location (5 cm bin with no rolling window) on every run, a population vector was calculated as spatially binned ΔF/F for all cells (N cells x 1, each cell was first normalized by peak activity level across track locations). These population vectors were correlated across both track location and run. To generate a run-by-run measure of population coherence, vector correlations were averaged across run comparisons, providing a PVC matrix of track location versus track location as in **Fig. S3A**. On-diagonal PVC was defined as the exact diagonal of each matrix (i.e. the correlation between population vectors at identical correlation, but different runs). For on/off diagonal contrast, on-diagonal was the average of PVC correlations with a spatial bins distance between 0 and 2, and off-diagonal was the average of PVC correlations with a spatial bin distance greater than 6. The beginning and end of the rack were considered connected for the bin distance calculation. The on/off diagonal contrast was the difference between the on-diagonal and off-diagonal averages.

### General linear mixed models

#### Spatial behavioral distributions

For mutual information and general linear mixed models, spatial behavioral distributions were calculated for mouse velocity, acceleration, lick probability, and reward distance, for every track location (5 cm bin with no rolling window). Prior to velocity calculation, time and mouse location were smoothed with a 2-element and 5-element rolling average, respectively, to reduce noise. Velocity for a given spatial bin was the average velocity of all time-points the mouse spent in that spatial bin within a run. Acceleration was calculated per-run based on the bin-averaged velocity and time spent in each spatial bin. The final velocity and acceleration distributions were the average of the per-run distributions. The lick probability distribution was the percentage of runs that the mouse licked with a given spatial bin. Reward distance was the distance in cm from the reward location to a given spatial bin. For all these calculations all times after reward delivery were excluded, so only reward distance was always positive.

#### Mutual Information

For each mouse session, Shannon mutual information^48,49^ (MI) was calculated pre-reward (individual spatial bins, 1-73) between spatial RBR consistency (R below) and behavioral distribution (velocity or lick probability, B below) as follows:

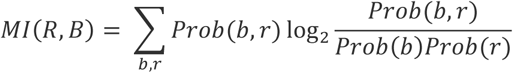

where Prob(b,r) was the joint probability of observing a given spatial RBR activity consistency level and behavioral variable level (velocity or lick probability), Prob(b) was the marginal probability of behavioral variable level, and Prob(r) was the marginal probability of spatial RBR activity consistency level. This calculation was performed per day with each pre-reward spatial bin used as a datapoint. Each variable was discretized into uniform bins between the minimum and maximum value of that variable. For **Figures S3j-l**, both RBR consistency and the behavioral variables were discretized into 6 uniform bins. Because MI can be sensitive to the choice of bin number, MI was calculated for additional bin number combinations (2-10 for each variable) for **Figures S3m**. Shuffled MI was calculated identically after randomly permuting the behavioral distribution across track locations. Plotted MI distributions show kernel smoothing (KS) density distribution across mouse session constrained between 0 and 1. Real distributions were compared with 10,000 shuffled MI distributions with a permutation test (i.e. p = the percentage of shuffle distribution averages greater than or equal to the real MI distribution average).

#### General linear mixed model fitting

General linear mixed model (GLMM) analysis^50,51^ was used to predict behavioral variables (velocity) as a function of activity variables (RBR consistency) and other behavioral variables (acceleration, lick probability, distance to reward) as fixed-effect predictors with mouse ID, day number, and their interaction as random effects. RBR consistency was averaged across FOV within each mouse. Individual pre-reward spatial bins (1-73) from every mouse and day were used as observations. GLMMs were fit to predict velocity (with all other variables) with 5-fold cross-validation across all observations using the MATLAB fitglme function. Velocity prediction used a normal response distribution. All predictor variables were standardized using z-score normalization. Predictor coefficients (β), p-values, and variance explained contribution (See “Dominance Analysis”) were the average values across the 5-fold cross validation. Predictor coefficients (β) and fixed-effect p-values (one-sample Student’s t-test) were the direct outputs of the fitglme function based on the training data. Random effect p-values were determined using a likelihood ratio test.

#### Dominance Analysis

To evaluate the relative contribution of each significant predictor to behavioral variance explained, dominance analysis was performed^51,52^. We determined the effect of removing each fixed-effect predictor from a model with all significant predictors and from all possible sub-models to determine the average contribution of each fixed-effect predictor. The full model and every possible sub-model were trained using the MATLAB fitglme function, as above. The full random effects structure was left in place for all sub-models. Random effects contributions were independently calculated using the individual random-effects variance, total fixed-effects variance, and residual effects variance^53^. This method produced relative variance explained contributions for each fixed- and random-effects predictor, while preserving the total contributions of fixed-effects and random-effects within the full model. Because the sum of the variance explained contributions across all predictors is roughly equal to variance explained by the full model^52^, the individual contributions can be thought of as the relative contribution to the final variance explained.

#### Pairwise population synchrony

Pairwise population synchrony (PPS) was calculated as the mean pairwise correlation between every pair of cells as a function of spatial location. Within each rolling window of 5 spatial bins (5 cm per bin) along the track, spatially binned ΔF/F for each was concatenated across runs for each cell, generating an activity matrix where each column represents the activity of one cell and each row represents the activity at each bin/run (i.e. # rows = 5 bins x # runs). The activity vector for every cell was then correlated with that in every other cell, and the average correlation across all cell pairs is the PPS for that rolling window location. Full track PPS was calculated as the average PPS of all spatial windows, while sub-region PPS was defined as the average PPS within the sub-region spatial windows defined above.

### Classification of grid, cue, and other cells

#### Significant transient detection

ΔF/F including only the significant calcium transients (ΔF/F_sig) was used for cell type classification to improve classification accuracy. For each cell, significant calcium transients were identified using amplitude and duration thresholds, such that the false-positive rate of significant transient identification was 1%^54^.

#### Detection of in-field and out-of-field periods using ΔF/F_sig traces

To calculate in-field period, regions of the track with significantly consistent activity, the mean ΔF/F_sig (averaged ΔF/F_sig within every 5 cm nonoverlapping spatial bin along the track) was compared to shuffles of the original ΔF/F_sig as described previously^21^. Each shuffle was calculated such that the original spatial location of each time point was preserved, but the ΔF/F_sig was shuffled by bisecting the full ΔF/F_sig time course at a random time point (between 5% and 95% of the whole time course) and swapping the order of the resulting halves. The mean ΔF/F_sig of the shuffle was then calculated as described above. A in-field period was defined as a region of at least 3 consecutive 5 cm bins (except that the fields at the beginning and end of the track could have 2 bins) that had a mean ΔF/F_sig higher than 80% of 1000 shuffles at the corresponding bins. Additionally, at least 20% of runs were required to have a significant calcium transient in the spatial field. An out-of-field was defined as two or more bins that had a mean ΔF/F_sig below 25% of 1000 shuffles. Bins with intermediate mean ΔF/F_sig remained unassigned.

#### Classification of grid cells

Grid cells were identified as described previously based on the features of in-field and out-of-field periods^21,24^: (a) Grid cells need to have at least two in-field periods. (b) Each grid cell must have more than L/(5w) transitions between in-field and out-of-field periods, where L represents track length and w represents mean field width (track length of in-field period) of the grid cell’s response. (c) The widest field width of the cell must be smaller than 5w. (d) At least 30% of the bins must be assigned to either in-field or out-of-field periods. (e) The ratio between in-field and out-of-field mean ΔF/F_sig must be greater than 2. All cells that met the grid cell criteria in a given imaging session, except those defined as cue cells (see below), were defined as grid cells.

#### Classification of cue cells

Cue scores were calculated as previously described^55^ using ΔF/F_sig traces. The mean ΔF/F_sig of the cell was first shifted to best match the environment cue template, which contained ones and zeros representing track areas with and without landmarks, respectively. A landmark zone was identified for each landmark by including the landmark itself and the surrounding region expanded by half of the landmark width on both sides. The correlations between shifted activity and cue template within individual landmark zones were calculated and were further averaged as cue score of the cell. 200 shuffled cue scores were generated for each cell using the same calculation but by randomizing the landmark location in the cue template. Shuffled cue scores from all cells across all FOVs and imaging days were pooled to generate an overall cue score distribution. A cell in a given imaging session was identified as a cue cell if its cue score was above the 95^th^ percentile of the shuffle cue score distribution.

#### Classification of other cells

In a given imaging session, all cells not identified as cue cells or grid cells were defined as other cells.

### Behavioral session criteria for optogenetics

Across the four experiments (non-reward, short non-reward, pre-reward, and combined stimulations), any session meeting either exclusion criterion was excluded from all analyses.

#### Adjusting the baselines (pre-stimulation)

We observed that in some optogenetics sessions, the percentiles of RBR predictive slowing in the pre-stimulation period, which served as baselines for the optogenetic stimulation, gradually increased or decreased, potentially caused by the slow acclimation of the mice to the task. These slowly evolving baseline trends could confound the behavioral effect of optogenetic stimulation. To solve this issue, we removed the sessions with unstable predictive slowing percentile in the pre-stimulation period. For each stimulation condition, we first calculated the averaged percentile of RBR predictive slowing in the pre-stimulation period (10 runs) across all sessions (AvgPS_AllSessions_). We then calculated the fractional difference (D) between the AvgPS_AllSessions_ in runs 1-7 (AvgPS_AllSessions_1_7) and runs 8-10 (AvgPS_AllSessions_8_10) for all sessions as follows:

D_AllSessions_ = (AvgPS_AllSessions_8_10 – AvgPS_AllSessions_1_7) / AvgPS_AllSessions_1_7.

We also similarly calculated the fractional difference between the averaged predictive slowing of runs 1-7 (PS_OneSession_1_7) and runs 8-10 (PS_OneSession_8_10) for each session as follows:

D_OneSession_ = (PS_OneSession_8_10 – PS_OneSession_1_7) / PS_OneSession_1_7.

If D_AllSessions_ was above 0 and below 0, we removed the individual sessions, the D_OneSession_ of which was above and below 0.5 and -0.5, respectively. If none of the sessions met these criteria, no session was removed.

After the above adjustments, several sessions in combined CS and RS were removed: 7 ChR2 CS sessions, 1 ChR2 RS session, 5 EGFP CS sessions.

Due to the high variability of predictive licking (i.e., predictive licking 2) across runs within the same session, similar adjustments led to the removal of more than half of the data in some stimulation conditions. Therefore, no baseline adjustment was conducted based on predictive licking. However, one adjustment was made for ChR2 mice with non-reward CS. The four mice in this group initially had 16 sessions, but 6 sessions had 100^th^ percentile of predictive licking 2 in pre-stimulation, leading to significantly different levels of CS and RS in this period. We removed these 6 sessions to ensure that the comparisons between CS and RS were made on sessions with comparable baseline levels.

#### Removing lick sessions with detection artifact

We observed that mice occasionally touched the lick tube using their paws during grooming or held the lick tube, causing a large amplitude licking signal that did reflect true lick signal. To avoid such signals, we removed the session if its licking signal in one 5 cm spatial bins in pre- and during-stimulation periods was greater than 99.98^th^ percentile across all bins in these periods combining all ChR2 and EGFP sessions. This led to the removal of 2 ChR2 sessions for non-reward CS, 3 ChR2 sessions for non-reward RS, 1 ChR2 session for short-non-reward CS, 1 ChR2 session for pre-reward CS, 1 ChR2 session for pre-reward RS, 1 ChR2 session for pre-reward RS, 2 ChR2 sessions for combined CS, and 2 ChR2 sessions for combined RS.

#### Predictive behaviors in pre-, during-, and post-stimulation, per session

The slowing or licking percentiles in the three periods were calculated by averaging the values across the 10 runs in individual periods.

#### Predictive behaviors in pre-, during-, and post-stimulation, per run

Run-by-run predictive slowing percentiles per session were normalized by subtracting the session’s mean pre-stimulation value. Due to the presence of invalid predictive licking in certain runs and sessions in pre-stimulation period, predictive licking percentiles per session were normalized by subtracting mean value across all sessions, rather than per session, in pre-stimulation period.

#### Slowing percentile around stimulation zones

This calculation was only made for non-reward and short-non-reward CS and RS. For each run, “stimulation slowing” was calculated by grouping individual slowing values (negative speed change) between adjacent ViRMEn-sampled speeds points in all three CS zone and took the mean. “Other slowing” was calculated by randomly picking 3 zones with the same length of CS zones in the remaining track locations, excluding the zone form reward to the end of the track. 1000 “other slowing” values were calculated. Slowing around stimulation zones was the percentile of “stimulation slowing” among the 1000 values.

#### Licking percentile around stimulation zones

This calculation was only made for non-reward and short non-reward CS and RS. For each run, “stimulation licking” was the mean number of licks within the three CS zones, each of which were expanded 6 cm to both sides. “Other licking” was calculated by randomly picking 3 zones with the same length of expanded CS zones in the remaining track locations, excluding the zone form reward to the end of the track. 1000 “other licking” values were calculated. Licking around stimulation zones was the percentile of “stimulation licking” among the 1000 values.

#### Slowing and licking percentiles around stimulation zones in pre-, during-, and post-stimulation, per session

The percentiles in the three periods were calculated by averaging the values across the 10 runs in each period within the session.

#### Slowing and licking percentiles around stimulation zones in pre-, during-, and post-stimulation, per run

Run-by-run slowing percentiles per session were normalized by subtracting the session’s mean pre-stimulation value. Due to the presence of invalid licking in certain runs and sessions in pre-stimulation period, percentiles per session were normalized by subtracting mean value across all sessions, rather than per session, in pre-stimulation period.

#### Speed along the track

The speed was calculated between adjacent ViRMEn data points (sampled at 60Hz), averaged within individual 1cm non-overlapping track bins, and smoothed per run by a 15-point gaussian window. The speeds across all sessions were further averaged per run (for the first run in pre-, during-, and post-stimulation period) or per three runs (the remaining runs).

#### Slowing (i.e., speed change) and slowing difference along the track for during- versus pre-stimulation periods

This analysis was for visualizing the slowing along the track in the two periods. The speed (per 1cm track bin) per run in each session was smoothed by moving averaging every N bins (N=20 for pre-reward CS/RS, N=50 for combined CS/RS), and the slowing was calculated per run as the speed change in adjacent bins (i.e. speed in one bin minus speed in the previous bin) in the track zones from 11 to 366 cm. The data in the first 10 cm were removed due to the artifact of teleportation. The speed change was further averaged per run as per-session speed change. The slowing difference between during- and pre-stimulation periods was calculated by subtracting the slowing per session values in pre-stimulation period from those in during-stimulation period.

#### Run-by-run number of licks along the track

The licks that occurred before reward delivery in each run from the beginning of the track to the reward location (366 cm) were first binned into 80 spatial bins. The number of licks per bin across all sessions were averaged per run (for the first run in pre-, during-, and post-stimulation period) or per three runs (the remaining runs). MATLAB function “ksdensity” was used to estimate the distribution of licks along the track.

#### Number of licks and lick difference along the track for during- versus pre-stimulation periods

For each period of each session, the total number licks occurred before reward delivery was summed across all 10 runs and binned into 80 spatial bins from the beginning of the track to 366 cm (reward location). The lick difference was calculated per session by subtracting the number of licks per bin in pre-stimulation from that of during-stimulation. The lick difference per session was further averaged.

#### Slowing before reward

The speed (per 1cm track bin) per run in each session was smoothed by moving averaging every 8 bins, and the slowing was calculated as the speed difference in adjacent bins within pre-reward zone. The slowing of the run was the averaged slowing within the zone. To best capture the speed change in each condition, pre-reward zone was 14 cm before reward for pre-reward CS/RS, and 22 cm before reward for combined reward CS/RS.

To compare the slowing between CS and RS, the slowing in each session/run was normalized by subtracting the mean value of pre-stimulation of the session.

#### Licking before reward

Due to the unreliable licking behavior, the number of licks was calculated by rolling averaging every 2 runs within a period. The total number of licks within pre-reward zone was calculated. To best capture the licks in each condition, pre-reward zone was 12 cm before reward for pre-reward CS/RS, and 27 cm before reward for combined reward CS/RS.

To compare the licking between CS and RS during-stimulation periods, the numbers of licks in each session/run was normalized by subtracting the mean value of pre-stimulation across all sessions.

#### Correlation between CS pattern and slowing pattern

The slowing along the track was first calculated per run per 1cm as in “***Slowing before reward***” but for the track area from 16 to 400 cm (“slowing pattern”), excluding the first 15 cm due to artifact and residual reward licking. The “stimulation pattern” was similarly created for the same track region in 1-cm spatial bins, with within-stimulation zone bins set to -1 and non-stimulation zone bins set to 0, to match the slowing pattern’s negative values indicating speed decreases. Within-stimulation zone was expanded 3 cm around the real stimulation zone in forward and backward directions to adapt to the variation in slowing behavior. Pearson correlation between “slowing pattern” per session and “stimulation pattern” was calculated.

#### Correlation between CS pattern and licking pattern

For a session within a period (pre-, during-, or post-stimulation), this calculation used licks along the track (before reward delivery) in each run. The distribution of licks per run (licking pattern) was estimated using MATLAB function “ksdensity” in 1cm estimation interval within the track region of 16-400 cm, excluding the first 15 cm due to artifact and residual reward licking. The “stimulation pattern” was similarly created for the same track region in 1-cm spatial bins, with within-stimulation zone bins set to 1 and non-stimulation zone bins set to 0, to match the licking pattern’s positive values indicating licking increase. To compromise the variation in lick location, each stimulation zone was expanded in backward and forward track directions. For non-reward and short-non-reward CS/RS, backward expansion was 12 cm, and forward expansion was 25 cm. Pearson correlation between “licking pattern” per session and “stimulation pattern” was calculated.

#### Sub-sampling RS to RS2

This procedure was conducted for non-reward RS sessions of ChR2 mice. There were originally 360 RSs. Only RSs with comparable stim-distances (distance between the current stimulation and adjacent ones, calculated based on stimulation centers) as CSs were included in RS2. The first stimulation in each run should be at least 72 cm away from the track start (as the first stimulation in CS) and 99 cm away from the second stimulation. The second stimulation should be at least 99 cm away from the first and third stimulations. The third stimulation should be at least 99 away from the second stimulation track end. This approach left 61 stimulations in RS2.

#### Aligning RS locations

To evaluate the effect of individual RSs, track locations in before, within, and after stimulation zones were aligned by the start location of “within stimulation zone”. Thus, the speed and number of licks within the three zones were calculated across multiple stimulations, runs and sessions.

#### Down-sampling CS to match the number of RS2

Each down-sampled CS dataset contained randomly picked CS matching the same number of stimulations as in RS2. 1000 datasets were generated. The speed and lick numbers were independently calculated for each dataset.

#### Comparison of speed and licking in different zones across aligned stimulations

Speed of individual runs was averaged in 1 cm spatial bins. For each stimulation of each session, the mean speed in before-stimulation zone (10-50cm before stimulation onset), preceding zone (10cm immediately preceding the stimulation onset), stimulation zone (15 cm), following zone (10cm immediately following the stimulation offset), and after-stimulation zone (10-50cm after reward offset) were grouped per zone. The comparison between the speed in before-stimulation zone and other zones was conducted by paired comparison so that the speeds of the zones around the same run were compared.

The zones for lick number comparison were similarly defined, except that the preceding and following zone were expended by 10cm to adapt to the wide distribution of licks. Accordingly, the before-stimulation and after-stimulation zones were shrunk by 10cm. For each stimulation of each session, the total numbers of licks in the five zones were calculated and compared pairwise.

Similar comparisons were made for 1000 down-sampled CS datasets.

#### Comparing slowing differences in reward zone and other track zones

The calculation used the difference in speed change between pre- and during-stimulation period, grouped in 1-cm bins from 0 to 366 cm and averaged across sessions (“***Slowing and slowing difference along the track for during- versus pre-stimulation periods****”*). The “reward zone” comprised the 23 bins before the reward (totaling 23 cm), defined based on predictive slowing and lick locations of good-performing mice in **Figs S11** and **S12**. The difference in speed change (a negative value indicate slowing) was the mean difference across these bins. The difference was calculated between the pre- and during-stimulation periods per session and further averaged into one value. Similar calculations were performed for other track zones of the same length, selected on a rolling basis (track location 1-5 cm, 2-6 cm, 3-7 cm, etc.). We then calculated the percentage of track zones with difference in speed change above that in the reward zone, which reflected their lower degree of slowing difference than the reward zone.

#### Comparing lick differences in reward zone and other track zones

The calculation used the lick difference between pre- and during-stimulation period, grouped in 80 non-overlapping bins (4.58 cm per bin) from 0 to 366 cm and averaged across sessions (“***Number of licks and lick difference along the track for during- versus pre-stimulation periods****”*). The “reward zone” comprised the 5 bins before the reward (totaling 22.9 cm, comparable to the 23 cm window for slowing difference calculation: ***Comparing slowing differences in reward zone and other track zones***), defined based on predictive slowing and lick locations of good-performing mice in **Figs S11** and **S12**. Lick difference was the mean lick difference across these 5 bins. The difference was calculated between the pre- and during-stimulation periods per session and further averaged into one value. Similar calculations were performed for other track zones of the same length, selected on a rolling basis (bins 1-5, 2-6, 3-7, etc.). We then calculated the percentage of track zones with lick differences below that in the reward zone, which reflected their lower degree of licking increase than the reward zone.

##### General data analysis and statistics

Image processing was performed using published MATLAB (MathWorks, version R2015aSP1) codes as described previously^12^. Data analysis was performed using ImageJ (Fiji, version 1.53q), MATLAB (MathWorks, versions R2015aSP1 and 2020a), and GraphPad Prism (version 9.3.1). Linear correlations and the corresponding *r* and *p* values were calculated using a two-tailed Pearson’s linear correlation coefficient. Mutual information distributions were compared to shuffles using a permutation test. For GLMM analysis, fixed-effect statistics were calculated using one-sample Student’s *t*-test and random-effect statistics were calculated using a likelihood ratio test. The comparison between every two periods (e.g., pre-, versus during-stimulation) were performed using Student’s paired *t*-test and *p* values were adjusted for multiple comparisons using the Bonferroni–Holm method. The comparison between run values in CS or RS in pre- versus during-stimulation condition was evaluated by Student’s t-test. The comparison between CS and RS slowing on a run-by-run basis was evaluated by two-way ANOVA. Due to the presence of invalid licking values, all comparison between CS and RS licking on a run-by-run basis was evaluated by Student’s t-test on all numbers combined across runs and sessions. The remaining comparisons were evaluated by Student’s *t*-test*. P* values less than 0.05 were considered significant (* ≤ 0.05, ** ≤ 0.01, *** ≤ 0.001). All figures show mean and standard error.

## Data availability

Processed data underlying all figures are available in the Source Data file. Further information and requests for resources and reagents should be directed to the lead contact, Dr. Yi Gu.

## Code availability

Custom code used for the analysis of data is available on GitHub at https://github.com/GuLab-NIH/Ma2025.

